# Dysregulated SETX activity at R-loops links transcriptional and splicing alterations to compromised neuronal identity in ALS4

**DOI:** 10.64898/2026.06.30.735532

**Authors:** Marta Giannini, Thierry Gostan, Amal Zine El Aabidine, Camille Bellières, Stephane Nedelec, Odil Porrua

**Affiliations:** Institut de Génétique Moléculaire de Montpellier, Univ Montpellier, CNRS, Montpellier, France; BCM, Univ Montpellier, CNRS, Inserm, Montpellier, France; Institut du Fer à Moulin, 75005 Paris, France; Université Paris Cité, CNRS, Inserm, Institut Jacques Monod, Paris, France

## Abstract

RNA dysregulation is an increasingly recognized driver of neurodegenerative diseases, yet the mechanisms linking defects in RNA metabolism to selective neuronal vulnerability remain poorly understood. Alterations in R-loop homeostasis have been observed in neurodegeneration, but it is unclear how these contribute to neuronal dysfunction.

To address these questions, we investigate ALS4, a juvenile form of amyotrophic lateral sclerosis that is caused by mutations in the helicase SETX, which functions in R-loops resolution and transcription termination. Using isogenic human induced pluripotent stem cell-derived motor neurons (MNs), we show that the ALS4-associated SETX L389S mutation induces axonal defects and widespread transcriptomic alterations, including reduced expression or altered splicing of transcripts critical for neuronal function. ALS4 MNs exhibit a transcriptional signature of cellular stress, aberrant cell cycle re-entry, and compromised neuronal identity, features that are partially shared across familial and sporadic ALS. Mechanistically, these defects are driven by aberrant activation of the TGF-β signaling pathway, as its pharmacological inhibition ameliorates axonal phenotypes. Finally, our analyses support a link between aberrant SETX activity at R-loops and alterations in transcription and splicing.

Altogether, our work reveals how perturbations at the interface of R-loop metabolism, transcription and RNA processing can reshape neuronal identity and drive disease.

## INTRODUCTION

RNA metabolism and transcriptional regulation are central determinants of neuronal homeostasis, and their dysregulation is increasingly recognized as a major driver of neurodegeneration (*1*). Mutations in genes encoding RNA-binding proteins, RNA-processing factors, and transcriptional regulators have been identified in a growing number of neurological disorders, including frontotemporal dementia, spinocerebellar ataxias, spinal muscular atrophy, and amyotrophic lateral sclerosis (ALS). Perturbations of RNA processing, including splicing, mRNA stability, transport, and surveillance, are thought to contribute to widespread transcriptional dysregulation and cellular dysfunction (*2*). In parallel, genome instability and DNA damage have emerged as recurrent features of neurodegeneration (*3*). Increasing evidence suggests that conflicts between transcription and genome maintenance processes may provide a link between these pathological hallmarks (*4*). However, the molecular mechanisms through which defects in RNA metabolism lead to the selective vulnerability of distinct neuronal populations remain poorly understood.

The study of mutations in Senataxin (SETX) provides a unique opportunity to address this question. SETX is an RNA/DNA helicase that plays important roles in transcription termination, genome stability and the resolution of R-loop structures (*5*). R-loops form during transcription when the nascent RNA abnormally reanneals with the template DNA displacing the non-template strand. While these structures partake in the regulation of gene expression and other physiological processes, their excessive accumulation or aberrant persistence can promote DNA damage and genome instability (*6*, *7*). Mutations in the *SETX* gene provoke two clinically distinct neurodegenerative diseases. Recessive loss-of-function mutations in *SETX* cause ataxia with oculomotor apraxia type 2 (AOA2), which selectively affects cerebellar neurons (*8*), while dominant gain-of-function mutations provoke a rare juvenile form of ALS (ALS4), a motor neuron (MN) disease (*9*).

ALS is characterized by the degeneration of MNs in the brain and spinal cord, leading to progressive paralysis of almost all skeletal muscles and death of patients with 5 years of symptoms onset. Roughly 10% of ALS cases are familial, most often caused by dominant mutations in about 30 genes (*10*, *11*). Although genetically heterogeneous, ALS is increasingly viewed as a disorder of RNA metabolism. Indeed, approximately half of the genes mutated in ALS encode either proteins with established roles in RNA metabolism, such as TDP43, FUS and SETX, or proteins that, when mutated, interfere with RNA regulatory pathways, such as C9ORF72 and SOD1 (*1*). The same genes are often mutated in sporadic ALS patients and, indeed, familial and sporadic ALS share clinical symptoms and cytopathological features. For instance, cytoplasmic aggregates of the RNA-binding protein TDP43 are a hallmark of almost all forms of ALS. In addition, DNA damage and genome instability are commonly observed in ALS patient tissues and disease models, further supporting a functional connection between defective RNA metabolism and impaired genome maintenance (*3*).

Consistent with this idea, alterations in R-loop homeostasis have been reported following mutation or depletion of several ALS-associated proteins, including C9ORF72 (*12*), TDP-43 (*13*, *14*), FUS (*15*) and SETX (*16*, *17*). However, it remains unclear whether and how deregulated R-loop metabolism could contribute to ALS pathology. In the context of ALS4, two groups reported reduced R-loop levels in fibroblasts from ALS4 patients compared to healthy individuals (*16*, *17*). In one of these studies, R-loop changes near promoter regions correlated with concomitant differences in the DNA methylation status and the expression of nearby genes (*16*), suggesting a model by which aberrant R-loop resolution by mutant SETX at gene promoters could lead to transcriptional dysregulation via epigenetic changes in ALS4. However, interpretation of these findings is complicated by the comparison of genetically distinct individuals, which limits causal inference, and by the use of fibroblasts, a cell type that differs markedly from the MNs selectively vulnerable in ALS4.

To overcome these limitations, in the present study we generate and characterize an isogenic human *in vitro* model of ALS4 by introducing the highly penetrant L389S *SETX* mutation into human induced pluripotent stem cells (iPSCs) that are differentiated into MNs. Using this system, we investigate how this ALS4-associated SETX mutation alters gene expression programs and MNs homeostasis. We show that ALS4 MNs exhibit axon shortening, accompanied by substantial transcriptome alterations including decreased expression and/or altered splicing of transcripts important for MN functions. Moreover, ALS4 MNs display transcriptional signatures reflecting cellular stress, aberrant cell cycle reactivation, and compromised neuronal identity, that are partially shared by more frequent forms of ALS. Our analyses identify downstream aberrant activation of transforming growth factor-β (TGF-β) superfamily signaling as a major contributor to these molecular and cellular phenotypes. Importantly, pharmacological inhibition of this pathway not only substantially alleviates gene expression deregulation associated with compromised neuronal identity, but also rescues axonal defects, providing a functional link between altered neuronal identity and MN impairment in ALS4. Finally, our findings challenge the current model linking R-loop perturbations to transcriptional deregulation in ALS4 and instead uncover new links between L389S SETX aberrant activity at R-loops and alterations in RNA expression and splicing.

Altogether, our findings provide major advances in the understanding of the molecular pathways linking RNA dysregulation to MN impairment in ALS4 and reveal a potential therapeutic strategy to treat ALS4 and possibly other ALS types.

## RESULTS

### The ALS4 mutation L389S provokes axonal defects at iPSC-derived MNs

To investigate the effects of ALS4 mutations in human motor neurons (MNs), we generated an isogenic *in vitro* model by introducing the ALS4-associated c.1166T>C (L389S) mutation into human iPSCs using CRISPR-Cas9 genome editing, followed by differentiation into spinal MNs. This approach enables the analysis of disease-associated phenotypes in genetically matched cells and provides access to the neuronal population primarily affected in ALS. We selected the L389S substitution because it is one of the most penetrant ALS4-associated mutations. We generated one heterozygous clone (HETERO), carrying a mutation in a single SETX allele, and two homozygous clones (HOMO), in which both alleles were edited, to potentially enhance phenotypic penetrance (see Fig. S1A for Sanger validation and Methods for details). As a wild-type (WT) control, we used a clone that underwent the full genome-editing procedure but did not incorporate the mutation.

iPSCs were differentiated into MNs using a well-established protocol in which post mitotic MNs are generated at day 14 (*18*, *19*). As previously reported (*18*, *19*), we typically obtained >95% cells expressing the pan-neuronal marker neurofilament light chain (NEFL) and 60-80% expressing the MN-specific markers ISLET1 and MNX1 (Figure 1A-B). The remainder neuronal populations correspond mainly to interneurons. We did not detect any significant difference in the fraction of cells expressing neuronal and MN-specific markers between lines, indicating that the L389S substitution does not impact the efficiency of iPSC differentiation to MNs (Figure 1B). We next assessed whether this SETX mutation decreases MN viability using the reagent cMAC, which is metabolized and become fluorescent only in viable cells. We did not detect any significant decrease in the number of cMAC/ISLET1 positive cells between mutant and WT MNs cultured for 8 days between day 14 and day 22 indicating that the L389S substitution does not compromise MN survival between days 14 and 22 (Figure 1C).

**Figure 1:**
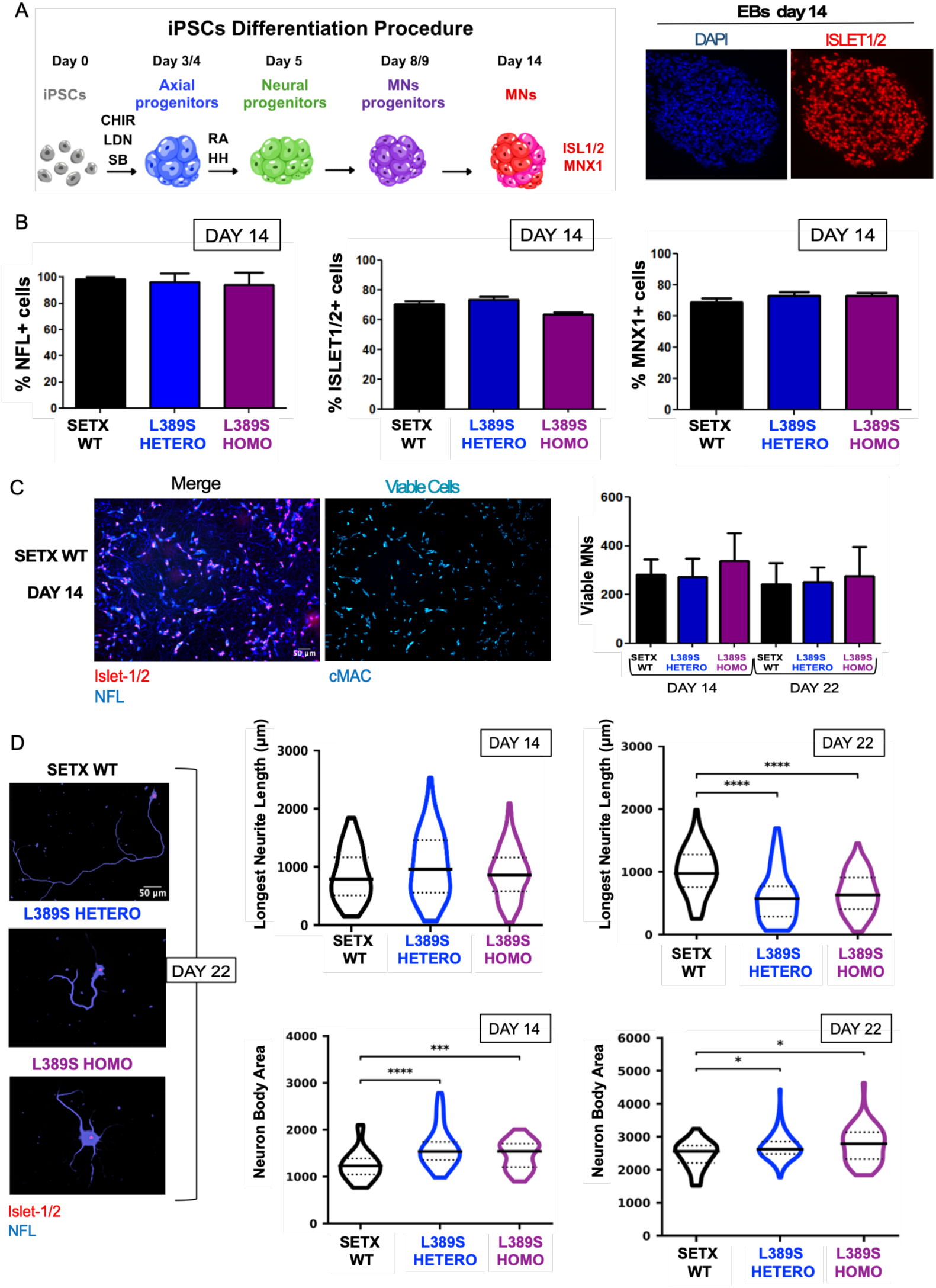
The L389S substitution causes a reduction of MNs axon length and changes in MNs morphology. **A)** Schematic representation of the procedure to differentiate iPSCs into mature MNs. The process passes through the stages of axial-like progenitors at day 3/4 induced by Wnt pathway activation with CHIR to neural progenitors at day 5, induced by Retinoic acid and Hedgehog (HH) signaling. MN progenitors are obtained at day 8/9 and by day 14, MNs express the specific markers ISL1/2 and MNX1. On the right, representative immunofluorescence images of embryoid bodies (EBs) on day 14 showing nuclear DAPI staining (blue) and expression of the MN marker ISL1/2 (red). **B)** Quantitative analysis of MNs differentiation efficiency assessed by immunofluorescence at day 14. The histograms show the percentage of viable cells positive for the neuronal marker NEFL and for the MN-specific markers ISL1 and MNX1. For ISLET1 and MNX1 markers, data represent the mean and standard error (SE) from 2 independent experiments from one SETX WT (one HETERO and one HOMO clones. For NEFL, data represent the mean and SE or standard deviation (SD) of 2 independent experiments with one SETX WT (two replicates in total), one HETERO (four replicates in total) and two HOMO (four replicates in total) clones. For each experiment, on average 100 cells for ISL1/MNX1 and 150 cells for NEFL were analyzed (see methods). **C)** Analyses of MN viability at days 14 and 22 after differentiation. **Left**: Representative images of MN clones showing viable cells identified by the fluorescent dye cMAC (cyan). Right: histogram displaying the quantification of the number of viable MNs at days 14 and 22. Data represent the mean and SE or SD from 2 independent experiments performed on one SETX WT (two replicates in total), one HETERO (four replicates in total) and two HOMO (four replicates in total) clones. For each experiment, on average 150 cells were analyzed (see methods). **D)** Analyses of MN axon length and soma size over time. **Left:** Representative images of MN plated at low density at day 22, immunolabeled for ISL1/2 (red, nuclei) and NEFL (blue, soma and neurites). Top right: violin plots displaying the distribution of the longest neurite length per cell at days 14 and 22. Bottom right: violin plots displaying the quantification of MN body area. Median values are indicated by solid central black line, and interquartile ranges by dashed lines. Data were pooled from 2 independent experiments performed on one SETX WT (two replicates in total), one HETERO (two replicates in total) and two HOMO (two replicates in total) clones. For each experiment, an average of 30 cells were analyzed per condition. *, P < 0.05; **, P < 0.01; ***, P < 0.001; ****, P < 0.0001 (Mann-Whitney U test, two-tailed).

Across ALS forms, a hallmark of early MN pathology is the impairment of axonal integrity. Axon structural abnormalities have been documented in SOD1 mutant mouse models (*20*), and axon shortening has been observed in iPSC-derived MNs carrying *TARDBP* mutations (*21*), establishing axon deficits as an early and tractable disease-relevant readout. Therefore, we assessed the effect of SETX ALS4 mutations in axon outgrowth, by analysing neurites length in mature MNs at days 14 and 22 after iPSCs differentiation (Figure 1D). Interestingly, while at day 14 ALS4 MN neurites are not significantly different from control MNs, at day 22 we detected a reproducible shortening of the longest neurite, which typically corresponds to the axon, in both heterozygous and homozygous L389S mutant clones (Figure 1D). This shortening could be due to axon degeneration instead of reduced axon outgrowth since the average size of mutant MNs axons was shorter at day 22 compared to day 14. We observed a similar behavior when monitoring the total axonal tree length (Figure S1B). We also found that the cell body of mutant MNs tended to have a larger size (Figure 1D), possibly reflecting perturbations in cell adhesion properties and/or the interaction with the extracellular matrix.

As mentioned above, the formation of cytoplasmic protein-RNA aggregates enriched in the TDP-43 protein represent a general hallmark of ALS as they have been found in post-mortem MNs in 98% of ALS patients including ALS4 patients (*22*, *23*). This is typically accompanied by TDP-43 relocation from the nucleus to the cytoplasm and it has been proposed that such aggregates play an important role in ALS pathogenesis (*24*). To assess whether our *in vitro* ALS4 model presents alterations in TDP-43 cellular localization, we monitored TDP-43 cellular distribution in WT and mutant MNs by immunofluorescence (Figure S1C). We did not detect any noticeable difference in the repartition of TDP-43 signal in the nucleus and the cytoplasm or formation of condensates/aggregates in the cytoplasm of mutant MNs. This result suggests that our model represents an early stage of ALS4 disease and indicates that the axonal phenotype of mutant MNs is not a consequence of the accumulation of toxic TDP-43 aggregates.

Together, these results establish a robust human ALS4 MN model in which early axonal defects arise in the absence of overt differentiation or TDP-43 localization abnormalities, thereby providing a framework to dissect the primary molecular events underlying disease onset.

### ALS4 MNs exhibit substantial gene expression dysregulation and compromised identity

SETX dysfunction has been shown to impact transcription and transcription-associated processes (*5*). To identify changes in gene expression that could underlie the observed axonal phenotype, we compared the transcriptome of wild-type (WT) and mutant (HETERO and HOMO) MNs by RNA-seq (Figure 2) at day 14 after differentiation. Parallel immunofluorescence experiments indicated similar expression of the MN marker Islet1/2 in all samples, ruling out any significant variation in the differentiation rate of the different lines (figure S2A). Principal component analyses (PCA) indicated a reproducible behavior for the different biological replicates of each condition as well as a robust distinctive expression pattern in the mutants relative to the WT (Figure 2A). In addition, heatmap analyses of several neuronal, MN and non-neuronal markers confirmed very low expression of typical markers of iPSCs, pMNs and non-MN neuronal types that are normally not obtained with this differentiation procedure, whereas the expression of pan-neuronal and MN markers was robustly high and very similar in all the analysed clones (Figure 2B).

**Figure 2:**
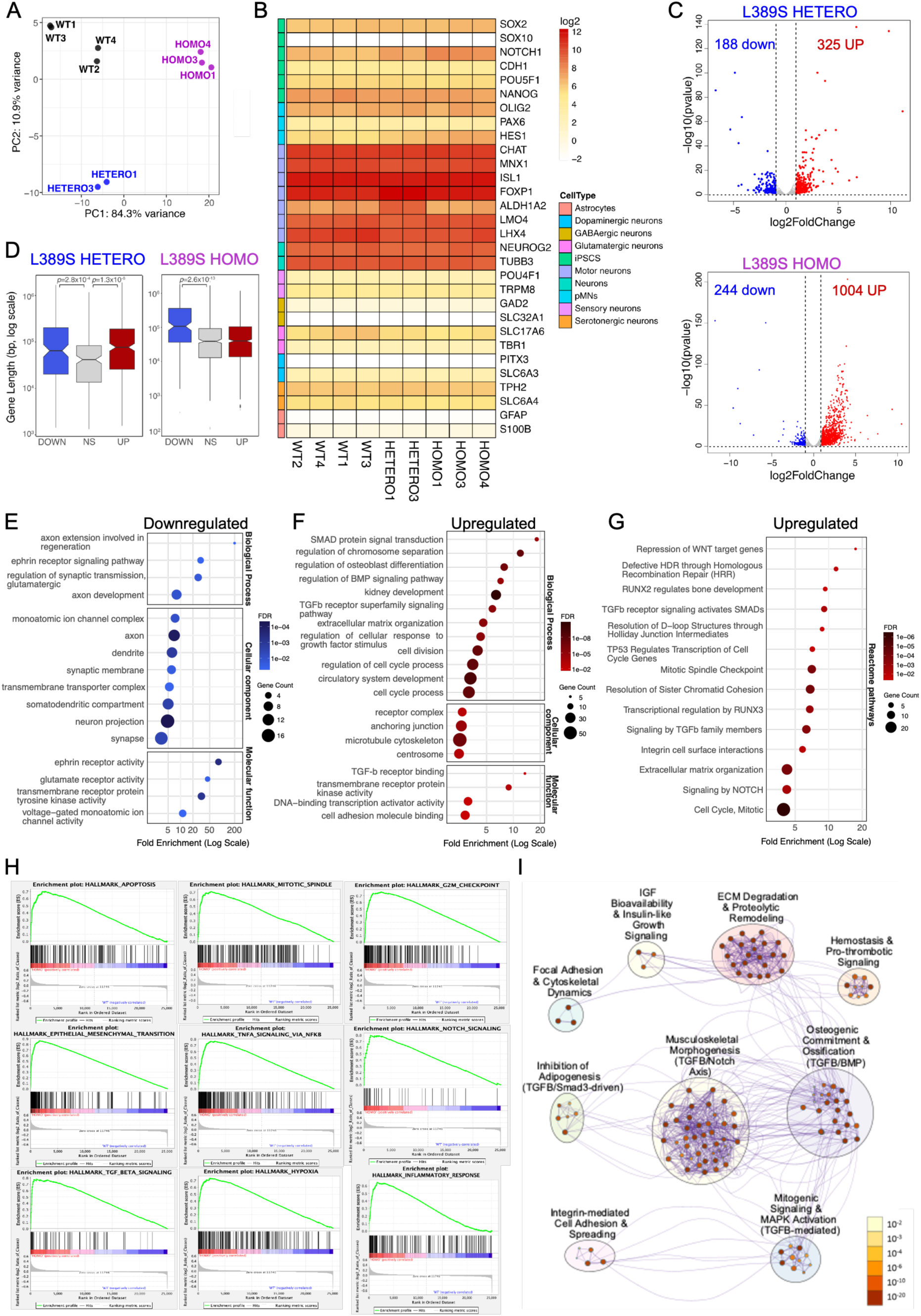
MNs carrying the L389S mutation present the deregulation of numerous genes and pathways. **A)** Principal Component Analysis (PCA) of RNA-seq samples. PCA plot illustrating the global transcriptomic variance between wild-type (WT), *SETX* L389S heterozygous (HETERO), and *SETX* L389S homozygous (HOMO) MNs. **B)** Expression profile of neuronal and non-neuronal specific markers. Heatmap displaying log2 expression levels of key markers for pluripotency (iPSCs), motor neuron progenitors (pMNs), and various neuronal subtypes (e.g., MNs, GABAergic, Glutamatergic). **C)** Differential gene expression analyses of L389S mutants performed by the DESeq package. Volcano plots showing significantly upregulated (UP, red) and downregulated (down, blue) genes in L389S HETERO (top) and L389S HOMO (bottom) MNs compared to WT. Numbers indicate the total count of differentially expressed genes (DEGs) based on a threshold of |log2FC| ≥ 1 and pvalue ≤ 0.05. **D)** Correlation between gene length and differential expression. Boxplots illustrate the distribution of gene lengths (base pairs, log10 scale) for downregulated (blue), non-significant (NS, grey), and upregulated (red) genes. To account for the high number of non-significant genes, the NS group represents a random subsample representative of the mean size of the deregulated groups. The horizontal line within each box indicates the median gene length, while the notches represent the 95% confidence interval for the median. Box boundaries represent the first and third quartiles (interquartile range, IQR), and whiskers extend to 1.5 times the IQR. Statistical significance was determined using a permutation-based Wilcoxon rank-sum test (100 iterations); reported p-values represent the median p-value across all permutations. **E–G)** Functional enrichment of common deregulated genes. Gene Ontology (GO) and Reactome pathway analyses performed on genes deregulated in both L389S HETERO and HOMO mutants. For all dot plots, the x-axis represents Fold Enrichment (Log Scale); dot size corresponds to the number of genes associated with each term; and the color gradient represents the False Discovery Rate (FDR) Plots correspond to a selection of representative categories, the full list of enriched classes are provided in Data S2. **H)** Gene Set Enrichment Analysis (GSEA) of hallmark pathways in ALS4 MNs. Representative GSEA enrichment plots for 9 of the top 10 significantly enriched (pvalue < 0.001) hallmark pathways in L389S HOMO mutant MNs. The same pathways were found significantly enriched for the HETERO mutant (see full list of enriched pathways for both types of mutants in Data S3 and S4). The green line denotes the enrichment score (ES) across the ranked gene list, with the peak indicating the maximum ES. Black bars represent individual genes from the hallmark set. The clustering of hits at the left (red/positive correlation) indicates significant upregulation in the mutant genotype. The grey profile shows the log2 fold-change for all genes, ordered from most upregulated (left) to most downregulated (right). **I)** Metascape enrichment analysis of genes upregulated (log2FC >0.9) in L389S HOMO mutant MNs compared to WT controls (p-value < 0.01, minimum enrichment > 1.5 and a minimum overlap of 3 genes). Network nodes represent enriched terms, with its size proportional to the number of genes mapped to that term. Nodes are grouped into clusters based on membership similarity (Kappa scores > 0.3). Each cluster is labeled according to its predominant biological theme. Nodes are colored by their p-value whith darker shades indicating higher statistical significance. Edges (purple lines) represent similarities between terms; thicker edges indicate a higher degree of shared genes between nodes.

Strikingly, we detected about 500 genes that are significantly deregulated (|log2FC| ≥1, pvalue ≤0.05) in heterozygous mutant clones compared to the corresponding WT isogenic control, and more than 1000 genes deregulated in homozygous mutant clones, consistent with an exacerbated phenotype when the two *SETX* alleles carry the L389S mutation (Figure 2C). While the heterozygous mutants presented relatively similar number of downregulated (188) and upregulated (325) genes, the expression pattern of homozygous clones was much more dominated by gene upregulation (i.e. 244 and 1004 genes appear down-and upregulated, respectively). A more detailed comparison of differentially expressed genes in the HETERO and HOMO mutants revealed a general similar trend in both, with about 500 genes being deregulated in the same direction in the two mutants, although in the homozygous clones the extent of the expression change is often higher and more numerous genes passed the pvalue threshold to be considered as significantly changed in expression (Figure S2B).

Consistent with the idea that the L389S mutation induces SETX gain of function, we did not detect the repression of the genes that were found to be downregulated in a previous study on SETX loss-of-function mutants (*25*), such as genes involved in autophagy, DNA repair or RNA processing (see list of deregulated genes in Data S1). Similarly, we did not detect an extension of 3’UTRs (Figure S2C), suggestive of transcription termination defects, that was also reported upon SETX depletion (*25*).

To search for features that might be enriched among genes differentially expressed in L389S MNs, we compared the length of up-and downregulated genes with that of not significantly deregulated genes (Figure 2D). We found that downregulated genes are significantly longer than genes without altered expression for both HETERO and HOMO mutants, suggesting a higher vulnerability of long genes to SETX dysregulated activity. Genes upregulated in HETERO mutants tend also to be longer, although downregulated genes include a higher proportion of very long genes, illustrated by the greater mean/median ratio (2.83 and 2.04 for downregulated and upregulated genes, respectively). The set of genes upregulated in HOMO mutants, which includes substantially more genes, was not characterized by a longer length. Importantly, in agreement with the axonal impairment that we observed in L389S MNs, gene ontology analyses of deregulated genes revealed that numerous genes associated with axon development and function are repressed, including for instance genes involved in the regulation of synapsis formation, axon regeneration, ephrin receptor activity and genes encoding axon and dendrites components (Figure 2E and full lists as Data S2). Interestingly, neuron-specific genes tend indeed to be particularly long (*26*).

In addition, although our analyses above indicate robust and efficient differentiation of mutant MNs, we observed upregulation of many genes that are not typically expressed in MNs, such as genes involved circulatory system and kidney development, regulation of chromosome separation and cell division. We also observed upregulation of genes related to the extracellular matrix (ECM) and cell adhesion as well as strong and highly significant enrichment of several GO terms related to the transforming growth factor β (TGF-β) superfamilly signaling pathway, which is known to regulate ECM genes (e.g. SMAD protein signal transduction, regulation of BMP signaling, TGF-β receptor binding…) (Figure 2F). Signaling by the TGF-β pathway involves the interaction of several types of cytokines such as TGF-βs, bone and morphogenetic factors (BMPs) and growth and differentiation factors (GDFs), among others, with specific membrane receptors to promote the phosphorylation and activation of SMAD transcriptional regulators. Activated SMADs then translocate to the nucleus and cooperate with other transcription factors to regulate the expression of multiple genes involved in development and tissue homeostasis (*27*). This result was particularly relevant since genes involved in the TGF-β superfamily signaling pathway were reported to be significantly upregulated in fibroblasts from ALS4 patients compared to healthy individuals and altered TGF-β signaling was observed in post-mortem neuronal tissues from patients (*16*). In this former report, upregulation of TGF-β genes was attributed to decreased expression of the *BAMBI* gene, encoding a TGF-β repressor. However, in our data *BAMBI* appears instead upregulated (see Data S1), pointing at a different mechanism underlying increased expression of TGF-β pathway genes in MNs.

The analysis of Reactome pathways enriched in the set of upregulated genes confirmed a signature suggestive of increased signaling of the TGF-β pathway. This was reflected not only by canonical pathway terms (e.g. “signaling by TGF-β family members”) but also by the enrichment of downstream effectors and cross-talk partners of the pathway. Specifically, we observed enrichment of “NOTCH signaling,” which is known to be synergistically activated by TGF-β cascades (*28*). We also detected pathways centered on the SMAD-responsive transcription factors RUNX2 and RUNX3 (*29*). Finally, this signature included structural processes downstream of TGF-β, such as “Integrin cell surface interactions” and “extracellular matrix organization” (Figure 2G). These analyses also confirmed a signature compatible with aberrant cell cycle reactivation (e.g. “mitotic spindle checkpoint” and “cell cycle, mitotic”) as well as a P53-related signature.

To explore more deeply the functions and interconnections of the genes deregulated in ALS4 MNs, we conducted complementary gene set enrichment analyses (GSEA) using the MSigDB Hallmark gene sets. As expected, an almost identical set of pathways appeared enriched in the HETERO and HOMO mutants (see representative pathways selected among the top 10 in Figure 2H and full lists in Data S3 and S4). Importantly, GSEA further supported and expanded the transcriptional signature revealed by GO and reactome pathway analyses. Notably, we detected again enrichment of G2-M checkpoint and mitotic spindle gene sets consistent with aberrant activation of cell cycle–related programs in post-mitotic neurons. This was accompanied by increased expression of apoptosis-related genes, indicating proximity to a pro-death threshold, as well as activation of inflammatory and stress-associated pathways, including TNFα signaling via NF-κB, complement, and broader inflammatory response signatures, together with enrichment of hypoxia, which suggests metabolic stress. In parallel, NOTCH, and epithelial–mesenchymal transition (EMT) pathways point to unstable neuronal identity and possible stress-induced transcriptional reprogramming. Because TGFβ signaling is known to directly activate EMT transcription factors (*30*) and functionally synergize with the NOTCH cascade (*28*), hyperactivation of the upstream TGF-β pathway could be a driver of the upregulation of NOTCH and EMT programs observed in L389S MNs. Finally, Metascape network-based enrichment analyses of upregulated genes further highlighted that transcriptional alterations in mutant MNs are organized around a central TGF-β signaling hub driving ECM remodeling and partial activation of non-neuronal gene expression programs (Figure 2I).

Altogether, our transcriptomic analyses indicate that expression of the L389S SETX mutant allele does not simply alter a restricted subset of neuronal genes but instead induces a broad transcriptional reprogramming in MNs indicative of partial loss of neuronal identity. This reprogramming is characterized by repression of long, neuron-specific genes, together with activation of stress-responsive, inflammatory, and non-neuronal pathways likely in connection with maladaptive signaling. These findings suggest the gain-of-function mutation L389S creates a vulnerable cellular state that may predispose MNs to degeneration.

### ALS4 MNs share transcriptional alterations with other forms of ALS

In order to assess whether ALS4 shares gene deregulation patterns with other familial forms of ALS, we compared our RNA-seq data with available datasets corresponding to transcriptomic analyses conducted in iPSC-derived MNs carrying ALS-causing mutations in either *C9ORF72, FUS, SOD1* or *TARDBP* gene, as well as a sporadic form of ALS and a PAN-ALS sample (see table 1 for dataset details). For all these datasets, iPSCs were generated from ALS patients and compared to different sets of healthy controls. To search for possible common signatures, we performed GSEA using the MSigDB Hallmark gene sets, which we found to be quite robust despite the high variability among the various datasets in terms of the type of control (isogenic for L389S datasets vs non-isogenic for the others) and differentiation procedure/time employed. First of all, we found an enrichment in the p53 pathway in all datasets, including both SETX L389S ones, except for the SOD1 dataset, supporting the idea that genome instability could be a general feature of non-SOD1 ALS, as previously proposed (*31*). Overall, the transcriptomic profile of MNs carrying the C9ORF72 mutation, which is the most frequent form of familial ALS, was the most similar to that of SETX L389S mutants, sharing the enrichment of pathways related to inflammation (e.g. inflammatory response, TNFα signaling via NF-κB), stress responses (e.g. KRAS signaling UP, hypoxia, p53 pathway, apoptosis), TGFβ signaling and compromised/unstable neuronal identity (i.e. EMT). Some upregulation of stress-related pathways was also detectable in mutant TARDBP MNs, while the aberrant cell cycle re-entry signature (i.e. upregulated mitotic spindle, E2F targets and G2/M checkpoint) was shared with mutant SOD1 MNs and to some extent also with FUS MNs. Together, these results indicate that our ALS4 model shares transcriptional signatures present in similar models of the most common familial forms of ALS.

**Table 1:** Summary of RNA-seq datasets of iPSC-derived MNs used in the ALS meta-analysis in. **figure 3A**. Public transcriptomic datasets integrated in the study were grouped according to the ALS mutation or disease category. All data were sourced from a previously published study (*31*), specifically by retrieving the differential expression results tables from the associated Shiny web application. For each mutation group, the table lists the associated GEO or repository accession identifiers, the total number of ALS samples, and the number of control samples reported in the original datasets. The Pan_ALS category represents the combined cohort including all mutation-specific and sporadic ALS datasets.

| **Group / Mutation** | **GEO / Dataset IDs** | **ALS samples (n)** | **Control samples (n)** | **References** |
|---|---|---|---|---|
| **C9orf72** | GSE52202, GSE143743, GSE139144, GSE168831, GSE201407, phs001231.v2.p1, AnswerALS | 60 | 27 | Sareen et al., 2013 (*33*); Abo-Rady et al., 2020 (*34*); Dafinca et al., 2020 (*35*); Catanese et al., 2021 (*36*); Sommer et al., 2022 (*37*); NeuroLINCS, 2022 (*38*); AnswerALS, 2022 (*39*) |
| **SOD1** | GSE54409, GSE95089, PRJNA361408, phs001231.v2.p1, AnswerALS | 20 | 7 | Kiskinis et al., 2014 (*40*); Wang et al., 2017(*41*); Bhinge et al., 2017 (*42*); NeuroLINCS, 2022 (*38*); AnswerALS, 2022 (*39*) |
| **FUS** | GSE77702, GSE94888, GSE168831, GSE203168 | 14 | 7 | Kapeli et al., 2016 (*43*); De Santis et al., 2017 (*44*); Catanese et al., 2021 (*36*); Hawkins et al., 2022 (*45*) |
| **TARDBP** | GSE147544, PRJEB47567 | 9 | 6 | Dafinca et al., 2020 (*35*) ; Smith et al., 2021 (*46*) |
| **Sporadic ALS** | phs001231.v2.p1, AnswerALS | 208 | 56 | NeuroLINCS, 2022; AnswerALS, 2022 (*39*) |
| **Pan_ALS (all studies combined)** | All datasets above | 311 | 103 | All references listed above |

To further expand these findings, we also compared the set of genes deregulated in the L389S homozygous mutant, which presents the most extreme transcriptional phenotype, with transcriptomic datasets generated from MNs biopsies from ALS patients suffering from either a sporadic form of this disease or the C9ORF72-associated familial ALS form (figure 3B and Data S5). We detected a highly significant group of 48 genes that appeared deregulated in all four datasets analyzed. Interestingly, GO analyses of this core genes revealed a very similar signature to that observed in L389S MNs including TGF-β signaling, processes controlled by this superfamily pathway (e.g. osteoblast differentiation, ECM, cell adhesion) or pathways downstream of (e.g. ERK, *32*) or synergizing with TGF-β (e.g. NOTCH pathway) and other processes suggestive of compromised neuronal identity (e.g. eye development, nephron development) and post-mitotic fate (e.g. positive regulation of cell proliferation).

**Figure 3:**
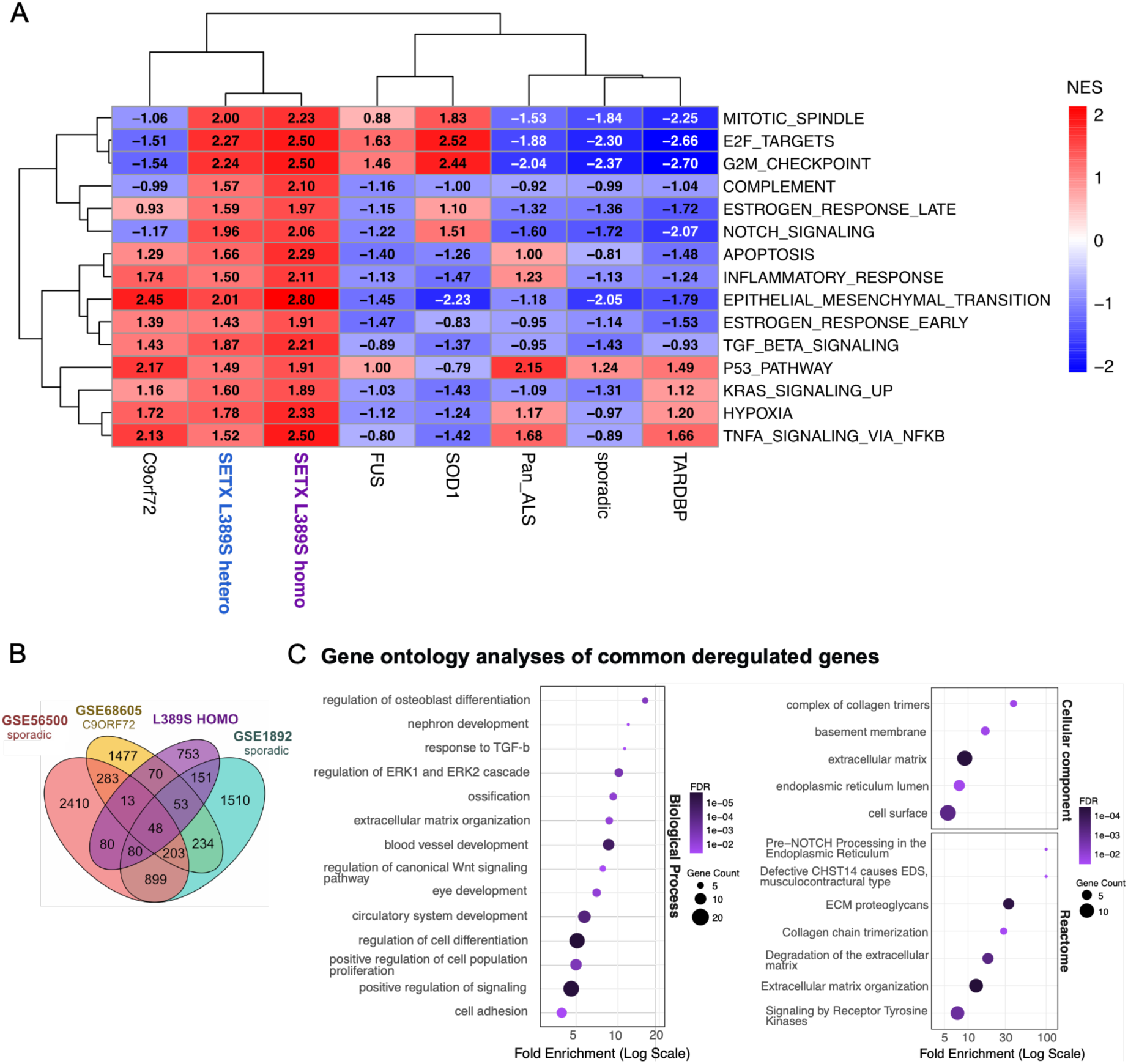
Comparison of gene deregulation patterns in ALS4 MNs and iPSC-derived MNs and post-mortem tissues with other familial or sporadic ALS forms. **A)** Gene Set Enrichment Analysis (GSEA) of hallmark pathways across ALS genotypes. Heatmap displaying Normalized Enrichment Scores (NES) for selected MSigDB hallmark gene sets across multiple iPSC-derived MN datasets, including *C9orf72*, *SETX* (L389S heterozygous and homozygous), *FUS*, *SOD1*, and *TARDBP* mutations, alongside Pan-ALS and sporadic samples (see table 1 for details). Red indicates positive enrichment (upregulated pathways), and blue indicates negative enrichment (downregulated pathways). Dendrograms represent unsupervised hierarchical clustering of genotypes (top) and pathways (left) based on NES profiles. The heatmap has been generated for pathways with padj <0.1 for both L389S heterozygous and homozygous mutants. **B)** Overlap of differentially expressed genes across independent ALS cohorts. Venn diagram illustrating the intersection of genes deregulated (|log2FC| ≥ 1 and pval ≤0.05) in the L389S HOMO mutant and in three transcriptomic datasets generated from post-mortem tissues of ALS patients (see methods for details). A core set of 48 genes was found to be commonly deregulated across all four datasets (pval <10^-10^). **C)** Functional annotation and pathway enrichment of the 48 core deregulated genes. Dot plots showing Gene Ontology (GO) enrichment for Biological Processes (left) and Cellular Components/Reactome pathways (right). The x-axis represents the fold enrichment on a log10 scale. Dot size corresponds to the number of genes associated with each term, while the color gradient represents the False Discovery Rate (FDR), with darker purple indicating higher statistical significance. Plots correspond to a selection of representative categories, the full list of enriched classes is provided in Data S5.

Altogether, our results suggest that deregulated TGF-β signaling and compromised neuronal identity are key features of not only ALS4 but also other forms of familial and sporadic ALS.

### The L389S mutation causes widespread splicing alterations that are largely uncoupled from changes in transcript abundance

Previous studies on other forms of familial and sporadic ALS have revealed substantial splicing alterations in ALS patient tissues (*47*, *48*). To assess whether the SETX ALS4 mutation L389S also provokes perturbations in pre-mRNA splicing, we analysed our RNA-seq data using the rMATS package (see methods for details). We detected a total of 410 exons in different 343 genes undergoing alternative splicing in the HETERO mutant and 478 exons with differential splicing in 404 genes in the HOMO mutant (figure 4B and D and Data S6). The majority of splicing alterations consisted in exon skipping, with a moderately higher tendency for the mutants to exhibit higher exon inclusion rates, followed by alternative 5’ and 3’ SS usage. Intron retention and differential usage of mutually exclusive exons remained marginal. Most of these splicing alterations are likely caused directly by aberrant SETX activity at nascent RNAs since we only detect significant changes in the expression of very few genes encoding splicing regulators in the mutants (2 genes encoding RNA-binding proteins for the HETERO and 4 genes for the HOMO, see Data S1). Strikingly, when we compared the list of genes presenting splicing alterations with those exhibiting changes in expression we observed almost no overlap, indicating that the perturbations in RNA levels and in pre-mRNA splicing are independent outcomes of mutant SETX dysregulated activity (figure 4C and E).

**Figure 4:**
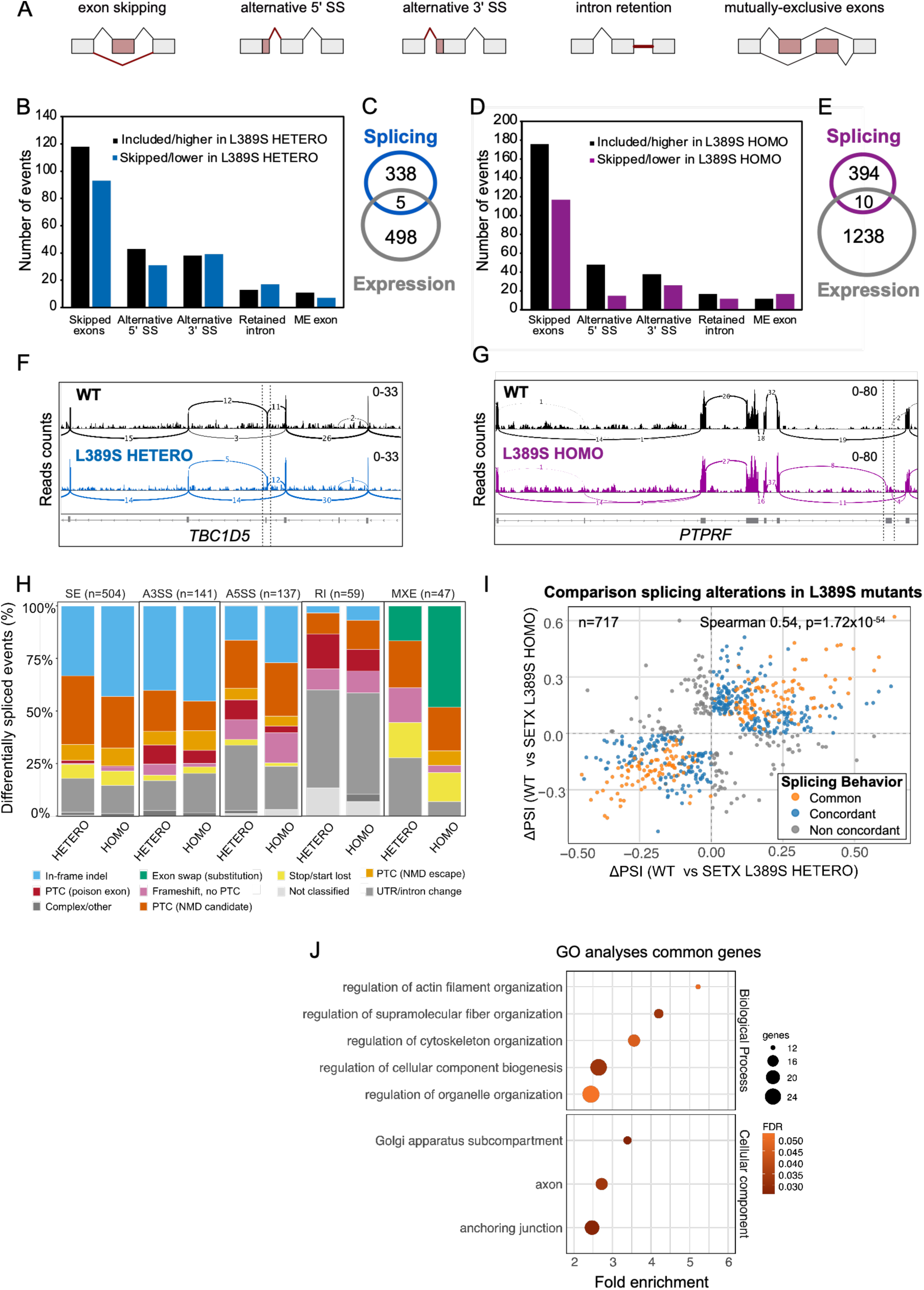
the *SETX* L389S mutation induces numerous pre-mRNA splicing alterations. **A)** Schematic representation of the five alternative splicing (AS) categories analyzed: exon skipping, alternative 5’ or 3’ splice site (SS) usage, intron retention, and mutually exclusive (ME) exons. **B–E)** Global analyses of splicing deregulation in *SETX* L389S mutants. Bar plots (B, D) quantify the distribution and directionality of statistically significant AS events (|ΔPSI| > 0.1 and FDR < 0.05) in L389S HETERO and HOMO MNs compared to WT, identified via rMATS (see Data S6). Venn diagrams (C, E) illustrate the minimal overlap between genes encoding transcripts exhibiting differential splicing (“Splicing”) and those with differential abundance (“Expression”), suggesting that splicing defects are largely independent of overall transcript levels. **F, G)** Integrative Genomics Viewer (IGV) snapshots of representative transcripts exhibiting alternative splicing in ALS4 mutants. Sashimi plots displaying RNA-seq read coverage and junction spanning across specific loci. Numbers on the curved lines indicate the count of junction-supporting reads for each condition. The *TBC1D5* mRNA is an example of higher exon skipping in HETERO mutants, while *PTPRF* is an example of higher exon inclusion in HOMO mutants. Exons undergoing AS are indicated by flanking dashed lines. **H)** Bar chart summarizing the predicted outcomes of splicing alterations on protein or transcript sequence. **I)** Scatter plot showing the ΔPSI of AS events between SETX L389S HETERO and HOMO mutants relative to WT (|ΔPSI| > 0.1 and FDR < 0.05 in at least one mutant). Splicing events are categorized based on concordance between genotypes: common (orange; same ΔPSI sign and nominal pvalue ≤ 0.05 in both mutants; n=225), concordant (blue; same ΔPSI sign and FDR < 0.05 in at least one mutant, excluding « common »; n=351), and non-concordant (grey; opposite ΔPSI sign; n=141). **J)** Functional enrichment of common genes carrying differentially spliced transcripts. Gene Ontology (GO) dot plot where the x-axis indicates fold enrichment, dot size reflects gene count, and color indicates statistical significance (FDR).

The majority of the detected splicing alterations are predicted to alter protein sequence (figure 4H and Data S7; e.g. in-frame insertions and deletions of coding sequence, inclusion of premature stop codons, frameshifts), while a smaller subset of them lead to changes in UTR or intronic sequences potentially altering RNA regulatory steps (e.g. RNA processing, localization, etc). Albeit we observed moderate overlap (15-18%, *p*=7.8×10^-41^) between the genes exhibiting splicing alterations passing the aforementioned ΔPSI and FDR thresholds in the HETERO and the HOMO mutants (Figure S3A-B), a more detailed analysis revealed a substantial global correlation between the two mutants (figure 4I) with about 80% of splicing alterations exhibiting a concordant behavior. Using more relaxed criteria based on concordance and nominal pvalue (figure 4I) we defined a list of 197 genes with common splicing alterations and we performed gene ontology analyses to search for functional enrichment. Among the functional categories that appear overrepresented in this gene set, we detected several ones that are particularly relevant for axonal structural integrity and synapsis formation and maintenance such as regulation of actin filament and cytoskeleton organization, axon components and anchoring junction (figure 4J). Moreover, when we focused on the stricter list of genes presenting significant splicing alterations in both mutants, we found that about half of them correspond to genes associated with neurodegenerative diseases (e.g. ALS, Alzheimer’s and Parkinson’s disease) and/or intellectual/psychological disorders (e.g. autism, schizophrenia, etc), supporting a role for the encoded proteins in the correct function of the nervous system.

These results suggest that, in addition to the above-mentioned perturbations in gene expression, the ALS4 mutation L389S can also induce changes in the identity of a subset of proteins involved in processes that are relevant for MN functions, potentially contributing the axonal and morphological defects of ALS4 MNs.

### R-loop-forming regions are associated with transcriptional and splicing alterations in ALS4 MNs

A previous study reported a global decrease in R-loops levels in fibroblasts from ALS4 patients carrying the L389S substitution compared to healthy individuals (*16*) and proposed that this reduction in R-loops plays an important role in gene deregulation in ALS4. To assess whether the L389S mutation provokes a global decrease of R-loop levels in human MNs, we performed slot blot analyses on genomic DNA preparations from WT, HETERO and HOMO mutant MNs using the S9.6 antibody that specifically recognizes RNA:DNA hybrids. Because this antibody can also bind double-stranded RNAs (dsRNAs), although with lower affinity, we treated our samples with RNase III, which degrades dsRNAs, and we loaded in parallel samples treated with RNase H, which degrades RNA:DNA hybrids, as a negative control (figure S4). We detected low, although quantifiable, signal in MN samples that completely disappeared upon RNase H treatment, confirming that it corresponded to RNA:DNA hybrids. Subsequent hybridization of blots with an antibody recognizing dsDNA was used as a loading control. We did not observe any significant difference in R-loop levels in any of the mutants compared to the WT, indicating that the L389S substitution, either in heterozygosis or in homozygosis, does not provoke a global reduction of cellular R-loop levels. This result suggests that other features of ALS4 patient cells rather than, or in addition to, the L389S mutation might underlie the reported decreased R-loop levels. We noticed that the mentioned study employed fibroblasts, which are proliferative cells, and R-loops are known to accumulate at sites of transcription-replication conflicts (*6*). To investigate the possibility that R-loops globally decrease only in dividing cells carrying the L389S mutation, we conducted slot blot analyses on WT and mutant iPSCs, MN progenitors (pMNs) and MNs in parallel. Strikingly, we detected more than 10-fold higher levels of R-loops in iPSCs compared to pMNs, which have a lower proliferative capacity, and MNs, which are post-mitotic (figure S4C), supporting the idea that DNA replication has a strong impact on cellular R-loop levels. However, in none of the three isogenic cell types analysed we detected any significant difference in global R-loop levels in the mutants relative to the WT.

The results above do not exclude the possibility that L389S SETX displays aberrant R-loop resolving activity at a subset of R-loops that is not large enough to produce global changes in the levels of these structures. Indeed, major sources of R-loop cellular levels are rRNA loci and mitochondria, and hence alterations in a subset of genomic R-loops would not necessarily be expected to have a major global effect. We reasoned that if R-loop alterations are the cause of part of the changes in gene expression and RNA splicing observed in L389S mutants, we might detect some enrichment for genes/transcripts with particularly high propensity to form R-loops among genes with differential expression or splicing. To address this possibility, we computed and compared an R-loop propensity score for genes that are significantly upregulated, downregulated or not differentially expressed in L389S mutants relative to the WT. Such score was calculated as the total R-loop signal measured by the DRIP-seq technique, which employs the S9.6 antibody to immunoprecipitate R-loops, normalized by the gene length and the transcription levels using RNA-seq datasets generated in the same study as a proxy for transcription. For these analyses we used previously published datasets generated in HEK293 cells that are particularly robust (*49*, *50*). These datasets correlated well with DRIP-seq datasets generated in iPSC-derived neurons (*51*), indicating a very similar overall R-loop distribution in both cell types (figure S5A). However, the iPSC-derived neurons datasets presented a relatively poor correlation with matched RNA-seq data, unlike the HEK293 DRIP-seq data, making them less suited for the computation of R-loop propensity (figure S5B-C). We therefore used HEK293 datasets and only focused on genes that are expressed both in MNs and HEK293 cells. Interestingly, we found that for both the HETERO and the HOMO mutants, deregulated genes tended to have a higher R-loop propensity score (figure 5A). This potential link between R-loop enrichment and sensitivity to SETX L389S mutation suggests that the expression of a subset of genes might be influenced by R-loop formation and that L389S SETX abnormal activity at these R-loops could possibly underlie their transcription changes.

**Figure 5:**
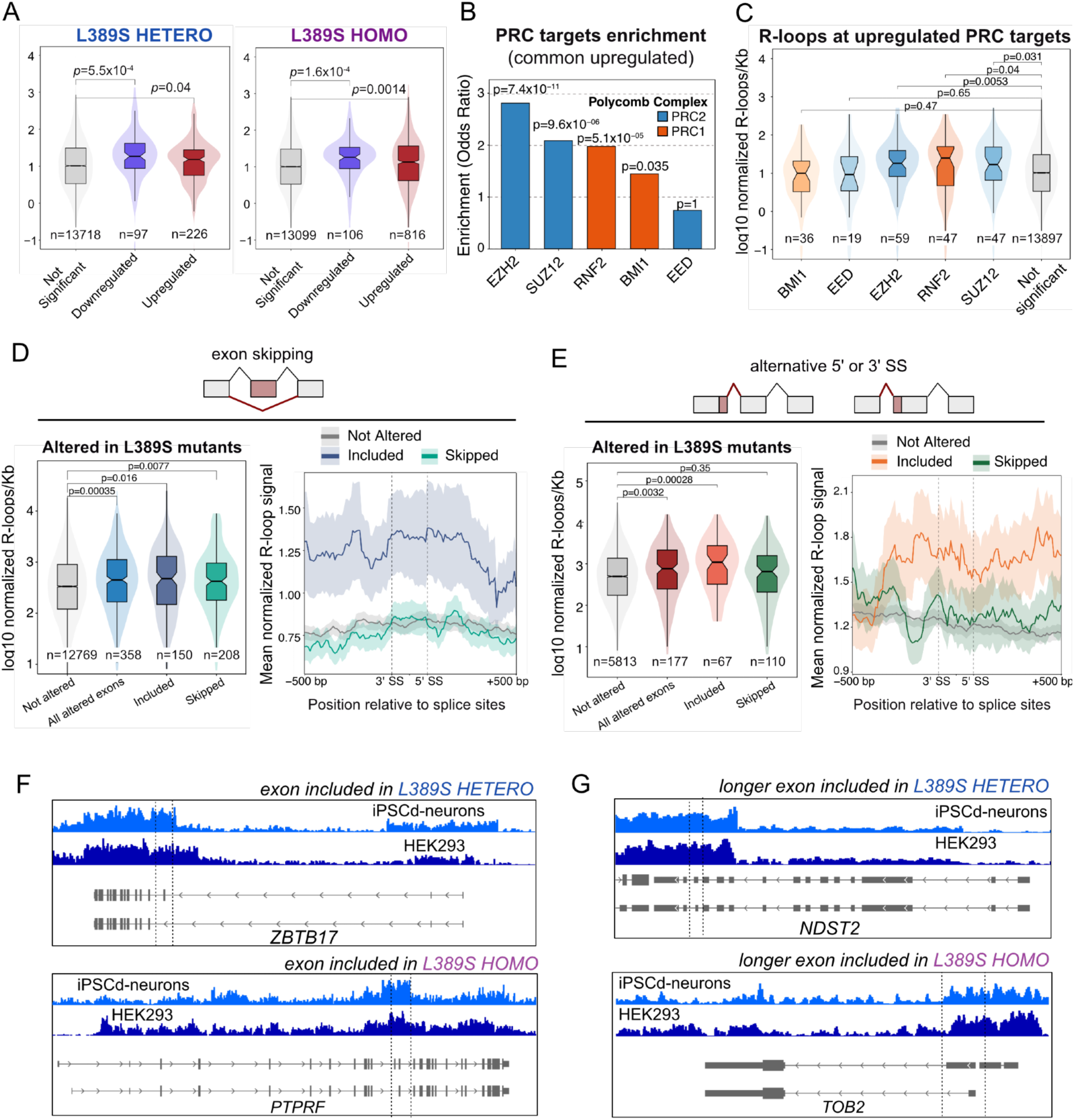
Genes with altered expression and exons undergoing alternative splicing in ALS4 MNs are enriched for R-loop-forming regions. **A)** Correlation between R-loop propensity and differential expression. Boxplots illustrate the distribution of R-loop propensity score (R-loop signal/Kb normalized by transcription) for downregulated (blue), not significantly deregulated (grey), and upregulated (red) genes in the L389S HETERO (left) and HOMO (right) mutants. The horizontal line within each box indicates the median, while the notches represent the 95% confidence interval for the median. Box boundaries represent the first and third quartiles (interquartile range, IQR), and whiskers extend to 1.5 times the IQR. Statistical significance was determined using a Wilcoxon rank-sum test (*p* denotes the p-value, ns= not significant). **B)** Enrichment of Polycomb Repressive Complexes (PRC) components target genes among common upregulated genes in both heterozygous and homozygous mutant MNs. *p* denotes the p-value of a Fisher exact test. PRC1 components are indicated in orange whereas PRC2 components are shown in blue. **C)** R-loop propensity at PRC target genes that are commonly upregulated in both L389S mutants compared to not significantly deregulated genes. Box plots and statistical analyses performed as in **A**). **D-E)** Correlation between R-loop propensity and splicing alterations. Left: boxplots representing the distribution of R-loop propensity score for exons not exhibiting significant splicing alterations (not altered) or with differential exon skipping (**D**) or alternative 5’ or 3’ SS usage (**E**). For alternative 5’/3’SS usage “included” or “skipped” refers to the higher or lower rate of inclusion of the longer version of the exon, respectively. Statistics and formatting of boxplots as in **A**). Right: metagene analyses of R-loop propensity at differentially spliced exons and flanking regions. Exons are scaled to 200 bp. Solid lines represent mean values, and shaded ribbons denote the standard error of the mean (SEM). **F-G)** IGV snapshots of examples of exons displaying AS in ALS4 mutants and with particularly high R-loop levels compared to other exons of the same gene. **F**) Examples of exons with higher rate of inclusion in L389S mutants. **G**) Examples of exons with higher inclusion of the longer form in mutant MNs. The histograms correspond to DRIP-seq datasets generated in the indicated cell lines (iPSCd-neurons denote cell populations enriched in neurons and generated by differentiation of iPSCs). Dashed lines enclose the exon with differential splicing.

Previously proposed mechanisms of transcriptional regulation involving R-loops include the recruitment of regulatory complexes such as Polycomb repressive complexes (PRC) to target genes by R-loop structures (*52*, *53*). Interestingly, we found that PRC target genes were significantly enriched among genes upregulated in both L389S mutants (figure 5B), including numerous genes involved in the TGF-β signaling pathway (Data S8). Moreover, we found that PRC targets tend to exhibit a significantly higher R-loop propensity score, in particular targets of the PRC1 component RNF2 and of PRC2 components EZH2 and SUZ12 (figure 5C), a phenomenon that appeared exacerbated in the HOMO mutant, consistent with a dose effect (figure S5D). Together these data support a link between altered R-loop levels and derepression of PRC-regulated genes in mutant MNs.

As previously mentioned, ALS4 MNs also exhibited several hundreds of splicing alterations. Therefore, we conducted similar analyses as above on genes carrying exons with differential splicing in mutant MNs. We focused on exon skipping and alternative SS usage, since these were the most frequent events. When comparing the R-loop propensity of transcripts with differential exon skipping and transcripts without changes in this type of splicing event, we did not detect any significant difference (Figure S6A) for any of the mutants. However, when we computed the same score for only exonic regions, we found a moderate but significant increase in the median R-loop propensity for exons with altered skipping in L389S mutants (figure 5D). Individual mutants (i.e. HETERO and HOMO) presented the same trend as in the combined analysis (figure S6B). Metagene analyses of normalized R-loop signal at regions around the altered exons revealed that exons presenting higher inclusion rates in mutant MNs are embedded in regions with higher R-loop propensity than non-altered exons and exons with higher skipping rates in the mutants. This suggests that R-loop formation at those exonic regions might interfere with their recognition by the splicing machinery and that enhanced R-loop removal by mutant SETX could promote their inclusion. In the case of alternative 5’ or 3’ SS usage, we detected a higher R-loop propensity score computed on the whole transcript, particularly for transcripts with increased inclusion of the longer form of the exon (Figure S6A). This effect was more prominent when focusing on exons (figures 5E and S6B). Metagene analyses indicated that exons with higher inclusion of the longer exon form fall within extensive regions of high R-loop propensity, while for exons with higher skipping some local increase in R-loop propensity could be observed at flanking introns, suggesting that in some of these cases R-loops might modulate the access/action of splicing regulators binding these intronic regions. Consistently, visual inspection of R-loop mapping datasets generated in HEK293 cells (i.e. used for the mentioned analyses) and in iPSC-derived neurons revealed that exons with higher inclusion in mutant MNs often overlap with a region of particularly high R-loop signal (figure 5F-G), while exons with higher skipping in the mutants are instead adjacent to strong R-loop forming regions (figure S6C-D). Importantly, when we analysed the properties that typically correlate with differences in exon inclusion/skipping or SS usage (i.e. exon length, GC content, GC contrast and SS strength), we only observed a moderate decrease in exon length and slight increase in GC content at the 3’ SS region for exons with higher skipping in the HOMO mutant while all the other features remained similar in altered and non-altered exons indicating that higher R-loop propensity is the property that better explains a majority of splicing alterations caused by the L389S mutation (figure S7 and S8).

Altogether, these results suggest that the presence of R-loops at particular exons could modulate splicing and that L389S SETX action at those R-loops might thus have an impact on the splicing outcome, and therefore on the identity of the proteins produced.

### The L389S substitution does not provoke DNA damage accumulation but sensitizes to DNA damage induction

Several reports have shown that SETX depletion or loss of function results in the accumulation of R-loops and DNA damage (*54–57*). Consistent with the idea that the L389S substitution provokes a gain of function, we did not detect any significant increase in cellular R-loop levels in mutant MNs. However, increased DNA damage and genome instability have been observed in models of several neurodegenerative diseases, including ALS, as well as in patient cells, especially at advanced disease stages (*3*, *50*, *58*). Moreover, our transcriptomic analyses have revealed the upregulation of genes and pathways related to the DNA damage response such as the G2/M checkpoint and the p53 pathway (figures 2 and 3). Therefore, we set out to analyse whether L389S mutant MNs exhibit an accumulation of DNA damage. To this end, we conducted immunofluorescence experiments on WT and mutant MNs at days 14 and 22 after iPSC differentiation using antibodies against the DNA damage markers γH2AX and 53BP1 (Figure 6 and figure S9). Phosphorylation of the histone variant H2AX (i.e. γH2AX) is a sensitive and early marker of double-strand breaks (DSBs), while 53BP1 is recruited to DSBs later and promotes DNA repair by non-homologous end joining (*59*, *60*). A higher number of marker foci is typically considered as indicative of increased DNA damage. In both control and mutant MNs, we observed very few γH2AX or 53BP1 foci per cell, with the majority of cells having none or only one, at both timepoints (i.e. day 14 and 22) indicating that the basal level of DNA damage in MNs in culture is very low and is not increased by the L389S substitution in SETX.

**Figure 6:**
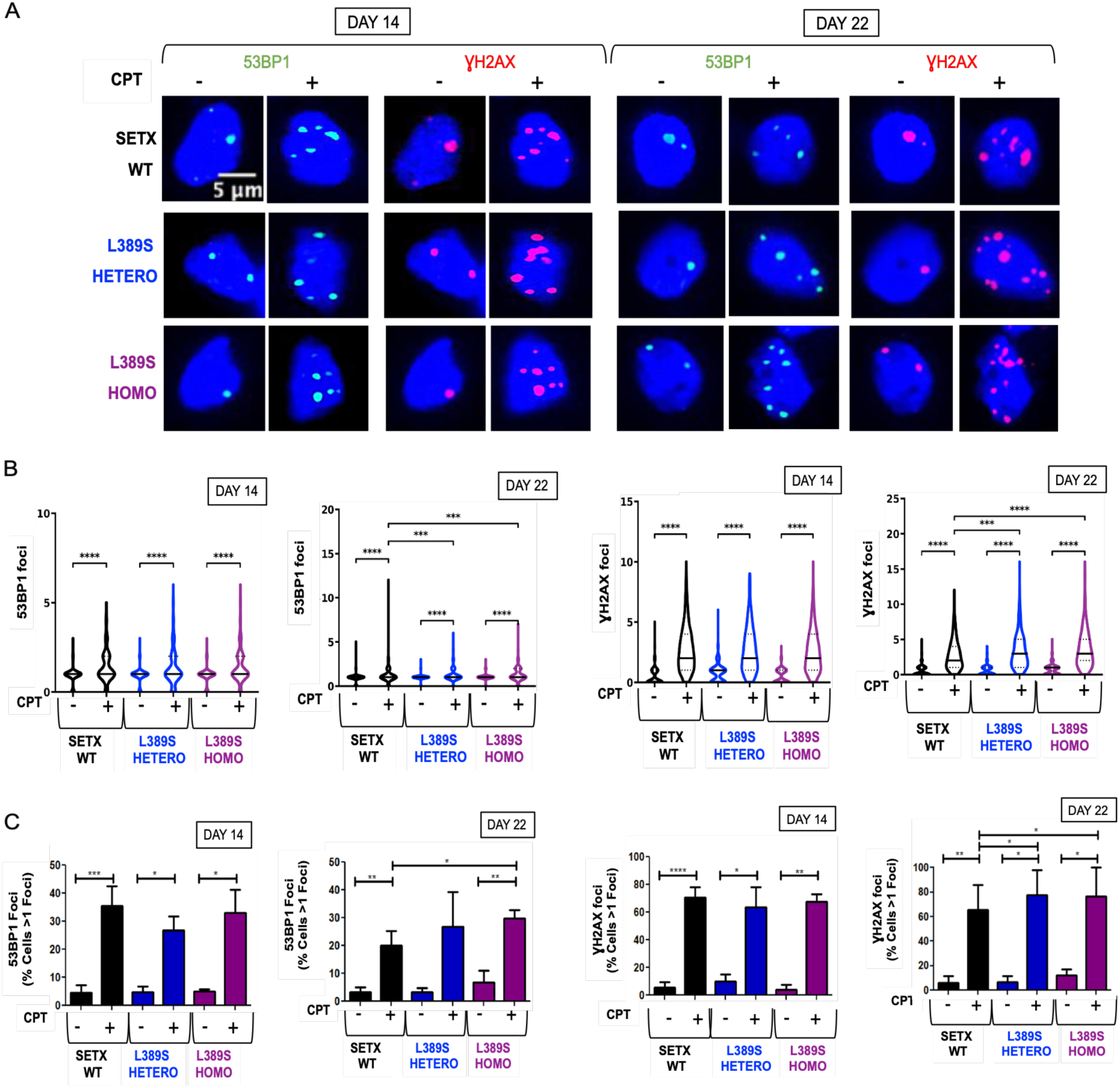
Analysis of DNA damage in L389S SETX MNs both in basal conditions and in the presence of the DNA damage inducer camptothecin (CPT). **A)** Representative immunofluorescence images showing nuclear foci of 53BP1 (green, foci) and γH2AX (red, foci) in ISL1/2-positive cells (blue, nuclei) of WT, HETERO and HOMO mutant clones at days 14 and 22 of differentiation. Cells were analysed under basal conditions (CPT-) and after treatment with CPT to induce DNA damage. **B)** Violin plots representing the distribution of the total number of 53BP1 and γH2AX foci in each ISL1/2-positive nucleus in absence and in presence of CPT in WT, HETERO and HOMO mutant clones at days 14 and 22. Median values are indicated by solid central black line, and interquartile ranges by dashed lines. The results show a significant increase in the DNA damage following treatment with CPT across all genotypes. Data were pooled from 3 independent experiments and correspond to two SETX WT (five replicates in total), one HETERO (three replicates in total) and one HOMO (three replicates in total) clones. Foci number count was evaluated by the creation of a dedicated Macro in Fiji (see methods for details). For each experiment, an average of 150 cells were analyzed per condition. *, P < 0.05; **, P < 0.01; ***, P < 0.001; ****, P < 0.0001 (Mann-Whitney U test, two-tailed). **C)** Histogram showing the quantification of the percentage of cells with > 1 nuclear focus for 53BP1 and γH2AX at days 14 and 22. Data represent the mean and SD for the same experiments as in panel B. *, P < 0.05; **, P < 0.01; ***, P < 0.001; ****, P < 0.0001 (Paired t-test, two-tailed). Paired t-test was used to account for batch effect.

Because our DNA damage assessment is based on the presence of DNA damage response factors, to verify that mutant MNs are not impaired in this response and that the number of DSB markers is reflecting the actual levels of DNA damage, we included as a control a treatment with camptothecin (CPT). This drug stabilizes the Topoisomerase I–DNA cleavage complex preventing religation of the cleaved DNA, which leads to accumulation of single-strand breaks that are converted to DSBs by other cellular processes (*61*). In the case of post-mitotic neurons, DSBs arise mainly from collisions between the transcription machinery and the Topoisomerase I–DNA cleavage complex. Upon CPT treatment, for both WT and mutant MNs we observed a robust increase in the number of γH2AX foci per cell, with some cells exhibiting more than 10 foci, and to a lesser, although still significant, extent in the number of 53BP1 foci. These results confirmed that the DNA damage response is not impaired in mutant MNs, and because no difference in DNA damage markers was observed at any timepoint in the absence of CPT, they seemingly exclude the possibility that the axonal phenotype of mutant MNs is due to DNA damage accumulation.

However, in the presence of CPT, we observed an increase in the fraction of cells with more than 2 foci (i.e. both γH2AX and 53BP1 foci) for mutant relative to WT MNs at day 22 (i.e. symptomatic stage), albeit not at day 14. These results underscore a higher vulnerability of mutant MNs to genotoxic stress. The pre-existing activation of stress, checkpoint, and apoptotic pathways revealed by our transcriptomic analyses (Figure 2H-I) possibly lowers the threshold for damage tolerance, thereby amplifying the conversion of transient lesions into persistent DNA double-strand breaks. Altogether, these results suggest a latent defect in genome maintenance mechanisms rather than constitutive DNA damage accumulation.

### Aberrant TGF-β signaling is a key driver of ALS4 MNs gene deregulation and axonal defects

As described in former section, our transcriptomic analyses revealed the upregulation of genes related to the TGF-β superfamily signaling pathway, a feature that is conserved in other forms of familial and sporadic ALS (see figures 2 and 3). We detected not only overexpression of genes encoding ligands, receptors and signal transducers, but also typical SMADs targets (see schematic model in figure 7A), including genes involved in EMT and ECM deposition and remodeling, supporting overactivation of the TGF-β pathway in ALS4 MNs. To assess this possibility, we analysed by immunofluorescence SMAD1/5/8 phosphorylation to monitor BMP activation and SMAD2 phosphorylation as a readout of active TGF-β signaling in WT and L389S mutant MNs at day 14 (i.e. before axonal symptoms, figure 7B and S10A). We found a significant increase in phosphorylated forms of the two regulators in both L389S heterozygous and homozygous MNs, confirming that TGF-β signaling is overactive in ALS4 MNs. These results, together with the gene expression analyses described above support the idea that increased activation of TGF-β signaling could be responsible for a substantial part of the gene deregulation phenotype of ALS4 MNs, which could play an important role in the axonal phenotype of mutant MNs. To further substantiate these notions, we treated WT and mutant cells with SB431542, an inhibitor of the kinase activity of TGFβ receptors, and LDN-193189, an inhibitor of BMP receptors (Figure 7A), from the MN progenitor stage to day 22 after iPSC differentiation and we analysed the expression of several genes that are deregulated in mutant MNs at day 14. We focused on the homozygous mutant, which exhibits stronger expression changes and allows more robust quantification. We found that the inhibitors substantially suppressed the upregulation of COL2A1, which is a marker of cartilage, FN1 and VIM, which are functional promoters of EMT, and more mildly IGFBP5, involved in angiogenesis and osteogenesis (figures 7C and S10B). Importantly, we also detected partial suppression of downregulation of the neurofilament component *NEFH* in the mutant, suggesting that overactivation of the TGF-β superfamily signaling pathway could also be at least partially responsible for the decreased expression of specific neuronal genes (figure 7C).

**Figure 7:**
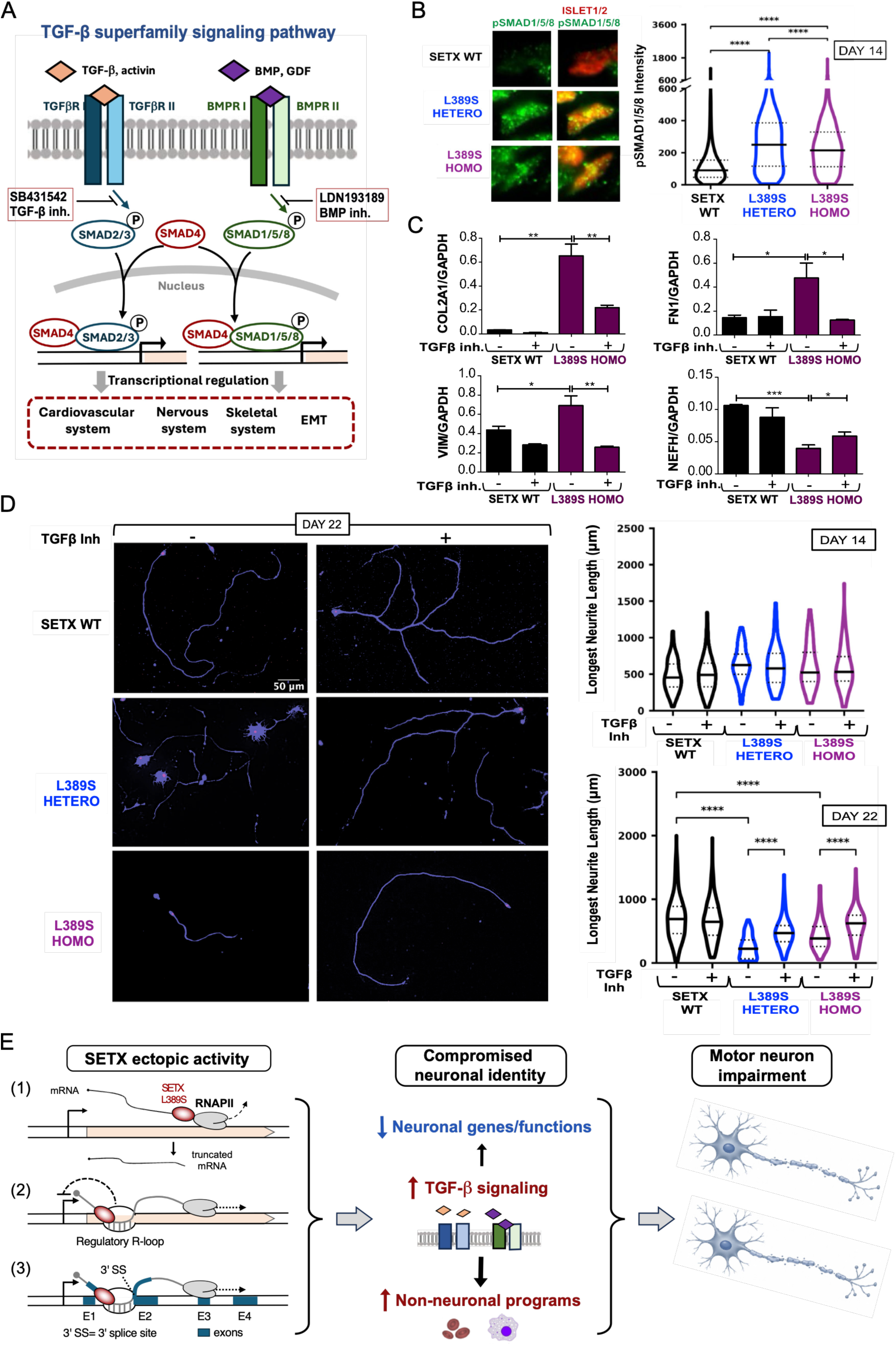
Suppressing TGF-β overactivation alleviates ALS4 MNs phenotypes. **A)** Schematic representation of the TGF-β signaling pathway mediated by two branches, from one side via SMAD2/3 by TGF-β/Activin receptors and from the other side via SMAD1/5/8 by BMP/GDF receptors. Specific inhibitors SB431542 (TGF-β Inh.) and LDN-193189 (BMP Inh.) block the signaling cascade by preventing the phosphorylation of the respective SMAD regulators. **B)** Analysis of BMP activation at day 14 by immunofluorescence. **Left:** representative images showing signal of pSMAD1/5/8 in SETX WT, HETERO and HOMO mutant MNs that are positive for the nuclear ISLET1/2 marker (red). **Right:** violin plots showing the intensity of nuclear fluorescence for pSMAD1/5/8, with median values indicated by solid central black line, and interquartile ranges by dashed lines. Data were pooled from 3 independent experiments performed on one SETX WT, one HETERO and one HOMO clones. For each experiment, an average of 100 cells were analyzed per condition. *, P < 0.05; **, P < 0.01; ***, P < 0.001; ****, P < 0.0001 (Mann-Whitney U test, two-tailed). **C)** Analysis of the effect of treatment with TGF-β and BMP inhibitors on the expression of several genes deregulated in L389S mutant MNs at day 14. Plots represent the values of RNA abundance measured by RT-dPCR (reverse transcription and digital PCR) analyses and normalized by the housekeeping *GAPDH* gene. Values correspond to the average and SD of three independent biological replicates. *, P < 0.05; **, P < 0.01; ***, P < 0.001; ****, P < 0.0001 (unpaired t-test, one-tailed). **D)** Analysis of the effect of treatment with TGF-β and BMP inhibitors on axon outgrowth. **Left:** Representative images of MN neurites plated at low density at day 22, immunolabeled for ISL1/2 (red, nuclei) and NEFL (blue, soma and neurites) in the absence or in the presence of TGF-β and BMP inhibitors SB431542 and LDN-193189. Right: violin plots showing the quantification of the longest neurite length at days 14 and 22 for the different genotypes, with median values indicated by solid central black line, and interquartile ranges by dashed lines. Data were pooled from 4 independent experiments performed on two SETX WT (six replicates in total), one HETERO (four replicates in total) and one HOMO (four replicates in total) clones. For each experiment, an average of 25 cells were analyzed per condition. *, P < 0.05; **, P < 0.01; ***, P < 0.001; ****, P < 0.0001 (Mann-Whitney U test, two-tailed). **E)** Working model on the molecular pathways linking SETX gain-of-function L389S mutation to MN degeneration. Based on our results and the known SETX molecular functions, we envision three not mutually-exclusive mechanisms by which SETX ectopic activity could induce mRNA expression and splicing changes: (1) ectopic transcription termination leading to gene repression, more prominent for long genes; (2) aberrant resolution of regulatory R-loops (example of a repressor R-loop); and (3) aberrant or enhanced resolution of R-loops modulating splicing (example of R-loop masking a 3’ SS). These alterations lead to compromised neuronal identity, partially driven by TGF-β aberrant signaling, which result in axonal defects and impaired MN function.

To assess whether the dual treatment with TGF-β and BMP inhibitors can rescue the axonal phenotype of mutant MNs, we measured MNs axon length with and without treatment (figures 7D and S10C). At day 14 we did not observe any major difference between the treated and non-treated cells. However, at day 22 we found that the treatment with the inhibitors supressed the characteristic axon shortening of mutant MNs to a substantial extent.

These results confirm that overactivation of TGF-β superfamily signaling plays a major role in the abnormal gene expression pattern and axonal phenotype of L389S MNs and, therefore, supports the hypothesis that dysregulation of this pathway is a key pathogenic event in ALS4.

## DISCUSSION

In this study, we investigate the molecular cascades through which mutations in the ubiquitously expressed RNA/DNA helicase SETX trigger the selective degeneration of MNs in ALS4. SETX is a highly conserved protein that has been attributed important functions at the interface between gene expression and genome instability. Most studies support roles for SETX in transcription termination and the resolution of R-loops and both activities rely on the catalytic domain of SETX (*62*); however, how they are regulated and coordinated remains unclear.

In close connection with this issue, a central unresolved question is how mutations in SETX alter its different activities and lead to neurodegeneration. SETX is a large protein that contains a superfamily 1 helicase domain (HD), responsible for its catalytic activity, and an N-terminal module involved in protein-protein interactions (Figure S11). In the case of AOA2, a recessive cerebellar ataxia, abundant *in cellulo* experiments, biochemical data and structural modeling indicate that AOA2-associated mutations in SETX lead to partial or complete loss of function by impairing protein expression, stability or catalytic activity, depending on the mutation (*5*). In contrast, ALS4-causing mutations are proposed to induce a toxic gain of function. Indeed, these mutations do not cause symptoms similar to AOA2 and instead affect different regions of the nervous system. Moreover, ectopic expression of an ALS4 mutant version of SETX gene in the presence of WT SETX endogenous alleles is enough to induce MN defects in mouse models (*22*). Further supporting the gain-of-function hypothesis, our ALS4 model does not display phenotypes typically associated with SETX loss of function such as transcription termination/3 ‘end processing defects or the accumulation of R-loops and R-loop-associated DNA damage (Figures S2 and S9).

From a molecular perspective, a former study in fibroblasts reported an interaction between SETX and the zinc finger protein ZPR1, which would be disrupted in cells carrying the ALS4 mutation L389S (*17*), leading the authors to propose that ZPR1 may act as a negative regulator of SETX R-loop-resolving activity. An alternative model emerges from a recent structure-function study of the *Chaetomium thermophilum* Sen1 (ctSen1) homologue of SETX (*63*), together with structural modeling of human SETX (Figure S11). It has been shown that the N-terminal domain (NTD) of ctSen1 mediates an intra-molecular interaction with a subdomain of the HD, the prong, which we have shown to be essential for helicase activity in the budding yeast homologue of SETX (*64*). This interaction locks the protein in a closed, inhibited, conformation, mirroring autoregulatory mechanisms previously described for other helicases. Interactions with regulatory proteins are thought to relieve this autoinhibition, ensuring that helicase activity is deployed at the appropriate time and place (*65*). The most penetrant ALS4 mutations are located at the NTD and HD of SETX. Although most of the NTD is not predicted to adopt stable secondary structure, AlphaFold predicts with relatively high confidence the folding of its extreme N-terminal region, which harbors the L389S substitution, and its proximity to the prong domain, where another penetrant ALS4 mutation (R2136H) is located. It is therefore tempting to speculate that ALS4 mutations could disrupt or destabilize this conserved autoinhibitory interaction resulting in aberrant SETX activity in untimely contexts. This model is not incompatible with the former, as ZPR1 interaction could possibly stabilize SETX autoinhibition, a hypothesis that remains to be investigated.

To shed light onto the molecular mechanisms by which SETX ectopic/aberrant activity leads to the selective degeneration of MNs in ALS4 we have generated an *in vitro* model consisting in iPSCs in which the L389S mutation was introduced into one or both SETX endogenous alleles. Multiple studies have established the validity of iPSC-derived MNs for investigating the molecular basis of MN diseases such as ALS, as this model recapitulates many aspects of human MN biology. Although these cells remain immature compared to adult MNs, they often reproduce pathological phenotypes observed in post-mortem tissues from diseased patients, thereby providing a valuable framework to dissect early pathogenic events (*66*). Compared to models based on patient-derived iPSCs analyzed alongside corrected isogenic controls, the introduction of the disease mutation into a healthy genetic background offers the advantage that observed phenotypes can be more directly attributed to the mutation itself, rather than to a combination of the mutation and patient-specific sensitizing polymorphisms.

The use of this *in vitro* model proved particularly informative in the context of ALS4. While a key ALS4-associated phenotype—namely, upregulation of genes involved in TGF-β signaling—had previously been reported in patient-derived fibroblasts (*16*), our analyses revealed a more complex and rich signature. This included cellular stress, inflammatory responses and partial activation of non-neuronal gene expression programs (e.g. genes involved in cell division, EMT), together with decreased expression of genes involved in neuron-specific functions. These features could not be captured in earlier studies using non-neuronal cell types. As mentioned above, the TGF-β superfamily signaling pathway controls numerous biological processes and plays essential roles in the development and homeostasis of the nervous, cardiovascular and skeletal systems as well as in the modulation of immune responses. Malfunctioning of TGF-β signaling has been linked to a wide range of diseases, including cancer, through its involvement in cell proliferation and the epithelial to mesenchymal transition, cardiac fibrosis, through enhanced inflammatory responses and exacerbated deposition of extracellular matrix components, and neurological disorders such as Alzheimer’s disease and ALS (*27*, *67*, *68*). Together with our metascape network analyses of genes deregulated in ALS4 MNs (figure 2I), these pieces of evidence led us to propose that overactivation of TGF-β signaling is a major driver of the transcriptional changes that we observe in ALS4 MNs. Overall, the gene expression pattern of ALS4 MNs reflects compromised neuronal identity and post-mitotic fate. In line with this, we observe aberrant upregulation of genes associated with cell cycle progression, a phenomenon that in post-mitotic neurons is typically linked to apoptotic cell death (*69*). Consistently, pathways related to apoptosis and p53 signaling are also activated in ALS4 MNs. However, at the time points analyzed, we did not detect a significant reduction in mutant MN survival, suggesting that ALS4 MNs are primed for apoptosis but have not yet fully engaged in cell death programs. Nevertheless, we found that the TGF-β superfamily signaling pathway is indeed overactive in mutant MNs and that dual treatment with TGF-β and BMP signaling inhibitors leads to partial to total suppression of upregulated expression of genes involved in cartilage and bone formation, angiogenesis and EMT (figure 7), providing a causal link between TGF-β increased activity and the aberrant gene expression pattern of ALS4 MNs.

Importantly, some key transcriptomic signatures of ALS4 MNs are shared by other forms of familial ALS (fALS), notably C9ORF72-associated ALS (figure 3), the most common fALS subtype. The core of this common signature includes processes regulated by the TGF-β pathway, consolidating the role of this pathway as a central hub in ALS disease. Although in the SOD1 datasets employed in our analyses the TGF-β signaling pathway did not appear among the enriched upregulated pathways, previous studies using iPSC-derived MNs and mouse models have reported its overactivation in SOD1-associated ALS (*70*, *71*). It is therefore possible that TGF-β dysregulation also contributes to other ALS forms, but that differences in differentiation protocols, sampling time points, and the dynamics of pathway activation may have limited its detection in our analyses. Beyond ALS, compromised neuronal identity has also been reported in Alzheimer’s disease, where aberrant TGF-β signaling has similarly been described (*72*, *73*), suggesting that these events may represent a broader hallmark of neurodegeneration.

Our ALS4 *in vitro* model also enabled us to link TGF-β overactivation to an important pathological phenotype, also observed in other ALS models, consisting in the progressive reduction of axon length. We show that TGF-β dysregulation precedes this phenotype, being detectable at day 14 of differentiation by both transcriptomic analyses (Figures 2 and 3) and immunofluorescence experiments (Figure 7 and S10), whereas axonal defects become apparent only at day 22. Moreover, treatment with TGF-β inhibitors substantially restores axon length at later stages, supporting a causal role for pathway overactivation in axonal impairment. Perturbations in ECM turnover and the activation of mesenchymal and plasticity programs, driven by sustained TGF-β activation, together with reduced expression of genes encoding axonal components may underlie these defects. Actually, we found that decreased expression of the neurofilament component NEFH was also partially suppressed by the treatment with TGF-β inhibitors, supporting a contribution of TGF-β overactivation in downregulation of neuronal genes and further strengthening the link between aberrant TGF-β signaling, compromised neuronal identity and axonal impairment.

Importantly, because the TGF-β pathway is activated in response to injury and is known to promote neuroregeneration, the previously reported evidences of increased TGF-β signaling in ALS4 patient post-mortem tissues could in principle reflect an involvement of this pathway in a compensatory or protective response to neurodegeneration. However, our data instead support the idea that abnormal activation of TGF-β signaling is an early and active driver of pathogenesis. Consistent with this view, previous studies in SOD1 ALS models have shown that TGF-β inhibition alleviates neuroinflammation, motor deficits, and axonal abnormalities, further supporting a pathogenic role for this pathway (*70*, *74*, *75*).

Notably, we did not detect cytoplasmic TDP-43 aggregation in our iPSC-derived ALS4 MNs, despite its reported presence in ALS4 mouse models and patient post-mortem tissues (*22*). This suggests that TDP-43 mislocalization and aggregation are not early pathogenic events but rather later manifestations of disease progression that are not recapitulated in our model, which represents early disease stages. Similarly, we did not observe accumulation of DNA damage under basal conditions, indicating that genome instability is unlikely to be an initiating event in axonal impairment. However, ALS4 MNs display an increased sensitivity to the DNA-damaging agent camptothecin (CPT), pointing to a latent genome instability phenotype that may contribute to disease progression at later stages.

Our new *in vitro* model allowed us not only to elucidate the role of the TGF-β pathway in ALS4 pathology, but also to revisit the current model about the mechanisms underlying overactivation of this pathway in ALS4. As previously mentioned, a study conducted on ALS4 patients fibroblasts proposed that increased TGF-β signaling was due to reduced expression of BAMBI, a transmembrane protein that competes with TGF-β receptors for their cognate ligands, thereby negatively modulating signaling (*16*). However, our transcriptomic data do not show any decrease in the expression of the *BAMBI* gene, but actually a moderate upregulation. This finding indicates that, at least in MNs, an alternative mechanism drives overactivation of TGF-β signaling.

In addition to changes in gene expression, ALS4 MNs exhibited hundreds of splicing alterations (Figure 4). The vast majority consisted of exon skipping or alternative 5’ or 3’ SS usage and were predicted to produce different protein isoforms (figure 4). Genes exhibiting splicing alterations almost do not overlap with those presenting expression changes in mutant MNs, indicating that splicing dysregulation is an independent yet complementary mechanism likely contributing to disease phenotypes. A large fraction of splicing changes followed the same trend in both heterozygous and homozygous clones, although some changes were found only in one of the two mutants. Since heterozygous and homozygous mutants display similar axonal impairment and morphological changes, we reasoned that the contributors to these phenotypes are among the transcripts that present common and robust splicing alterations in both mutants. Indeed, among these common genes, we found several previously associated with neurodegenerative diseases-including other ALS forms and Alzheimer’s disease-suggesting that altered activity of the encoded proteins, due to changes in aminoacid composition, might contribute to ALS4 MN defects.

To understand how ectopic/aberrant SETX activity could produce the observed changes in gene expression and RNA splicing, we searched for features enriched in genes that are deregulated or exhibit alternative splicing. First of all, we found that for both heterozygous and homozygous mutant MNs downregulated genes were considerably longer than genes without significant changes in gene expression (figure 2D). Because SETX interacts-and presumably travels with-transcribing RNAPII (*76*), we hypothesized that the L389S substitution might enable spurious transcription termination by SETX along the gene body. This would lead to the production of truncated, non-functional transcripts that are degraded by RNA quality control pathways, accompanied by decreased levels of full-length transcripts. If ectopic termination events are not frequent, downregulation would be more prominent for long genes. Therefore, aberrant premature transcription termination by L389S SETX might explain part of the gene repression events we observed (Figure 7E).

Beyond transcription termination, we also explored whether aberrant SETX-mediated R-loop resolution could contribute to the observed gene expression dysregulation. R-loops are now recognized as pivotal regulators of gene expression, capable of modulating the binding of regulatory proteins in both positive and negative ways. For instance, at promoter regions R-loops can promote the recruitment of ten-eleven-translocation DNA demethylases and also prevent the association of DNA methyltransferases with the DNA, both of which would result in lower levels of DNA methylation and contribute to active transcription (*7*). Conversely, a subset of DNA regions bound by Polycomb repressive complexes (PRCs) contain R-loops and data in mouse embryonic stem cells and *Drosophila* suggest that R-loops can enhance the recruitment of these proteins, thus promoting gene repression (*52*, *53*). We reasoned that for genes whose expression is controlled by R-loops, these structures would form more frequently or would be more stable to be able to exert robust effects, thus providing stronger signals in R-loop mapping experiments. With this hypothesis in mind, we computed an R-loop propensity score for each expressed gene and found that this score was significantly higher for genes exhibiting changes in expression in ALS4 MNs compared to non-deregulated genes. This result suggests that a fraction of genes could be deregulated in ALS4 mutants due to increased/aberrant removal of R-loops controlling their expression by mutant SETX. As mentioned above, our results support the idea that a large share of the expression changes we detect in mutant MNs are due to overexpression and overactivation of TGF-β pathway genes, and a subset of them have been shown to be repressed by PRCs (*77*). Interestingly, we found that PRC targets, including the *BAMBI* gene, are enriched among genes upregulated in L389S mutant MNs and upregulated PRC targets present a significantly higher R-loop propensity compared to not significantly deregulated genes suggesting that at least some TGF-β genes are upregulated by enhanced removal of R-loops that normally recruit PRCs.

We also found evidence that R-loop abnormal removal could contribute to the splicing alterations detected in ALS4 MNs. Indeed, exons exhibiting alternative splicing in mutant MNs displayed a higher R-loop propensity score than those without splicing alterations, a property that was more prominent for exons exhibiting a higher inclusion rate in mutant MNs (figure 5). Moreover, we found that exons/alternative SS that are more often included/used in the mutants appeared embedded in regions with higher propensity to form R-loops. This suggests that the trapping of SS in RNA:DNA hybrids might preclude their recognition by the spliceosome and that the enhanced resolution of these structures by mutant SETX can favor their exposure to the splicing machinery. For exons/alternative SS that present higher skipping rates in ALS4 MNs we found some local R-loop enrichment at adjacent regions, suggesting a possible role for these R-loops in modulating the recruitment of splicing regulators instead of the core spliceosome (figure 5 and S6).

R-loop removal is likely not the only possible mechanism by which mutant SETX could affect splicing. For instance, SETX translocation along the nascent RNA could potentially interfere with SS recognition or the binding of positive or negative splicing regulators. It will be interesting to assess whether SETX does play a role in splicing under physiological conditions and how disease mutations alter this possible SETX function. Although splicing alterations were previously reported in the context of SETX loss of function (*25*), the profound transcriptomic alterations observed in the same experiments-including deregulation of genes encoding RNA processing factors-complicated the attribution of these splicing changes to a direct action of SETX. Consequently, this aspect remains to be further explored.

Finally, a key remaining question is why are MNs the most vulnerable to the SETX dysfunction caused by ALS4 mutations?. It is known that genes involved in neuron-specific functions are among the longest ones in the genome. Accordingly, genes downregulated in ALS4 MNs are particularly long and gene ontology analyses revealed mainly neuronal-associated terms, such as axon development, synapse, etc. Among all neuronal types, MNs possess exceptionally long axons to innervate very distal parts of the body. This makes them hypersensitive to defects in axonal transport - a process crucial for proper formation and maintenance of axons and synapses (*78*). Remarkably, genes undergoing alternative splicing in ALS4 MNs were enriched in processes that are relevant for axonal transport (e.g. actin and cytoskeleton organization and anchoring junction). In conclusion, the combined reduction in the expression of genes crucial for neuronal functions alongside the potentially altered identity and activity of proteins involved in axonal transport and axon structural integrity might underlie the specific sensitivity of MNs to ALS4 mutations.

Overall, our study shed light onto SETX function in neurons by linking its aberrant activity to coordinated defects in gene expression and RNA processing. Our findings position SETX as a key regulator of neuronal transcriptome integrity and a safeguard of neuronal identity. Our results support a model in which R-loops act as regulatory elements of gene expression whose inappropriate resolution perturbs transcriptional programs, and further suggest that R-loops can directly influence RNA splicing decisions, thereby expanding their functional repertoire in gene regulation.

Importantly, our work also opens new therapeutic perspectives as the rescue of axonal defects by TGF-β inhibition indicates that targeting this pathway could be beneficial in ALS. Because TGF-β inhibitors are already used in cancer therapy, substantial knowledge regarding their administration and safety could be leveraged to design rational therapeutic strategies for ALS4 and potentially other ALS forms.

## MATERIALS AND METHODS

### Human induced pluripotent stem cell (iPSC) lines culture and genome editing

The generation of the SETX L389S iPSC lines, including gRNA/ssODN design, nucleofection, and clonal selection, was performed by the ICV-IPS Cell Culture Platform at the Institut du Cerveau (ICM) (Pitié-Salpêtrière Hospital, Paris, France). The iPSC line WTC11 was used for CRISPR-Cas9 genome editing. Cells were maintained on Laminin-521 (STEMCELL Technologies) or Matrigel (Corning)-coated plates. Culture media consisted of mTeSR Plus (STEMCELL Technologies), supplemented with CloneR or STEMGENT HES Cell Cloning & Recovery Supplement during critical steps such as single-cell recovery and clonal selection. For dissociation, StemPro Accutase (Life technologies) or UltraPure 0.5M EDTA (Invitrogen) were employed depending on the desired grade of dissociation. To introduce the SETX L389S mutation, a single guide RNA (crRNA) and a single-stranded oligodeoxynucleotide (ssODN) donor template were designed using CRISPOR (*79*) and Benchling web tools:

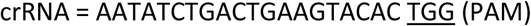

The ssODN template is a 101-bp Ultramer DNA oligo (IDT) containing the target mutation (red) and silent mutations (blue) to prevent re-cutting and facilitate screening= CACGCTGACCACCATTGGCTACGGGGACAAGTACCCCCAGACCTGGAACG**g**CAG**a**CTCCTTGCGGCAACCTTC ACCCTCATCGGTGTCTCCTTCTTCGCGC

Genome editing was performed via ribonucleoprotein (RNP) delivery and Homology-directed repair (HDR) template. Briefly, 1×10^6^ iPSCs were nucleofected using the 4D-Nucleofector System with the P3 Primary Cell Kit (Lonza). The RNP complex consisted of 225 pmol of each RNA (crRNA and tracrRNA-ATTO+) and 120 pmol of Cas9 Nuclease, while HDR templated consisted of 500 pmol of the ssODN repair template supplied with 2 nmol of Electroporation Enhancer. Twenty-four hours post-nucleofection, ATTO-positive cells were isolated via Fluorescence-Activated Cell Sorting (FACS) and plated at a low density (10 cells/cm^2^) on Laminin-521 with CloneR supplement to allow for clonal expansion and selection. After approximately one week, individual clones were mechanically picked and expanded in 96-well plates for downstream molecular analysis and cryopreservation. Genomic DNA was extracted from expanded clones, amplified by PCR using DreamTaq or Platinum Taq DNA Polymerase and specific primers flanking the SETX target region (Table S1) and the presence of the desired mutation was verified by Sanger sequencing. Potential off-target (OT) effects were monitored at four predicted sites using specific primer sets and Sanger sequencing to ensure genomic integrity (Table S1):

OT1 (gene RGS7BP): gATATCTGgCTGAAGaACAC TGG

OT2 (ACO16730.1 chr2:13726000-13729000): AATATaTGgCTGAAaTACAC AGG

OT3 (gene AGPAT9): AATtTCTGAaTGAAaTACAC TGG

OT4 (gene RASSF8): AtaATCTtACTGAAGTACAC AGG

(in green mismatches)

To ensure the maintenance of a normal karyotype following CRISPR manipulation, the edited lines were screened for Copy Number Variations (CNVs) using the iCS-digital™ PSC test (Stem Genomics). This digital PCR-based assay utilized 24 probes covering critical regions of the human genome. Analysis for each clone confirmed a normal diploid status (copy number ∼2) for all tested autosomal loci and a single X chromosome (consistent with the male sex of the line), with no detected CNVs.

### Differentiation of iPSC to MN and pMNs culture

Human iPSC spheroid-based differentiation was performed as previously described (*18*, *19*). iPSCs were dissociated at day 0 with Accutase (Life Technologies) for 3-5 min at 37°C and resuspended in differentiation medium N2B27 [Advanced DMEM F12, Neurobasal vol:vol (Life Technologies) supplemented with N2 (Life Technologies), B27 without Vitamin A (Life Technologies), penicillin/streptomycin 1%, β-mercaptoethanol 0.1% (Life Technologies), Glutamax (Life Technologies)]. Cells were seeded in ultra-low attachment 6 well plates (Corning) at 4×10^5^ cells/ml to form embryoid bodies (EBs). From day 0 to day 4, the medium was supplemented with SB431542 (Stemgent, 20 µM) and LDN-193189 (Stemgent, 0.1 µM) for dual-SMAD inhibition, CHIR-99021 (Selleck, 4 µM) to activate Wnt signaling and Y-27632 (Stem Cell tech, 10 µM, present only for the first 48 hours) to enhance cell survival. On day 4, EBs were patterned toward a ventral spinal fate by replacing the induction factors with Retinoic Acid (RA, Sigma, 100 nM) and Smoothened Agonist (SAG, Merck 500 nM). Providing new media with the same cytokines at day 7, this caudalizing and ventralizing treatment was maintained until day 9 to specify MN progenitors (pMNs). On day 9 EBs were dissociated using 0.05% Trypsin-EDTA (Life Technologies). The trypsin reaction was quenched with FBS, and the cell suspension was further dissociated in cold Neurobasal medium supplemented with DNase I (Roche, 25 mg/mL). After cellular count in a Neubauer chamber, pMNs were frozen in CryoStor® CS10 with a maximum of 2×10^6^ cells/ml per cryovial and stored in liquid nitrogen or used to generate MNs. To promote optimal neuronal attachment and maturation, culture plates were prepared using a two-step coating procedure. Surfaces were first treated with a 0.2% Polyethyleneimine (PEI)-1.4% Borax solution for 2 hours at 37°C as previously shown (*80*). After rinsing abundantly with water, plates were subsequently coated with Mouse Laminin (Life Technologies, 4 µg/mL in F12 medium) for at least 4 hours 37°C or overnight at 4°C. Cryovials were removed from ultra-low temperature storage and immediately immersed in a 37°C water bath until approximately half of the contents were thawed. The cell suspension was then gently transferred into 10 mL of pre-warmed DMEM/F12 medium (Life Technologies). Cells were centrifuged and after supernatant removal, the pellet was resuspended in complete N2B27 maturation medium. Progenitors were seeded at a particular density depending on the specific experiment onto the laminin-coated surfaces (1×10^6^ for nucleic acid extraction, 3.000 cells for axon length and 100.000 cells for the other immunostainings). To enhance initial survival and promote differentiation by Notch signaling inhibition, N2B27 medium was supplemented with Y-27632 (Stem Cell tech, 10 µM), Retinoic Acid (RA, Sigma, 100 nM) and DAPT (Stem Cell tech, 10 µM). At day 11, the medium was refreshed with N2B27 supplemented with Retinoic Acid (RA, Sigma, 100 nM) and DAPT (Stem Cell tech, 10 µM). By day 14, cells were considered post-mitotic MNs, which was verified by immunofluorescence (see below) and maintained in N2B27 medium supplemented till day22 with a cocktail of neurotrophic factors (NTFs), including BDNF, GDNF, CNTF, NT3, HGF and IGF1 (Prepotech, all at 10 ng/mL except for HGF at 2 ng/ml).

### Immunofluorescence experiments

EBs and plated MNs were rinsed with PBS then fixed with 4% PFA for 5 min at 4°C and rinsed with PBS three times for 5 min. EBs were cryoprotected with 30% sucrose and embedded in OCT (Leica) prior to sectioning with a cryostat. Immunostainings were performed as follows: sections were incubated with a saturation solution (PBS, 10% FBS, 0.2% Triton) for 10 min. Primary antibodies (Table S2) were diluted in a staining solution (PBS, 2% FBS, 0.2% Triton) and incubated overnight at 4°C in a humidified chamber. After four PBS washes (10 min each), secondary antibodies (Table S2) were added for 1 h at room temperature. After three PBS washes, DAPI was added to cells (Invitrogen, 1:3000) for 5 min. Cells or slices were then mounted in Fluoromount (SouthernBiotech).

### Cell viability analyses (cMAC)

Cell viability was assessed using the fluorescent thiol-reactive probe blue cMAC (7-amino-4-chloromethylcoumarin, Thermo Fisher Scientific). The dye is designed to freely pass through cell membranes and undergo a glutathione S-transferase-mediated reaction, creating a cell-impermeant fluorescent adduct. Before MNs fixation, the 10 mM stock solution was diluted in N2B27 medium (1:1000). Cells were incubated with the CMAC-containing medium for 35 minutes at 37°C to allow for optimal dye loading and enzymatic conversion. Following incubation, cells were fixed and immunostained for fluorescence microscopy analysis as previously described.

### Image acquisition and analyses

The efficiency of iPSC differentiation into spinal MNs was assessed through a custom CellProfiler pipeline designed for automated cell counting. To ensure the analysis was restricted to viable cells, nuclear segmentation of CMAC-positive cells was performed using the StarDist plugin in Fiji. The resulting objects were subsequently used into CellProfiler and filtered based on morphological dimensions to exclude debris or cell aggregates. The percentage of the viable population expressing neuronal and MN markers (e.g., ISLET1/2, NEFL) was calculated based on fluorescent intensity thresholds. This automated approach provided a quantitative and reproducible validation of the differentiation efficiency across different experimental clones.

To quantify axonal length and body area, MNs were identified by the co-expression of the nuclear marker ISLET1/2 and the cytoskeletal marker NEFL. Semi-automated analysis of neuritic extensions was performed using AutoNeuriteJ (*81*), an ImageJ/Fiji-based plugin. The software is able to segment ISLET1/2-positive soma and NEFL-positive neurites, then after creation of individual neuron images and their skeletization, it allows for the precise measurement and classification of the identified neurites. Final measures include axon length represented by the longest identified neurite, soma area and total axonal tree length.

TDP-43 nucleocytoplasmic distribution was quantified using a custom pipeline in CellProfiler. Nuclear Compartment was defined by the ISLET1/2-positive signal to isolate MN nuclei. Cytoplasmic compartment was identified by the NEFL-positive signal in the immediate proximity of the nucleus. The pipeline measured the integrated intensity of the TDP-43 signal within these two compartments. The results were expressed as the ratio of nuclear/cytoplasmic intensity relative to the total cellular signal.

DNA damage was assessed by quantifying the presence of specific foci (H2AX or 53BP1). Analysis was performed using a custom Fiji (ImageJ) macro. To ensure consistency across samples, a fixed intensity threshold was applied to subtract the background. Foci were identified based on a local increase in pixel intensity above the threshold, and the number of foci per nucleus was automatically recorded. To provide a comprehensive characterization of DNA damage foci we used three complementary approaches: (i) quantification of the absolute number of foci per nucleus to assess total damage; (ii) determination of the percentage of foci-positive cells (defined as cells containing ≥1 focus per nucleus) to evaluate the prevalence of damaged neurons within the population; and (iii) foci distribution analysis across the cell population to identify potential heterogeneities in the DNA damage response under each experimental condition.

Images were acquired at two core platforms. At IGMM, samples were imaged on a Leica THUNDER system using a Leica DFC9000 GT monochrome camera controlled by LAS X software. At IFM, images were gathered with a ZEISS Celldiscoverer 7 system utilizing a ZEISS Axiocam 506 monochrome camera and ZEN software. Hardware settings (excitation energy, gain, and offset) were kept uniform across all groups to enable direct quantitative cross-comparison.

To achieve unbiased population sampling, more than 10 non-overlapping fields were recorded per well across independent samples. Fields were selected automatically via a software-defined grid on the ZEISS Celldiscoverer 7, or randomized manually by systematic navigation across the sample plane on the Leica microscope. Quantitative cell profiling included all eligible targets within every field, yielding the per-well biological averages displayed in the figures.

### Transcriptomic analyses by RNA-seq

Total RNA was extracted from MNs at day 14 (1×10^6^ for each clone). Cells were lysed directly in the culture plate, and RNA was isolated using the NucleoZole RNA Kit (Macherey-Nagel) according to the manufacturer’s instructions. Samples were treated with DNase I using TURBO DNA-*free*™ Kit (Thermo Fisher Scientific) for 20 minutes at 37°C. RNA concentration and purity were assessed using a NanoDrop™ spectrophotometer (Thermo Fisher Scientific). RNA integrity was checked using an Agilent 2100 Bioanalyzer with the Eukaryote Total RNA Nano and Pico kit (Agilent Technologies).

RNA-seq libraries preparation and sequencing was performed by the MGX Platform at the Institut de Génomique Fonctionnelle (Montpellier, France). RNA-seq libraries were prepared using the Stranded Total RNA prep with Ribo-Zero Plus kit (Illumina, San Diego, CA, USA) according to the manufacturer’s instructions. Briefly, an enzymatic depletion removed abundant RNA: Human/mouse/rat cytoplasmic and mtRNA, globin transcripts and bacterial rRNA. Then, depleted RNAs were fragmented using divalent cations at elevated temperature and reverse-transcribed using random hexamers, reverse transcriptase and actinomycin D. Deoxy-TTP was replaced by dUTP during the second strand synthesis to prevent its amplification by PCR. Double stranded cDNAs were adenylated at their 3’ ends and ligated to Illumina’s pre-index anchors. Ligated cDNAs were PCR amplified for 13 cycles with primers including unique dual indexes (UDI) and the PCR products were purified using AMPure XP Beads (Beckman Coulter Genomics, Brea, CA, USA). The size distribution of the resulting libraries was monitored using a Fragment Analyzer (Agilent Technologies, Santa Clara, CA, USA) and the libraries were quantified using the KAPA Library quantification kit (Roche, Basel, Switzerland). The libraries were denatured with NaOH, neutralized with Tris-HCl, and diluted to 150 pM. Clustering and sequencing were performed on a NovaSeq 6000 (Illumina, San Diego, CA, USA) using the paired-end 2×50 nt protocol on 1 lane of a flow cell SP (Illumina sequencing technology). Image analyses and base calling were performed using the NovaSeq Control Software and the Real-Time Analysis component (Illumina). Demultiplexing was performed using Illumina’s conversion software (bcl-convert 4.2.4). The quality of the raw data was assessed using FastQC (0.12.1) from the Babraham Institute (https://www.bioinformatics.babraham.ac.uk/projects/fastqc/) and the Illumina software SAV (Sequencing Analysis Viewer). FastqScreen (0.15.3) was used to identify potential contamination. Adapter sequences were removed, and low-quality bases were trimmed using Cutadapt (version 1.13, Python 3.4.5 *82*).

High-quality reads were aligned to the humain reference genome (GRCh38) using TopHat2 (*83*), and gene-level quantification of mapped reads was performed with HTSeq-count (version 0.6.1p1, (*84*) using the NCBI human genome annotations (release 110).

Differential gene expression analysis was conducted in R using the DESeq package (*85*) using standard parameters and genes for which we did not detect at least 20 reads in at least two samples were filtered out. Volcano plots illustrating significantly differentially expressed genes were generated with a custom R script.

Principal Component Analysis (PCA) was performed on the rLog-normalized count data. To reduce technical noise and prioritize informative features, the analysis was restricted to the 500 genes exhibiting the highest variance across all samples. PCA was computed using the prcomp function in R, with scaling disabled to preserve the homoscedastic properties achieved through the rLog transformation. The relationships between samples were visualized by plotting the first and second principal components (PC1 and PC2).

Heatmap analysis of the expression of specific neuronal and non-neuronal markers across samples was generated using the pheatmap R package and the rLog-normalized count matrix from DESeq analyses as input. We filtered the dataset to include only genes corresponding to our predefined list of cell-type-specific markers.

To assess the relationship between the transcriptomic changes in heterozygous (HETERO) and homozygous (HOMO) clones, the log2 fold change (log2FC) values derived from the respective DESeq analyses were integrated using a custom Rscript. Genes were categorized based on their statistical significance (p < 0.05) and the magnitude of log2FC to identify gene sets that were uniquely or commonly regulated across genotypes. The overall correlation between the HETERO and HOMO log2FC profiles was quantified using the Pearson correlation coefficient. This association was visualized using a scatter plot, where a global linear regression line was fitted to the entire dataset using the lm method to illustrate the general trend of transcriptional behavior across all detected genes, regardless of their individual significance.

### Functional enrichment analyses

Gene Ontology (GO) enrichment analyses were performed using the Gene Ontology Consortium web platform (https://geneontology.org/) with the PANTHER classification system overrepresentation test. Enrichment was assessed across the GO categories biological process, cellular component, and molecular function, as well as Reactome pathways. Statistical significance was determined using Fisher’s exact test with false discovery rate (FDR) correction for multiple testing (Benjamini–Hochberg method). The reference list corresponded to the default background set (all genes in the Homo sapiens genome). Terms with adjusted *p* values below 0.05 were considered significantly enriched. Selected non-overlapping GO terms with a minimal fold enrichment of 3 were selected for representation in main figures using ggplot2 and the full results are provided as supplementary material.

Gene Set Enrichment Analysis (GSEA, *86*) was performed comparing differentially expressed genes in L389S SETX homozygous and L389S heterozygous clones with respect to SETX WT. Genes were ranked using the log2 ratio of classes metric, and statistical significance was estimated through 1,000 permutations of gene sets Enrichment was performed against the Hallmark gene set collection (h.all.v2025.1.Hs.symbols.gmt) from the Molecular Signatures Database (MSigDB v2025.1). This collection was selected to identify well-defined biological states and process patterns. Statistical significance was determined using 1.000 gene set permutations, with a significance threshold of a nominal p-value < 0.05 and an FDR q-value < 0.25. The results were visualized through enrichment plots and heatmaps of the leading-edge genes.

Functional enrichment analysis was performed on genes significantly upregulated (defined by log2FC > 0.9) in L389S SETX homozygous clone against SETX WT. Gene list annotation and enrichment were conducted using Metascape (http://metascape.org) and Homo sapiens as reference species. Enrichment was performed against multiple ontologies, including GO Biological Processes, Canonical Pathways, Reactome Gene Sets, KEGG Pathways, WikiPathways, Panther Pathways and Hallmark Gene Sets. Terms with a p-value <0.01, a minimum count of 3 and an enrichment factor >1.5 were collected and grouped into clusters based on their membership similarities. The resulting network and enrichment clusters were exported and further visualized using Cytoscape (v3.10.4). Cluster partitioning was performed in Cytoscape using the AutoAnnotate app and the MCL (Markov Clustering) algorithm based on membership similarity (Kappa score> 0.3). Descriptive cluster labels were generated via semantic analysis of term occurrence (WordCloud app). In the resulting network, nodes represent enriched terms, while edges represent physical interactions related to shared biological pathways.

Functional annotation of genes encoding transcripts with robust and significant splicing alterations in both HETERO and HOMO mutant MNs was performed using the Database for Annotation, Visualization, and Integrated Discovery (DAVID, *87*) and the GAD_disease segment. The identified associations with neurodegenerative and/or intellectual/psychological disorders were represented by a dot plot using a custom Rscript.

### Differential splicing analyses

Alternative splicing events were quantified using rMATS (version 4.1.2). Aligned reads in BAM format, together with the corresponding GTF annotation file (GRCh38, NCBI release 110), were provided as inputs. rMATS identifies five types of alternative splicing events: skipped exons (SE), alternative 5’ splice sites (A5SS), alternative 3’ splice sites (A3SS), mutually exclusive exons (MXE), and retained introns (RI). Differential splicing between experimental groups was evaluated using only junction reads and a false discovery rate (FDR) ≤0.05 and |ΔPSI| > 0.1 thresholds, and only events supported by at least five reads per replicate were retained. Summary plots were generated with custom R scripts.

To assess the consequences of splicing alterations on resulting protein product and/or transcript sequence, rMATS 0-based start coordinates were converted to 1-based; the coordinate convention was verified empirically against GENCODE v38 exon boundaries (78–94% exact-match rate with the +1 conversion versus <1% without). For each retained event, the predicted protein-sequence consequence was determined by frame-anchored isoform reconstruction against the GENCODE v38 annotation. For the gene overlapping the event locus (strand-aware overlap, preferring the gene whose symbol matched the rMATS annotation), candidate protein-coding transcripts were ranked by biological priority — MANE Select first, then APPRIS principal, then Ensembl canonical, then the longest complete CDS. On the selected transcript, the two isoform forms — exon inclusion (or long form) and exon skipping (or short form) — were reconstructed as spliced transcript sequences, translated in silico from the annotated CDS start, and compared. Retained-intron events were reconstructed by inserting the intron defined by the two flanking splice sites between the two consecutive annotated flanking exons; the rMATS “riExon” was treated as the composite upstream-exon–intron–downstream-exon segment and only the intron boundaries were used. Mutually-exclusive-exon events were reconstructed in two ways: when exactly one of the two exons was present in the anchor transcript, the alternative exon was swapped in on the same backbone (single-anchor swap); when both exons were present as an adjacent tandem pair, the two isoforms were reconstructed as backbone-minus-second-exon (1st-exon form) and backbone-minus-first-exon (2nd-exon form). Events were classified as: « in-frame indel » (the alternative segment length is divisible by 3 and introduces no stop codon (net in-frame insertion or deletion); « PTC » (a premature termination codon arising earlier than the reference stop, subdivided into in-frame stops within the segment and frameshift-driven stops); « frameshift without PTC » (a frameshift producing a C-terminal extension or no in-frame stop; « stop_lost / start_lost » (removal of the reference stop or start codon); « exon swap » (substitution); « UTR or intronic » (the alternative segment lies outside the CDS); « complex/other » (not able to clasify within former categories).

For MXE events both isoforms are real, expressed products, a premature stop was therefore called on whichever form terminated early, and an in-frame swap that changes sequence without changing length was classified as an exon swap (substitution) rather than a PTC.

PTC-bearing transcripts were evaluated for nonsense-mediated decay (NMD) sensitivity using the >55-nt rule: a stop codon located more than 55 nt upstream of the last exon–exon junction of the same transcript was classified as an NMD candidate.

Events with no matching annotated isoform and those whose rMATS flanking exons were discordant with the chosen transcript were re-analysed with an annotation-light, flank-anchored method. The reading frame was anchored from the event’s own rMATS flanking exons rather than the full canonical transcript: for each flanking splice boundary, all annotated coding exons sharing that boundary supplied the reading-frame phase relative to their CDS start. The segment and the skipping product were then translated in the anchored frame to flag poison exons (an in-frame stop within the segment) and to localise the PTC on the frameshifted isoform.

To analyse the GC content of exons with and without differential splicing in ALS4 MNs, exonic and flanking intronic sequences corresponding to the hg38 human reference genome were extracted the using R packages GenomicRanges and BSgenome.Hsapiens.UCSC.hg38. Nucleotide frequencies (A, C, G, T) across target regions were quantified using the Biostrings package and the percentage GC content was calculated for: exon body (i.e. the entire target exon); 3’SS region (-50 bp intron to +15 bp exon); and 5’SS region (-15 bp intron to +50 bp exon). GC contrast was computed as the difference in GC content between the exon body and the average of its immediate flanking introns (100 bp upstream and downstream). Group distributions and median values across categories were visualized using combined violin and box plots generated with ggplot2 and ggpubr.

To compute the SS strength for exons presenting differential splicing in L389S mutants genomic sequences surrounding the splice junctions were extracted from the human reference genome (GRCh38/hg38) using GenomicRanges and BSgenome: the 5ʹSS was extracted as a 9-bp sequence spanning positions-3 to +6 relative to the 5ʹ exon-intron junction, while the 3ʹSS was extracted as a 23-bp sequence spanning positions-20 to +3 relative to the intron-exon 3ʹ junction. The 5’SS strength was calculated via a Position Weight Matrix (PWM) log-likelihood scoring approach relative to uniform background nucleotide frequencies (0.25), incorporating a pseudo-count of 0.01 to handle zero probabilities, as previously described (*88*). The 3’SS strength was computed using an empirical score combining the presence of an invariant 3ʹ canonical AG dinucleotide with the pyrimidine count (C/T) across the upstream 18-bp polypyrimidine tract.

Pairwise statistical comparisons of splice site scores between altered exon subsets (“all altered”, “higher inclusion”, and “more skipped”) and the “not altered” reference group were evaluated using two-sided Wilcoxon rank-sum tests. Data distributions, medians, and sample counts were visualized using combined violin, boxplot, and individual jitter plots.

### Meta-analyses of ALS datasets

To search for common trancriptomic signatures between previously reported datasets generated from iPSC-derived MNs carrying ALS-mutations and ALS4 datasets generated in the present study, we first normalized all datasets by Z-score transformation using the scale function in R. To resolve tied ranks and ensure numerical stability during permutations, a negligible amount of Gaussian noise (SD = 10^-10^) was added to the scores. Gene set enrichment analysis was performed using the fgsea R package (v1.28.0) against the MsigDB Hallmark Gene Set collection (v2023.2). The analysis utilized a minimum gene set size of 5 and a maximum of 500.

To account for multiple hypothesis testing across the 50 Hallmark pathways, p-values were adjusted using the Benjamini-Hochberg (BH) procedure to control the False Discovery Rate (FDR). Pathways were considered statistically significant only if the adjusted p-value (padj) was less than 0.05.

Significant pathways were aggregated into a master Normalized Enrichment Score (NES) matrix. To maintain a comprehensive comparison, if a pathway reached significance in at least one condition, its NES values were extracted for all other conditions. The final results were visualized as a heatmap using the pheatmap package, employing Euclidean distance clustering to group datasets with similar enrichment profiles. To identify conserved SETX-related signatures, pathways were filtered for those exhibiting significant enrichment (padj <0.1) and a Normalized Enrichment Score (NES) > 1.2 in both SETX L389S heterozygous and homozygous datasets.

To compare genes deregulated in our ALS4 model with expression changes in available transcriptomic datasets from post-mortem tissues of ALS patients, we first retrieved the GSE56500 (3 sporadic ALS patients and 6 healthy controls), GSE18920 (12 sporadic ALS and 3 controls), and GSE68605 (8 C9ORF72 ALS and 3 controls) datasets using the *GEOquery* R package. Their expression profiles underwent log2 transformation and quality filtering. Differential expression analysis was performed across using the *limma* package by fitting a linear model to compare ALS samples against controls. Statistical significance was determined using moderated t-statistics via the Empirical Bayes method, and p-values were adjusted for multiple testing using the False Discovery Rate (FDR). The resulting processed data were compared with the DESeq2 result table of the L389S HOMO mutant (see details above). The overlapping gene sets among the four populations were visualized using Venn diagrams. To determine whether the observed overlaps were greater (or smaller) than expected by chance, a Monte Carlo resampling approach was employed to construct empirical null distributions.

For each population, the observed sample size (n_1, n_2, n_3, n_4) was recorded. At each iteration of the simulation (K = 10,000), four random samples of corresponding sizes were independently drawn from the reference universe of all possible elements. For every possible intersection—including pairwise, three-way, and four-way intersections—the number of shared elements was computed to generate an empirical null distribution. The observed overlap size (O_obs_) was then compared to this distribution, and an empirical p-value was calculated using the formula:

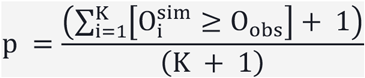

where O_i_^sim^ represents the overlap size at iteration i. This calculation incorporates the correction for finite permutations (*89*). Finally, p-values were adjusted using the Benjamini–Hochberg procedure to control the FDR across all simultaneous tests. This null hypothesis assumes that population membership is independent, meaning deviations reflect non-random co-occurrence or mutual exclusivity of elements. All intersecting gene sets appeared highly significant (p <10^-4^).

### Global quantification of cellular R-loops by slot blot

Genomic DNA was extracted from MNs at day 14 (1×10^6^ for each clone) using a manual phenol-chloroform protocol optimized for R-loop preservation. Cells were lysed in lysis buffer (10 mM Tris, 20 mM EDTA, 1% SDS) and incubated at 50°C with Proteinase K (Euromedex, 500 μg/mL). To ensure high purity and minimize shearing, the lysate was processed using Phase Lock Gel tubes (Qiagen). An equal volume of Phenol:Chloroform:Isoamyl Alcohol (25:24:1) was added, and the aqueous phase was separated by centrifugation. DNA was then precipitated with ethanol 100% and sodium acetate (0.3M final, pH5.2) in the presence of glycogen blue (Life technologies), washed with 80% ethanol and resuspended in low-EDTA TE buffer (10 mM Tris, 0.5 mM EDTA). Genomic DNA was fragmented by an overnight (ON) digestion at 37°C using a Restriction Enzyme Cocktail (New England Biolabs, EcoRI-HF and HindIII-HF). Then genomic DNA was treated for 2 hours at 37°C with RNase III (Ambion, AM2290), to avoid S9.6 antibody interference with dsRNA, and with RNase H (New England Biolabs, M0297L), as negative control specific for R-loops. DNA concentration was quantified using the Qubit™ dsDNA BR (Broad Range) Assay Kit with a Qubit™ Fluorometer (Thermo Fisher Scientific). Two dilutions of each sample were loaded on nylon membrane (GE Healthcare, Amersham Hybond-N+) using a vacuum slot blot apparatus (Bio-Dot® and Bio-Dot SF Microfiltration Apparatus). After loading, the DNA was fixed to the membrane via UV cross-linking (2 times with 0.12 J). The membrane was immediately blocked with 5% milk in PBS-Tween 0.02% and incubated overnight at 4°C with the S9.6 antibody (1μg/mL final, Table S2). R-loops detection was performed by chemiluminescence using SuperSignal™ West Femto (ThermoFisher). For loading control, the membrane was subsequently incubated with an anti-dsDNA antibody (1:1000, Table S2) and revealed with SuperSignal™ Pico Plus (ThermoFisher). Signals were quantified using Fiji.

### Polycomb repressive complexes target genes enrichment analyses

Precomputed target-gene tables for core PRC2 (EZH2, SUZ12, EED) and PRC1 (RNF2, BMI1) subunits were obtained from ChIP-Atlas (https://chip-atlas.org; *90*) for the hg38 assembly based on binding events within TSS ±5kb. For each factor, genes were ranked by their average MACS2 binding score, and the top 2,000 targets were selected. To restrict targets to a neural context, gene promoters were filtered by intersecting with replicated H3K27me3 ChIP-seq narrowPeak files obtained from ENCODE (the PRC1 mark H2AK119ub is not available in ENCODE and could not be assayed). A neural-union reference was constructed across 10 diverse neural biosamples (brain, dorsolateral prefrontal cortex, neural progenitor cell, neuronal stem cell, neural cell, bipolar neuron, mid-neurogenesis radial glial cells, NT2/D1, SK-N-SH, and neural crest cell). A promoter was categorized as marked if it overlapped ≥ 1 peak within any constituent neural reference. Strand-aware gene models and coordinates were established using GENCODE GRCh38.113 annotations.

The lists of upregulated genes in L389S mutants (« common » and HOMO) were tested for enrichment against each neural Polycomb target set. Significance and odds ratios were evaluated using a one-sided Fisher’s exact test (“greater”) against a background universe of all expressed/tested genes (n = 18,527). *p*-values were adjusted for multiple testing using the Benjamini–Hochberg method.

### R-loop datasets and analyses

To compute an R-loop propensity score for each expressed gene we used available DRIP-seq datasets generated in HEK293 together with their corresponding RNA-seq data (DESeq2 normalized counts file) performed in the same cells and conditions (GSE102474, *49*). DRIP-seq datasets from iPSC-derived neurons (GSE324290, *51*) were used to verify signal robustness et reproducibility across cell lines.

R-loop signal was quantified and integrated with gene expression data using custom scripts in R (version ≥4.0) with packages from Bioconductor, including rtracklayer and GenomicRanges, as well as dplyr, data.table, ggplot2, and ggpubr. Gene/exon coordinates were obtained from a GRCh38 GTF annotation file. Total R-loop signal at gene body or exon was derived from DRIP-seq bigwig files by summing per-base signal intensities across each region using genomic overlap operations. The total R-loop signal per region was normalized by region length (base pairs) to obtain a length-adjusted R-loop density. A transcription-normalized R-loop metric was calculated by dividing the length-normalized R-loop signal by the corresponding average RNA-seq expression value. Genes with missing or zero RNA-seq expression values were excluded. Genes/exons were stratified into the different categories to be compared. For visualization, the transcription-normalized R-loop signal was scaled (×1000) and log10-transformed, with a small offset (10⁻³) added prior to transformation to avoid undefined values.

Data were represented using ggplot2 and statistical comparisons between groups were performed using non-parametric tests: pairwise comparisons were conducted using the Wilcoxon rank-sum test

Metagene analyses of normalized R-loop signal at exons differentially spliced in L389S mutants and flanking regions were performed in R using packages from the Bioconductor project and CRAN. R-loop signal density across these genomic intervals was profiled using DRIP-seq bigwig data imported via the rtracklayer package. Strand orientation and genomic coordinate manipulations were handled using GenomicRanges. For each exon, signal values were extracted across a 500 bp upstream flank (divided into 50 bins), a scaled 200 bp exon body (divided into 20 bins), and a 500 bp downstream flank (divided into 50 bins). To control for transcription differences, the extracted DRIP-seq coverage was normalized against average RNA-seq expression counts. Mean metagene profiles and standard errors of the mean (SEM) were computed for each splicing category and plotted using ggplot2 and ggpubr.

### DNA damage induction

To induce DNA damage, MNs at days 14 and 22 were treated with Camptothecin (CPT, Euromedex). The Topoisomerase I inhibitor was prepared as a 20 mM stock solution and diluted 2000X in N2B27 medium for 3 hours at 37°C. Following treatment, cells were immediately processed for fixation and immunostaining to analyze DNA damage foci formation.

### TGF-β and BMP inhibition

TGF-β signaling was inhibited from the progenitor stage (day 9) to target a pre-symptomatic developmental window. Upon thawing, treated MN progenitors were cultured in N2B27 medium supplemented with the dual-SMAD inhibitors SB431542 (Stemgent 20 µM) and LDN-193189 (Stemgent 0.1 µM), which block the kinase activity of ALK4/5/7 receptors and ALK1/2/3/6 receptors, respectively. This treatment was maintained for every change of the media till day 22.

### Reverse Transcription and Quantitative PCR (RT-qPCR)

Total RNA (150 ng) was reverse-transcribed using M-MLV reverse transcriptase (200 U/µL; New England BioLabs) with 10 mM oligo(dT) and 50 ng/µL random hexamers at 37°C for 45 min. No-RT control reactions (without enzyme) were included for each sample. cDNA was stored at −20°C.

Quantitative PCR was performed using the LightCycler 480 SYBR Green I Master reagent (2X; Roche, REF04887352001) and a LightCycler 480 instrument in 384-well plates (Roche, REF04729749001) with 2 µL of cDNA and 5 µM of specific primers per reaction. Thermal cycling consisted of an initial denaturation at 95°C for 5 min, followed by 50 cycles of 95°C for 15 s, 55°C for 15 s, and 72°C for 20 s, concluding with a melting curve analysis (65°C to 95°C) and a cooling step at 40°C for 30 s. Absolute quantification and reaction efficiency were determined using the LightCycler 480 SW software (v1.5.1.62) against a genomic DNA standard curve. mRNA expression values were normalized to the housekeeping gene *GAPDH*.

### Digital PCR (dPCR)

Transcripts for which variability was too high with standard qPCR were quantified by dPCR on the QIAcuity One platform (QIAGEN) using EvaGreen-based detection. Reactions (40 µL total volume) contained 13.3 µL of 3X EvaGreen PCR Master Mix, 0.4 µM of each forward and reverse primer, and 2 µL of cDNA (diluted based on prior qPCR expression levels). No-template controls (NTCs) replaced cDNA with nuclease-free water.

Reactions were loaded onto QIAcuity Nanoplate 26K plates, sealed with specialized foil, and primed using the manufacturer’s protocol. Amplification conditions comprised an initial polymerase activation at 95°C for 2 min, followed by 40 cycles of 95°C for 15 s, 58°C for 15 s, and 72°C for 15 s, with a final cooling step at 40°C for 5 min. Imaging was acquired via the Green channel (200 ms exposure, gain setting 2).

Data were analyzed using the Absolute Quantification module of the QIAcuity Software Suite (v2.5.0.1; QIAGEN), with thresholds set relative to NTC signals. Target concentrations (copies/µL) were converted to ng/µL using amplicon length, Avogadro’s number, and an average molecular weight of 660 Da/bp. Relative expression levels were calculated by normalizing target concentrations to *GAPDH*. Primers for qPCR and dPCR are listed in table S1.

## Funding

Fondation pour la Recherche Médicale grant MND202004011479 (S.N., O.P)

French National Research Agency grant ANR-24-CE12-1756 (S.N., O.P.)

Institut National de la santé et de la recherche médicale (S.N.)

Centre National de la Recherche Scientifique (T.G., A.Z.A, O.P)

## Author contributions

Conceptualization: O.P.

Methodology: M.G., T.G., S.N., O.P.

Experimentation: M.G., C.B.

Data analysis: M.G, T.G, A.Z.A, O.P.

Supervision: O.P.

Writing—original draft: M.G., O.P.

Writing—review & editing: S.N., T.G., O.P.

## Other contributions

We thank all the members of the O.P lab for insightful discussions and Marie Antoine for lab management and help in iPSC and MN culture. We thank Simona Gribaudo and Maeliss Calon for training M.G. in iPSC culture and differenciation into MNs, Nicolas Jardin for developing the CMAC pipeline of analysis and Vincent Vanoosthuyse for providing protocols and advice on R-loop analyses by slot blot. We acknowledge Stephanie Bigou for genetic modification of iPSC lines. Genome editing was carried out at the Paris Brain Institute iPSC core facility (RRID:SCR_027891), who has received funding from the program “Investissements d’avenir” ANR-10-IAIHU-06 and is part of the DIM C-BRAINS, funded by the Conseil Régional d’Ile-de-France. MGX acknowledges financial support from France Génomique National Infrastructure, funded as part of the ‘Investissement d’Avenir’ program managed by the Agence Nationale pour la Recherche (ANR-10-INBS-09). We acknowledge the imaging facility MRI, member of the national infrastructure France-BioImaging (https://ror.org/01y7vt929) supported by the French National Research Agency (ANR-24-INBS-0005 FBI BIOGEN).

## REFERENCES

1. Z. Butti, S. A. Patten, RNA Dysregulation in Amyotrophic Lateral Sclerosis. Front. Genet. 9 (2019).

2. J. K. Nussbacher, R. Tabet, G. W. Yeo, C. Lagier-Tourenne, Disruption of RNA Metabolism in Neurological Diseases and Emerging Therapeutic Interventions. Neuron 102, 294–320 (2019).

3. J. R. Kok, N. M. Palminha, C. Dos Santos Souza, S. F. El-Khamisy, L. Ferraiuolo, DNA damage as a mechanism of neurodegeneration in ALS and a contributor to astrocyte toxicity. Cell. Mol. Life Sci. 78, 5707–5729 (2021).

4. C. Beghѐ, H. Harpham, Y. Barberic, N. Gromak, R-loops in neurodegeneration. Curr Opin Genet Dev 92, 102345 (2025).

5. M. Giannini, O. Porrua, Senataxin: A key actor in RNA metabolism, genome integrity and neurodegeneration. Biochimie, S0300-9084(23)00184–0 (2023).

6. J. R. Brickner, J. L. Garzon, K. A. Cimprich, Walking a tightrope: The complex balancing act of R-loops in genome stability. Mol Cell 82, 2267–2297 (2022).

7. C. Niehrs, B. Luke, Regulatory R-loops as facilitators of gene expression and genome stability. Nat Rev Mol Cell Biol 21, 167–178 (2020).

8. M.-C. Moreira, S. Klur, M. Watanabe, A. H. Németh, I. Le Ber, J.-C. Moniz, C. Tranchant, P. Aubourg, M. Tazir, L. Schöls, M. Pandolfo, J. B. Schulz, J. Pouget, P. Calvas, M. Shizuka-Ikeda, M. Shoji, M. Tanaka, L. Izatt, C. E. Shaw, A. M’Zahem, E. Dunne, P. Bomont, T. Benhassine, N. Bouslam, G. Stevanin, A. Brice, J. Guimarães, P. Mendonça, C. Barbot, P. Coutinho, J. Sequeiros, A. Dürr, J.-M. Warter, M. Koenig, Senataxin, the ortholog of a yeast RNA helicase, is mutant in ataxia-ocular apraxia 2. Nat. Genet. 36, 225–227 (2004).

9. Y.-Z. Chen, C. L. Bennett, H. M. Huynh, I. P. Blair, I. Puls, J. Irobi, I. Dierick, A. Abel, M. L. Kennerson, B. A. Rabin, G. A. Nicholson, M. Auer-Grumbach, K. Wagner, P. De Jonghe, J. W. Griffin, K. H. Fischbeck, V. Timmerman, D. R. Cornblath, P. F. Chance, DNA/RNA helicase gene mutations in a form of juvenile amyotrophic lateral sclerosis (ALS4). Am. J. Hum. Genet. 74, 1128–1135 (2004).

10. F. Akçimen, E. R. Lopez, J. E. Landers, A. Nath, A. Chiò, R. Chia, B. J. Traynor, Amyotrophic lateral sclerosis: translating genetic discoveries into therapies. Nat Rev Genet 24, 642–658 (2023).

11. N. Suzuki, A. Nishiyama, H. Warita, M. Aoki, Genetics of amyotrophic lateral sclerosis: seeking therapeutic targets in the era of gene therapy. J Hum Genet 68, 131–152 (2023).

12. C. Walker, S. Herranz-Martin, E. Karyka, C. Liao, K. Lewis, W. Elsayed, V. Lukashchuk, S.-C. Chiang, S. Ray, P. J. Mulcahy, M. Jurga, I. Tsagakis, T. Iannitti, J. Chandran, I. Coldicott, K. J. De Vos, M. K. Hassan, A. Higginbottom, P. J. Shaw, G. M. Hautbergue, M. Azzouz, S. F. El-Khamisy, C9orf72 expansion disrupts ATM-mediated chromosomal break repair. Nat Neurosci 20, 1225–1235 (2017).

13. M. Giannini, A. Bayona-Feliu, D. Sproviero, S. I. Barroso, C. Cereda, A. Aguilera, TDP-43 mutations link Amyotrophic Lateral Sclerosis with R-loop homeostasis and R loop-mediated DNA damage. PLoS Genet 16, e1009260 (2020).

14. M. Wood, A. Quinet, Y.-L. Lin, A. A. Davis, P. Pasero, Y. M. Ayala, A. Vindigni, TDP-43 dysfunction results in R-loop accumulation and DNA replication defects. Journal of Cell Science 133, jcs244129 (2020).

15. V. F. Thompson, D. R. Wieland, V. Mendoza-Leon, H. I. Janis, M. A. Lay, L. M. Harrell, J. C. Schwartz, Binding of the nuclear ribonucleoprotein family member FUS to RNA prevents R-loop RNA:DNA hybrid structures. Journal of Biological Chemistry 299 (2023).

16. C. Grunseich, I. X. Wang, J. A. Watts, J. T. Burdick, R. D. Guber, Z. Zhu, A. Bruzel, T. Lanman, K. Chen, A. B. Schindler, N. Edwards, A. Ray-Chaudhury, J. Yao, T. Lehky, G. Piszczek, B. Crain, K. H. Fischbeck, V. G. Cheung, Senataxin Mutation Reveals How R-Loops Promote Transcription by Blocking DNA Methylation at Gene Promoters. Molecular Cell 69, 426–437.e7 (2018).

17. A. Kannan, J. Cuartas, P. Gangwani, D. Branzei, L. Gangwani, Mutation in senataxin alters the mechanism of R-loop resolution in amyotrophic lateral sclerosis 4. Brain 145, 3072–3094 (2022).

18. Y. Maury, J. Côme, R. A. Piskorowski, N. Salah-Mohellibi, V. Chevaleyre, M. Peschanski, C. Martinat, S. Nedelec, Combinatorial analysis of developmental cues efficiently converts human pluripotent stem cells into multiple neuronal subtypes. Nat. Biotechnol. 33, 89–96 (2015).

19. V. Mouilleau, C. Vaslin, R. Robert, S. Gribaudo, N. Nicolas, M. Jarrige, A. Terray, L. Lesueur, M. W. Mathis, G. Croft, M. Daynac, V. Rouiller-Fabre, H. Wichterle, V. Ribes, C. Martinat, S. Nedelec, Dynamic extrinsic pacing of the HOX clock in human axial progenitors controls motor neuron subtype specification. Development 148, dev194514 (2021).

20. F. Tian, W. Yang, D. A. Mordes, J.-Y. Wang, J. S. Salameh, J. Mok, J. Chew, A. Sharma, E. Leno-Duran, S. Suzuki-Uematsu, N. Suzuki, S. S. Han, F.-K. Lu, M. Ji, R. Zhang, Y. Liu, J. Strominger, N. A. Shneider, L. Petrucelli, X. S. Xie, K. Eggan, Monitoring peripheral nerve degeneration in ALS by label-free stimulated Raman scattering imaging. Nat Commun 7, 13283 (2016).

21. N. Egawa, S. Kitaoka, K. Tsukita, M. Naitoh, K. Takahashi, T. Yamamoto, F. Adachi, T. Kondo, K. Okita, I. Asaka, T. Aoi, A. Watanabe, Y. Yamada, A. Morizane, J. Takahashi, T. Ayaki, H. Ito, K. Yoshikawa, S. Yamawaki, S. Suzuki, D. Watanabe, H. Hioki, T. Kaneko, K. Makioka, K. Okamoto, H. Takuma, A. Tamaoka, K. Hasegawa, T. Nonaka, M. Hasegawa, A. Kawata, M. Yoshida, T. Nakahata, R. Takahashi, M. C. N. Marchetto, F. H. Gage, S. Yamanaka, H. Inoue, Drug screening for ALS using patient-specific induced pluripotent stem cells. Sci Transl Med 4, 145ra104 (2012).

22. C. L. Bennett, S. G. Dastidar, S.-C. Ling, B. Malik, T. Ashe, M. Wadhwa, D. B. Miller, C. Lee, M. B. Mitchell, M. A. van Es, C. Grunseich, Y. Chen, B. L. Sopher, L. Greensmith, D. W. Cleveland, A. R. La Spada, Senataxin mutations elicit motor neuron degeneration phenotypes and yield TDP-43 mislocalization in ALS4 mice and human patients. Acta Neuropathol. 136, 425–443 (2018).

23. B. A. Berning, A. K. Walker, The Pathobiology of TDP-43 C-Terminal Fragments in ALS and FTLD. Front. Neurosci. 13 (2019).

24. A. Prasad, V. Bharathi, V. Sivalingam, A. Girdhar, B. K. Patel, Molecular Mechanisms of TDP-43 Misfolding and Pathology in Amyotrophic Lateral Sclerosis. Front Mol Neurosci 12, 25 (2019).

25. P. Richard, S. Feng, Y.-L. Tsai, W. Li, P. Rinchetti, U. Muhith, J. Irizarry-Cole, K. Stolz, L. A. Sanz, S. Hartono, M. Hoque, S. Tadesse, H. Seitz, F. Lotti, M. Hirano, F. Chédin, B. Tian, J. L. Manley, SETX (senataxin), the helicase mutated in AOA2 and ALS4, functions in autophagy regulation. Autophagy 17, 1889–1906 (2021).

26. I. Lopes, G. Altab, P. Raina, J. P. de Magalhães, Gene Size Matters: An Analysis of Gene Length in the Human Genome. Front Genet 12, 559998 (2021).

27. J. Massagué, D. Sheppard, TGF-β signaling in health and disease. Cell 186, 4007–4037 (2023).

28. A. Blokzijl, C. Dahlqvist, E. Reissmann, A. Falk, A. Moliner, U. Lendahl, C. F. Ibáñez, Cross-talk between the Notch and TGF-β signaling pathways mediated by interaction of the Notch intracellular domain with Smad3. J Cell Biol 163, 723–728 (2003).

29. Y. Ito, K. Miyazono, RUNX transcription factors as key targets of TGF-β superfamily signaling. Current Opinion in Genetics & Development 13, 43–47 (2003).

30. Y. Hao, D. Baker, P. ten Dijke, TGF-β-Mediated Epithelial-Mesenchymal Transition and Cancer Metastasis. Int J Mol Sci 20, 2767 (2019).

31. O. J. Ziff, J. Neeves, J. Mitchell, G. Tyzack, C. Martinez-Ruiz, R. Luisier, A. M. Chakrabarti, N. McGranahan, K. Litchfield, S. J. Boulton, A. Al-Chalabi, G. Kelly, J. Humphrey, R. Patani, Integrated transcriptome landscape of ALS identifies genome instability linked to TDP-43 pathology. Nat Commun 14, 2176 (2023).

32. R. Derynck, Y. E. Zhang, Smad-dependent and Smad-independent pathways in TGF-β family signalling. Nature 425, 577–584 (2003).

33. D. Sareen, J. G. O’Rourke, P. Meera, A. K. M. G. Muhammad, S. Grant, M. Simpkinson, S. Bell, S. Carmona, L. Ornelas, A. Sahabian, T. Gendron, L. Petrucelli, M. Baughn, J. Ravits, M. B. Harms, F. Rigo, C. F. Bennett, T. S. Otis, C. N. Svendsen, R. H. Baloh, Targeting RNA foci in iPSC-derived motor neurons from ALS patients with a C9ORF72 repeat expansion. Sci Transl Med 5, 208ra149 (2013).

34. M. Abo-Rady, N. Kalmbach, A. Pal, C. Schludi, A. Janosch, T. Richter, P. Freitag, M. Bickle, A.-K. Kahlert, S. Petri, S. Stefanov, H. Glass, S. Staege, W. Just, R. Bhatnagar, D. Edbauer, A. Hermann, F. Wegner, J. L. Sterneckert, Knocking out C9ORF72 Exacerbates Axonal Trafficking Defects Associated with Hexanucleotide Repeat Expansion and Reduces Levels of Heat Shock Proteins. Stem Cell Reports 14, 390–405 (2020).

35. R. Dafinca, P. Barbagallo, L. Farrimond, A. Candalija, J. Scaber, N. A. Ababneh, C. Sathyaprakash, J. Vowles, S. A. Cowley, K. Talbot, Impairment of Mitochondrial Calcium Buffering Links Mutations in C9ORF72 and TARDBP in iPS-Derived Motor Neurons from Patients with ALS/FTD. Stem Cell Reports 14, 892–908 (2020).

36. A. Catanese, S. Rajkumar, D. Sommer, D. Freisem, A. Wirth, A. Aly, D. Massa-López, A. Olivieri, F. Torelli, V. Ioannidis, J. Lipecka, I. C. Guerrera, D. Zytnicki, A. Ludolph, E. Kabashi, M. A. Mulaw, F. Roselli, T. M. Böckers, Synaptic disruption and CREB-regulated transcription are restored by K+ channel blockers in ALS. EMBO Mol Med 13, e13131 (2021).

37. D. Sommer, S. Rajkumar, M. Seidel, A. Aly, A. Ludolph, R. Ho, T. M. Boeckers, A. Catanese, Aging-Dependent Altered Transcriptional Programs Underlie Activity Impairments in Human C9orf72-Mutant Motor Neurons. Front Mol Neurosci 15, 894230 (2022).

38. NeuroLINCS Consortium, J. Li, R. G. Lim, J. A. Kaye, V. Dardov, A. N. Coyne, J. Wu, P. Milani, A. Cheng, T. G. Thompson, L. Ornelas, A. Frank, M. Adam, M. G. Banuelos, M. Casale, V. Cox, R. Escalante-Chong, J. G. Daigle, E. Gomez, L. Hayes, R. Holewenski, S. Lei, A. Lenail, L. Lima, B. Mandefro, A. Matlock, L. Panther, N. L. Patel-Murray, J. Pham, D. Ramamoorthy, K. Sachs, B. Shelley, J. Stocksdale, H. Trost, M. Wilhelm, V. Venkatraman, B. T. Wassie, S. Wyman, S. Yang, NYGC ALS Consortium, J. E. Van Eyk, T. E. Lloyd, S. Finkbeiner, E. Fraenkel, J. D. Rothstein, D. Sareen, C. N. Svendsen, L. M. Thompson, An integrated multi-omic analysis of iPSC-derived motor neurons from C9ORF72 ALS patients. iScience 24, 103221 (2021).

39. E. G. Baxi, T. Thompson, J. Li, J. A. Kaye, R. G. Lim, J. Wu, D. Ramamoorthy, L. Lima, V. Vaibhav, A. Matlock, A. Frank, A. N. Coyne, B. Landin, L. Ornelas, E. Mosmiller, S. Thrower, S. M. Farr, L. Panther, E. Gomez, E. Galvez, D. Perez, I. Meepe, S. Lei, B. Mandefro, H. Trost, L. Pinedo, M. G. Banuelos, C. Liu, R. Moran, V. Garcia, M. Workman, R. Ho, S. Wyman, J. Roggenbuck, M. B. Harms, J. Stocksdale, R. Miramontes, K. Wang, V. Venkatraman, R. Holewenski, N. Sundararaman, R. Pandey, D.-M. Manalo, A. Donde, N. Huynh, M. Adam, B. T. Wassie, E. Vertudes, N. Amirani, K. Raja, R. Thomas, L. Hayes, A. Lenail, A. Cerezo, S. Luppino, A. Farrar, L. Pothier, C. Prina, T. Morgan, A. Jamil, S. Heintzman, J. Jockel-Balsarotti, E. Karanja, J. Markway, M. McCallum, B. Joslin, D. Alibazoglu, S. Kolb, S. Ajroud-Driss, R. Baloh, D. Heitzman, T. Miller, J. D. Glass, N. L. Patel-Murray, H. Yu, E. Sinani, P. Vigneswaran, A. V. Sherman, O. Ahmad, P. Roy, J. C. Beavers, S. Zeiler, J. W. Krakauer, C. Agurto, G. Cecchi, M. Bellard, Y. Raghav, K. Sachs, T. Ehrenberger, E. Bruce, M. E. Cudkowicz, N. Maragakis, R. Norel, J. E. Van Eyk, S. Finkbeiner, J. Berry, D. Sareen, L. M. Thompson, E. Fraenkel, C. N. Svendsen, J. D. Rothstein, Answer ALS, a large-scale resource for sporadic and familial ALS combining clinical and multi-omics data from induced pluripotent cell lines. Nat Neurosci 25, 226–237 (2022).

40. E. Kiskinis, J. Sandoe, L. A. Williams, G. L. Boulting, R. Moccia, B. J. Wainger, S. Han, T. Peng, S. Thams, S. Mikkilineni, C. Mellin, F. T. Merkle, B. N. Davis-Dusenbery, M. Ziller, D. Oakley, J. Ichida, S. Di Costanzo, N. Atwater, M. L. Maeder, M. J. Goodwin, J. Nemesh, R. E. Handsaker, D. Paull, S. Noggle, S. A. McCarroll, J. K. Joung, C. J. Woolf, R. H. Brown, K. Eggan, Pathways disrupted in human ALS motor neurons identified through genetic correction of mutant SOD1. Cell Stem Cell 14, 781–795 (2014).

41. L. Wang, F. Yi, L. Fu, J. Yang, S. Wang, Z. Wang, K. Suzuki, L. Sun, X. Xu, Y. Yu, J. Qiao, J. C. I. Belmonte, Z. Yang, Y. Yuan, J. Qu, G.-H. Liu, CRISPR/Cas9-mediated targeted gene correction in amyotrophic lateral sclerosis patient iPSCs. Protein Cell 8, 365–378 (2017).

42. A. Bhinge, S. C. Namboori, X. Zhang, A. M. J. VanDongen, L. W. Stanton, Genetic Correction of SOD1 Mutant iPSCs Reveals ERK and JNK Activated AP1 as a Driver of Neurodegeneration in Amyotrophic Lateral Sclerosis. Stem Cell Reports 8, 856–869 (2017).

43. K. Kapeli, G. A. Pratt, A. Q. Vu, K. R. Hutt, F. J. Martinez, B. Sundararaman, R. Batra, P. Freese, N. J. Lambert, S. C. Huelga, S. J. Chun, T. Y. Liang, J. Chang, J. P. Donohue, L. Shiue, J. Zhang, H. Zhu, F. Cambi, E. Kasarskis, S. Hoon, M. Ares Jr., C. B. Burge, J. Ravits, F. Rigo, G. W. Yeo, Distinct and shared functions of ALS-associated proteins TDP-43, FUS and TAF15 revealed by multisystem analyses. *Nat Commun* 7, 12143 (2016).

44. R. D. Santis, L. Santini, A. Colantoni, G. Peruzzi, V. de Turris, V. Alfano, I. Bozzoni, A. Rosa, FUS Mutant Human Motoneurons Display Altered Transcriptome and microRNA Pathways with Implications for ALS Pathogenesis. Stem Cell Reports 9, 1450–1462 (2017).

45. S. Hawkins, S. C. Namboori, A. Tariq, C. Blaker, C. Flaxman, N. S. Dey, P. Henley, A. Randall, A. Rosa, L. W. Stanton, A. Bhinge, Upregulation of β-catenin due to loss of miR-139 contributes to motor neuron death in amyotrophic lateral sclerosis. Stem Cell Reports 17, 1650–1665 (2022).

46. A. S. T. Smith, C. Chun, J. Hesson, J. Mathieu, P. N. Valdmanis, D. L. Mack, B.-O. Choi, D.-H. Kim, M. Bothwell, Human Induced Pluripotent Stem Cell-Derived TDP-43 Mutant Neurons Exhibit Consistent Functional Phenotypes Across Multiple Gene Edited Lines Despite Transcriptomic and Splicing Discrepancies. Front. Cell Dev. Biol. 9 (2021).

47. M. Prudencio, V. V. Belzil, R. Batra, C. A. Ross, T. F. Gendron, L. J. Pregent, M. E. Murray, K. K. Overstreet, A. E. Piazza-Johnston, P. Desaro, K. F. Bieniek, M. DeTure, W. C. Lee, S. M. Biendarra, M. D. Davis, M. C. Baker, R. B. Perkerson, M. van Blitterswijk, C. T. Stetler, R. Rademakers, C. D. Link, D. W. Dickson, K. B. Boylan, H. Li, L. Petrucelli, Distinct brain transcriptome profiles in C9orf72-associated and sporadic ALS. Nat Neurosci 18, 1175–1182 (2015).

48. J. P. Ling, O. Pletnikova, J. C. Troncoso, P. C. Wong, TDP-43 repression of nonconserved cryptic exons is compromised in ALS-FTD. Science 349, 650–655 (2015).

49. S. G. Manzo, S. R. Hartono, L. A. Sanz, J. Marinello, S. De Biasi, A. Cossarizza, G. Capranico, F. Chedin, DNA Topoisomerase I differentially modulates R-loops across the human genome. Genome Biol 19, 100 (2018).

50. M. Jangi, C. Fleet, P. Cullen, S. V. Gupta, S. Mekhoubad, E. Chiao, N. Allaire, C. F. Bennett, F. Rigo, A. R. Krainer, J. A. Hurt, J. P. Carulli, J. F. Staropoli, SMN deficiency in severe models of spinal muscular atrophy causes widespread intron retention and DNA damage. Proceedings of the National Academy of Sciences 114, E2347–E2356 (2017).

51. E. A. LaMarca, A. Saito, A. Plaza-Jennings, S. Espeso-Gil, A. Hellmich, M. B. Fernando, B. Javidfar, W. Liao, M. Estill, K. Townsley, A. Florio, J. E. Ethridge, C. Do, B. Tycko, L. Shen, A. Kamiya, N. M. Tsankova, K. J. Brennand, S. Akbarian, R-loop landscapes in the developing human brain are linked to neural differentiation and cell type-specific transcription. Transl Psychiatry 16, 250 (2026).

52. K. Skourti-Stathaki, E. T. Triglia, M. Warburton, P. Voigt, A. Bird, A. Pombo, R-Loops Enhance Polycomb Repression at a Subset of Developmental Regulator Genes. Molecular Cell 73, 930–945.e4 (2019).

53. C. Alecki, V. Chiwara, L. A. Sanz, D. Grau, O. Arias Pérez, E. L. Boulier, K.-J. Armache, F. Chédin, N. J. Francis, RNA-DNA strand exchange by the Drosophila Polycomb complex PRC2. Nat Commun 11, 1781 (2020).

54. J. Brustel, Z. Kozik, N. Gromak, V. Savic, S. M. M. Sweet, Large XPF-dependent deletions following misrepair of a DNA double strand break are prevented by the RNA:DNA helicase Senataxin. Sci Rep 8, 3850 (2018).

55. S. Cohen, N. Puget, Y.-L. Lin, T. Clouaire, M. Aguirrebengoa, V. Rocher, P. Pasero, Y. Canitrot, G. Legube, Senataxin resolves RNA:DNA hybrids forming at DNA double-strand breaks to prevent translocations. Nat Commun 9, 533 (2018).

56. R. Kanagaraj, R. Mitter, T. Kantidakis, M. M. Edwards, A. Benitez, P. Chakravarty, B. Fu, O. Becherel, F. Yang, M. F. Lavin, A. Koren, A. Stewart, S. C. West, Integrated genome and transcriptome analyses reveal the mechanism of genome instability in ataxia with oculomotor apraxia 2. Proc Natl Acad Sci U S A 119, e2114314119 (2022).

57. M. P. Crossley, C. Song, M. J. Bocek, J.-H. Choi, J. Kousorous, A. Sathirachinda, C. Lin, J. R. Brickner, G. Bai, H. Lans, W. Vermeulen, M. Abu-Remaileh, K. A. Cimprich, R-loop-derived cytoplasmic RNA-DNA hybrids activate an immune response. Nature 613, 187–194 (2023).

58. S. Almalki, M. Salama, M. J. Taylor, Z. Ahmed, R. I. Tuxworth, C9orf72-related amyotrophic lateral sclerosis-frontotemporal dementia and links to the DNA damage response: a systematic review. Front Mol Neurosci 18, 1671906 (2025).

59. M. Zimmermann, T. de Lange, 53BP1: pro choice in DNA repair. Trends in Cell Biology 24, 108–117 (2014).

60. E. P. Rogakou, D. R. Pilch, A. H. Orr, V. S. Ivanova, W. M. Bonner, DNA double-stranded breaks induce histone H2AX phosphorylation on serine 139. J Biol Chem 273, 5858–5868 (1998).

61. Y. H. Hsiang, R. Hertzberg, S. Hecht, L. F. Liu, Camptothecin induces protein-linked DNA breaks via mammalian DNA topoisomerase I. J Biol Chem 260, 14873–14878 (1985).

62. Z. Hasanova, V. Klapstova, O. Porrua, R. Stefl, M. Sebesta, Human senataxin is a bona fide R-loop resolving enzyme and transcription termination factor. *Nucleic Acids Res*, gkad092 (2023).

63. C. D. Appel, O. Bermek, V. P. Dandey, M. Wood, E. Viverette, J. G. Williams, J. Bouvette, A. A. Riccio, J. M. Krahn, M. J. Borgnia, R. S. Williams, Sen1 Architecture: RNA-DNA Hybrid Resolution, Autoregulation, and Insights into SETX Inactivation in AOA2. *Mol Cell* 83, 3692-3706.e5 (2023).

64. B. Leonaitė, Z. Han, J. Basquin, F. Bonneau, D. Libri, O. Porrua, E. Conti, Sen1 has unique structural features grafted on the architecture of the Upf1-like helicase family. EMBO J. 36, 1590–1604 (2017).

65. P. Donsbach, D. Klostermeier, Regulation of RNA helicase activity: principles and examples. Biological Chemistry 402, 529–559 (2021).

66. S. Lee, E. J. Huang, Modeling ALS and FTD with iPSC-derived neurons. Brain Res. 1656, 88–97 (2017).

67. M. Galbiati, V. Crippa, P. Rusmini, R. Cristofani, E. Messi, M. Piccolella, B. Tedesco, V. Ferrari, E. Casarotto, M. Chierichetti, A. Poletti, Multiple Roles of Transforming Growth Factor Beta in Amyotrophic Lateral Sclerosis. Int J Mol Sci 21, 4291 (2020).

68. R. Kashima, A. Hata, The role of TGF-β superfamily signaling in neurological disorders. Acta Biochimica et Biophysica Sinica 50, 106–120 (2018).

69. K. Herrup, R. Neve, S. L. Ackerman, A. Copani, Divide and Die: Cell Cycle Events as Triggers of Nerve Cell Death. J Neurosci 24, 9232–9239 (2004).

70. S. C. Namboori, P. Thomas, R. Ames, S. Hawkins, L. O. Garrett, C. R. G. Willis, A. Rosa, L. W. Stanton, A. Bhinge, Single-cell transcriptomics identifies master regulators of neurodegeneration in SOD1 ALS iPSC-derived motor neurons. Stem Cell Reports 16, 3020–3035 (2021).

71. F. Endo, O. Komine, N. Fujimori-Tonou, M. Katsuno, S. Jin, S. Watanabe, G. Sobue, M. Dezawa, T. Wyss-Coray, K. Yamanaka, Astrocyte-derived TGF-β1 accelerates disease progression in ALS mice by interfering with the neuroprotective functions of microglia and T cells. Cell Rep 11, 592–604 (2015).

72. X. Xiong, B. T. James, C. A. Boix, Y. P. Park, K. Galani, M. B. Victor, N. Sun, L. Hou, L.-L. Ho, J. Mantero, A. N. Scannail, V. Dileep, W. Dong, H. Mathys, D. A. Bennett, L.-H. Tsai, M. Kellis, Epigenomic dissection of Alzheimer’s disease pinpoints causal variants and reveals epigenome erosion. Cell 186, 4422–4437.e21 (2023).

73. R. von Bernhardi, F. Cornejo, G. E. Parada, J. Eugenín, Role of TGFβ signaling in the pathogenesis of Alzheimer’s disease. Front Cell Neurosci 9, 426 (2015).

74. D. Gonzalez, X. Cuenca, M. L. Allende, Knockdown of tgfb1a partially improves ALS phenotype in a transient zebrafish model. Front Cell Neurosci 18, 1384085 (2024).

75. D.-Y. Lee, Y. N. Kwon, K. Lee, S. J. Kim, J.-J. Sung, Dual effects of TGF-β inhibitor in ALS - inhibit contracture and neurodegeneration. J Neurochem 168, 2495–2514 (2024).

76. Ö. Yüce, S. C. West, Senataxin, defective in the neurodegenerative disorder ataxia with oculomotor apraxia 2, lies at the interface of transcription and the DNA damage response. Mol. Cell. Biol. 33, 406–417 (2013).

77. A. P. Bracken, N. Dietrich, D. Pasini, K. H. Hansen, K. Helin, Genome-wide mapping of Polycomb target genes unravels their roles in cell fate transitions. Genes Dev. 20, 1123–1136 (2006).

78. J. N. Sleigh, A. M. Rossor, A. D. Fellows, A. P. Tosolini, G. Schiavo, Axonal transport and neurological disease. Nat Rev Neurol 15, 691–703 (2019).

79. CRISPOR: intuitive guide selection for CRISPR/Cas9 genome editing experiments and screens | Nucleic Acids Research | Oxford Academic. https://academic.oup.com/nar/article/46/W1/W242/4995687.

80. C. Lefebvre-Omar, E. Liu, C. Dalle, B. L. d’Incamps, S. Bigou, C. Daube, L. Karpf, M. Davenne, N. Robil, C. Jost Mousseau, S. Blanchard, G. Tournaire, C. Nicaise, F. Salachas, L. Lacomblez, D. Seilhean, C. S. Lobsiger, S. Millecamps, S. Boillée, D. Bohl, Neurofilament accumulations in amyotrophic lateral sclerosis patients’ motor neurons impair axonal initial segment integrity. Cell Mol Life Sci 80, 150 (2023).

81. B. Boulan, A. Beghin, C. Ravanello, J.-C. Deloulme, S. Gory-Fauré, A. Andrieux, J. Brocard, E. Denarier, AutoNeuriteJ: An ImageJ plugin for measurement and classification of neuritic extensions. PLoS One 15, e0234529 (2020).

82. M. Martin, Cutadapt removes adapter sequences from high-throughput sequencing reads. EMBnet.journal 17, 10–12 (2011).

83. D. Kim, G. Pertea, C. Trapnell, H. Pimentel, R. Kelley, S. L. Salzberg, TopHat2: accurate alignment of transcriptomes in the presence of insertions, deletions and gene fusions. Genome Biol 14, R36 (2013).

84. S. Anders, P. T. Pyl, W. Huber, HTSeq--a Python framework to work with high-throughput sequencing data. Bioinformatics 31, 166–169 (2015).

85. S. Anders, W. Huber, Differential expression analysis for sequence count data. Genome Biol 11, R106 (2010).

86. A. Subramanian, P. Tamayo, V. K. Mootha, S. Mukherjee, B. L. Ebert, M. A. Gillette, A. Paulovich, S. L. Pomeroy, T. R. Golub, E. S. Lander, J. P. Mesirov, Gene set enrichment analysis: A knowledge-based approach for interpreting genome-wide expression profiles. Proceedings of the National Academy of Sciences 102, 15545–15550 (2005).

87. B. T. Sherman, M. Hao, J. Qiu, X. Jiao, M. W. Baseler, H. C. Lane, T. Imamichi, W. Chang, DAVID: a web server for functional enrichment analysis and functional annotation of gene lists (2021 update). Nucleic Acids Res 50, W216–W221 (2022).

88. N. Sheth, X. Roca, M. L. Hastings, T. Roeder, A. R. Krainer, R. Sachidanandam, Comprehensive splice-site analysis using comparative genomics. Nucleic Acids Res 34, 3955–3967 (2006).

89. B. Phipson, G. K. Smyth, Permutation P-values should never be zero: calculating exact P-values when permutations are randomly drawn. Stat Appl Genet Mol Biol 9, Article39 (2010).

90. S. Oki, T. Ohta, G. Shioi, H. Hatanaka, O. Ogasawara, Y. Okuda, H. Kawaji, R. Nakaki, J. Sese, C. Meno, ChIP-Atlas: a data-mining suite powered by full integration of public ChIP-seq data. EMBO Rep 19, e46255 (2018).

