## Supplementary figures and tables for "Dysregulated SETX activity at R-loops links transcriptional and splicing alterations to compromised neuronal identity in ALS4"

Marta Giannini *et al.*

**This PDF file includes:**

Figures S1 to S11

Tables S1 and S2

**Other Supplementary Materials for this manuscript include the following:**

Data S1 to S8

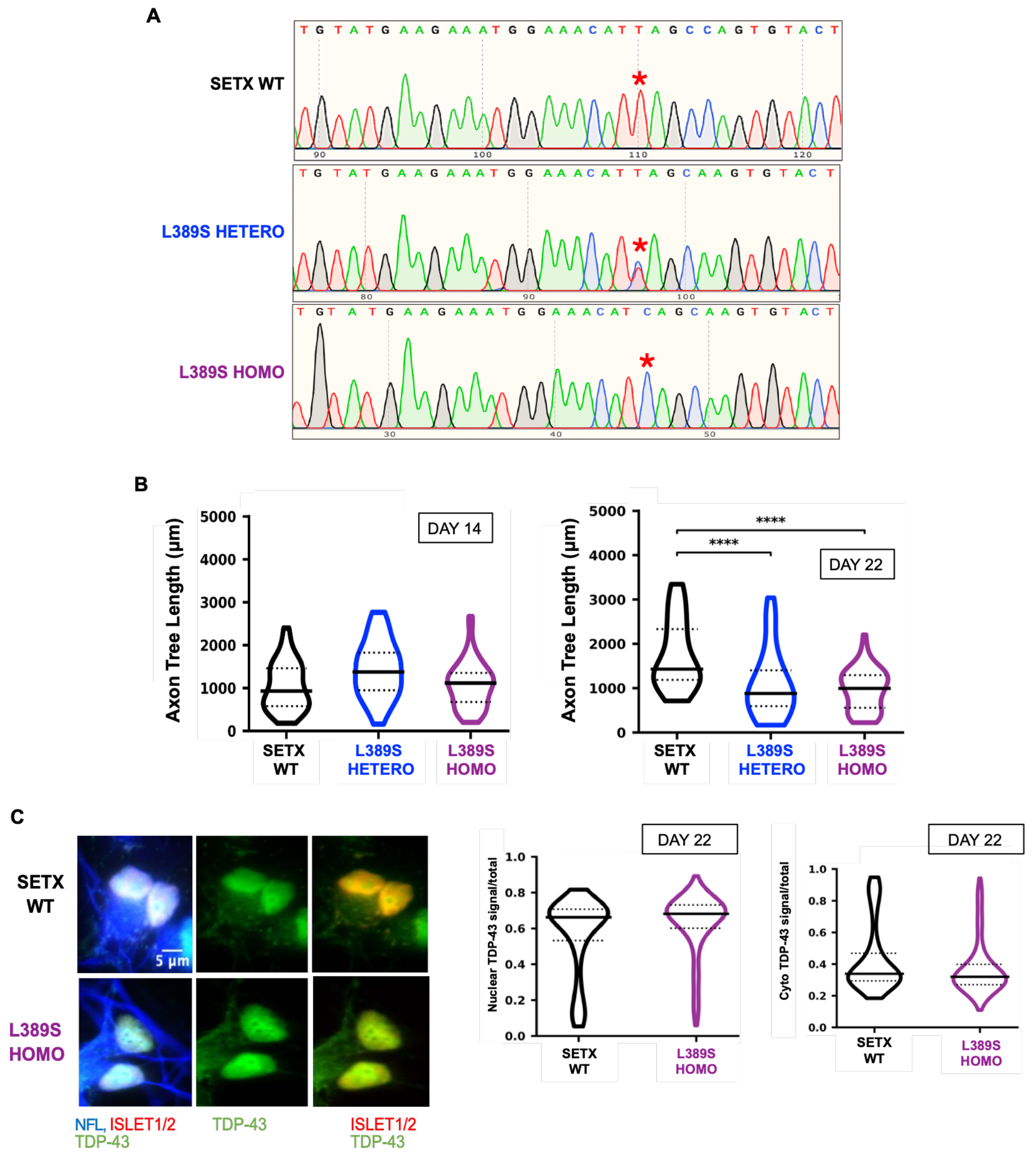


**Figure S1: Validation and characterization of L389S iPSCs-derived MNs**

**A)** Example of electropherograms corresponding to the sequencing of the region of the SETX gene containing the L389S mutation. The red asterisk indicates the position of the nucleotide that is mutated (c.1166T>C) in heterozygous (L389S HETERO, double peak) and homozygous (L389S HOMO, single peak) clones.

**B)** Quantitative analysis of total axon tree length on day 14 and day 22 after iPSC differentiation. The violin plots show the distribution of the data, the median (solid line) and the interquartile range (dotted lines). Data were pooled from 2 independent experiments corresponding to two replicates of one SETX WT clone, two replicates of one HETERO clone and one replicate of each of the two HOMO clones. Neurite tracing was performed using AutoNeuriteJ in Fiji (see methods for details). For each experiment, an average of 30 cells were analyzed per condition. *, $P < 0.05$; **, $P < 0.01$; ***, $P < 0.001$; ****, $P < 0.0001$ (Mann-Whitney U test, two-tailed).

**C)** Analyses of TDP43 cytoplasmic mislocalization and aggregation by immunofluorescence in SETX WT and L389S HOMO MNs at day 22. Left: representative images of one out of two independent experiments. Cells are labelled for TDP-43 (green), ISL1/2 (red, nuclei) and NEFL (blue, soma and neurites). The violin plots (median ± quartiles) represent the quantification of the fraction of TDP-43 fluorescence signal in either the nucleus or the cytoplasm with respect to the total detected signal (i.e. Nuclear TDP-43 signal/total and Cyto TDP-43 signal/total, respectively). Data were pooled from 2 independent experiments performed on one SETX WT and one L389S HOMO clone. For each experiment, in average 150 cells were analyzed per condition.

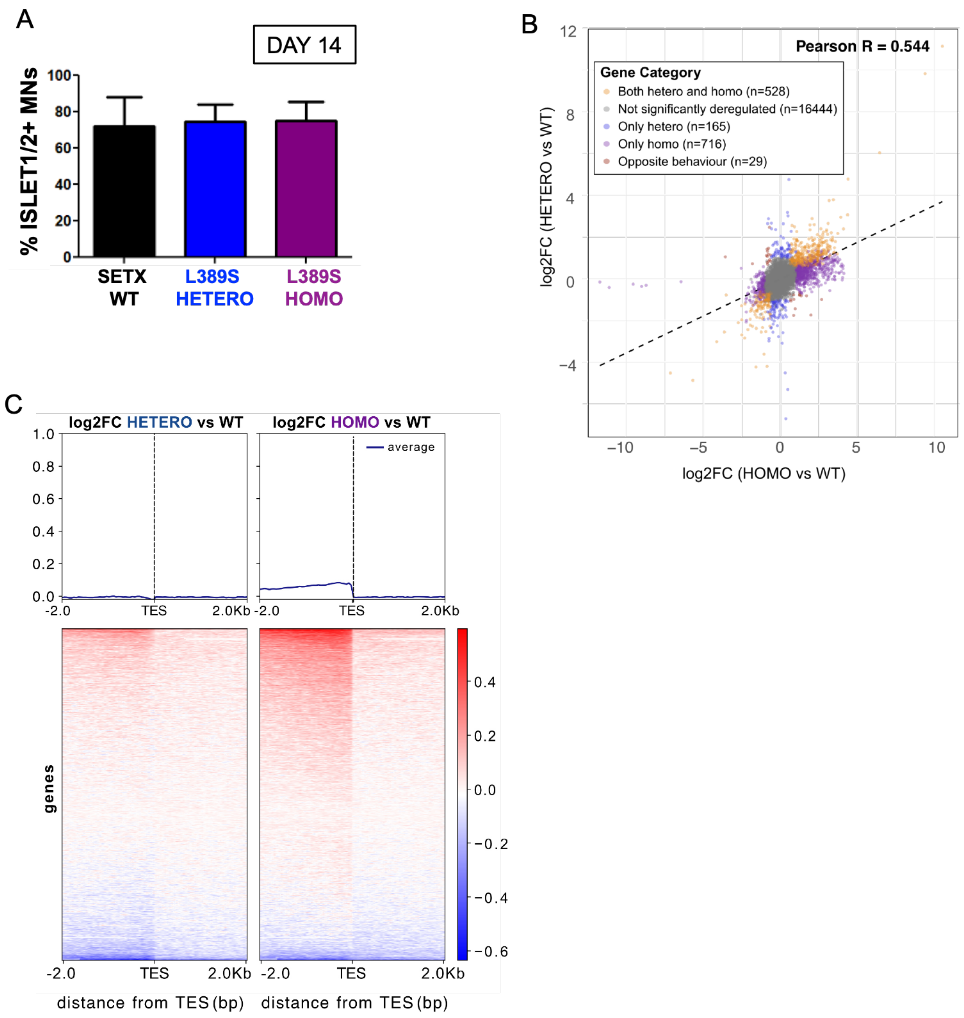


**Figure S2: supplementary data related to figure 2**

**A)** Quantitative analysis of MNs differentiation efficiency is performed in parallel on the same samples used for RNA-seq analyses. Islet1/2 is monitored as MN-specific marker. The histogram shows the percentage of viable cells positive for the MN-specific markers ISL1/2. Data represent the mean and SD from 2 independent experiments with one SETX WT (four replicates in total), one HETERO (two replicates in total) and two HOMO clones (three replicates in total). For each experiment, in average 150 cells were analyzed per condition.

**B)** Scatter plot comparison of DEGs in HETERO and HOMO mutant MNs. Nodes represent individual genes that are coloured according to the category they belong to as indicated in the legend. A dashed line represents a linear regression line. A significant positive correlation was observed between the two mutants with a globally more exacerbated pattern in the homozygous mutants.

**C)** Heatmaps and metagene analyses of possible transcription termination /mRNA 3'end processing defects in L389S HETERO and HOMO mutant MNs. Heatmaps represent the log2 of the fold change (FC) of the RNAseq signal in the mutants relative to the WT around the TES (transcript end site) of protein-coding genes. The summary plot on the top was calculated using the average values for each position.

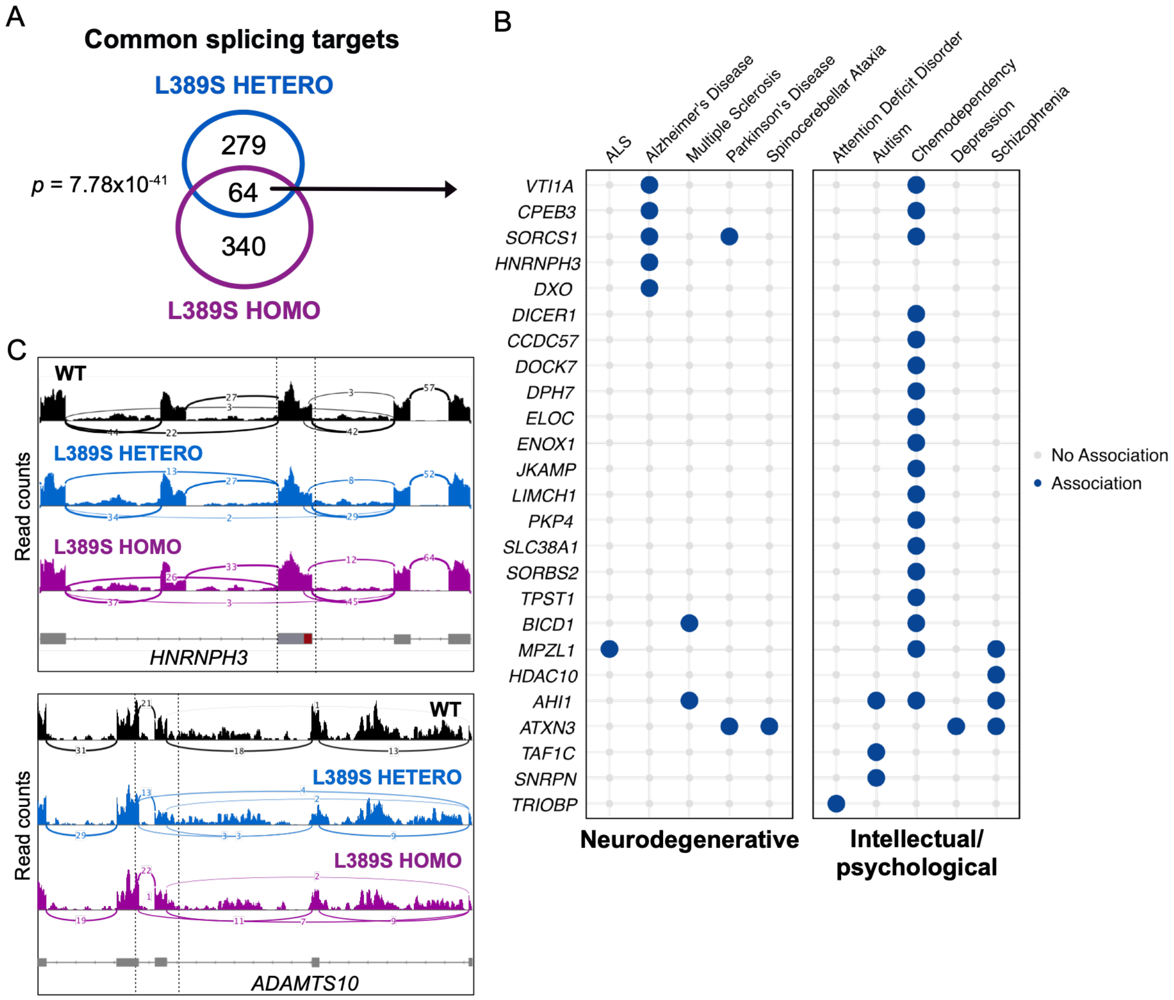


**Figure S3: Functional analyses of the strict set of genes with splicing alterations in both L389S heterozygous and homozygous mutants**

**A**) Venn diagram illustrating the overlap between genes displaying splicing alterations in the HETERO and the HOMO mutants. *p* corresponds to the p-value computed with a hypergeometric test (one-tailed).

**B**) Dot plot indicating the subset of common genes with previously reported association with neurodegenerative diseases and/or intellectual/psychological disorders.

**C**) IGV snapshots of examples of transcripts displaying the same splicing alteration in both types of mutants. Sashimi plots represent RNA-seq read coverage and numbers on the curved lines indicate the count of reads corresponding to exon junctions for each genotype.

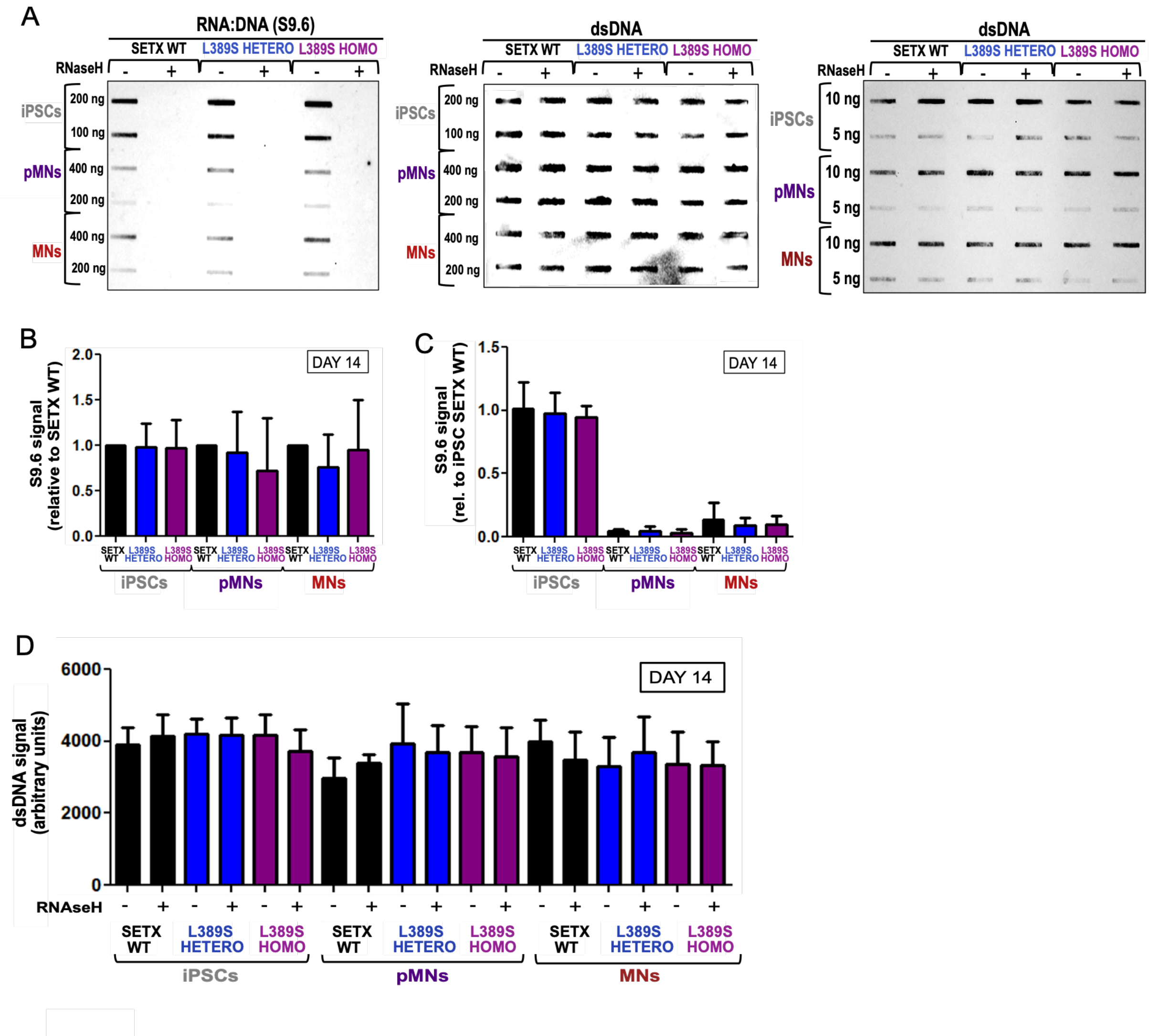


**Figure S4: Global analyses of cellular R-loops in WT and L389S MNs by slot blot.**

**A)** Representative images of global R-loops levels by slot blot using the S9.6 antibody the anti-dsDNA antibody as loading control (one out of three independent biological replicates). Comparison of R-loop levels in iPSC, pMNs and MNs carrying either WT or L389S version of SETX allele in heterozygosis or homozygosis. Samples were treated with (+) or without (-) RNase H to confirm the specificity of the S9.6 signal for RNA:DNA hybrids and with RNAse III to avoid antibody cross-reaction with double-stranded RNA. The blot in the center corresponds to the same blot as in the left but hybridized with the anti-dsDNA antibody. The blot in the right corresponds to the same samples but loading lower concentrations of DNA (10 ng and 5 ng) to demonstrate the linearity of the signal.

**B)** Quantification of S9.6 intensity signal normalized against the respective WT control for each stage of differentiation.

**C)** Quantification of the S9.6 intensity signal normalized against the mean of the SETX WT iPSC clone.

**D)** Quantification of the intensity of the dsDNA signal for the 10 ng samples across all genotypes and treatment conditions (-/+RNase H) in the three stages of differentiation confirming similar loading. For **B-D** data represent the mean and SD from 3 independent experiments with one SETX WT, one HETERO and one HOMO clone. Statistical test (Paired t-test, two-tailed) indicated not significant difference between the R-loop levels in the mutants relative to the WT.

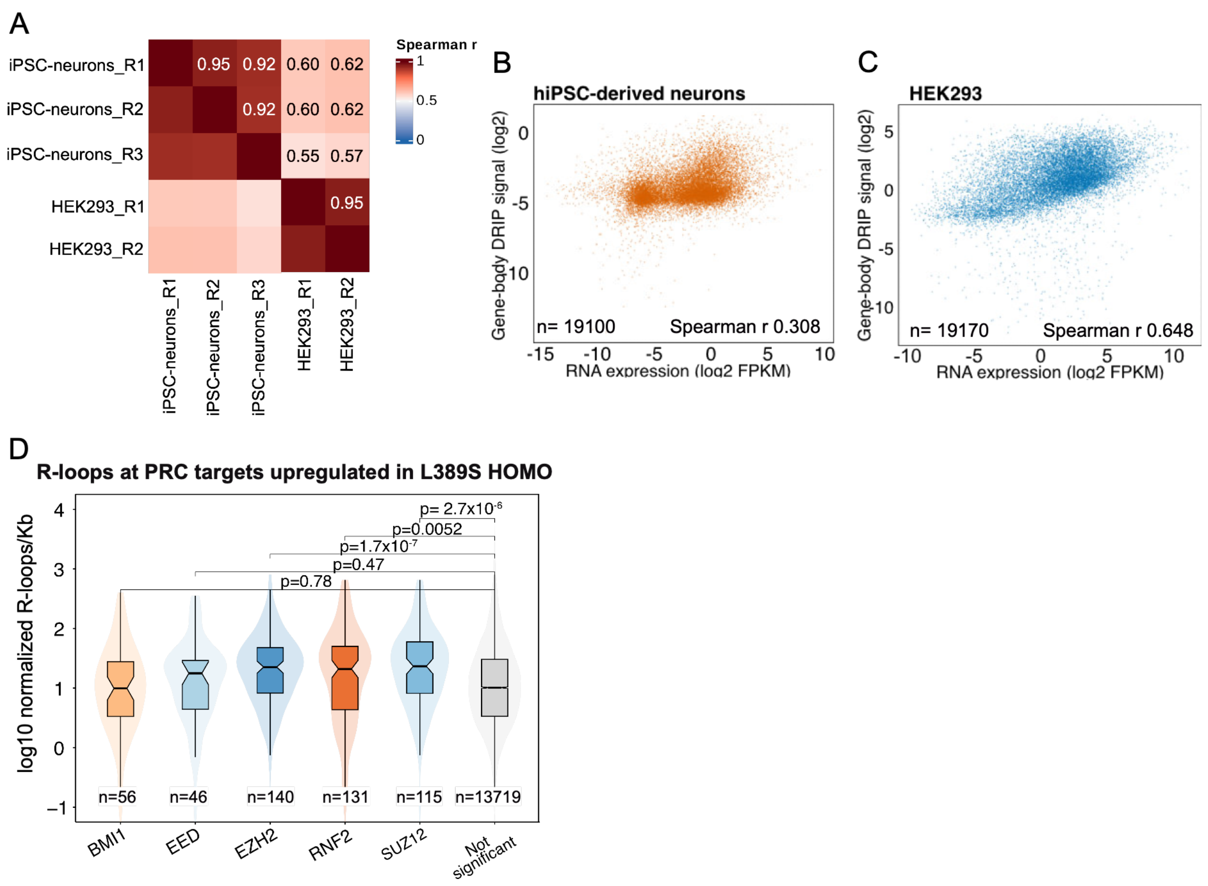


**Figure S5: Quality control and comparison of DRIP-seq datasets generated in iPSC-derived neurons and HEK293 cells.**

**A)** Correlation between biological replicates and datasets generated in iPSC-derived neurons (51) and HEK293 cells (49) used for R-loops analyses in figure 5. Analyses were performed on 10 kb intervals using processed bigwig files.

**B)** and **C)** Correlation between DRIP-seq signal and transcription levels measured by RNA-seq in the same cells and study. Spearman correlations used all mapped genes with finite, nonnegative RNA-seq and DRIP-seq values.

**D)** R-loop propensity at PRC target genes that are upregulated in the L389S HOMO mutants compared to not significantly deregulated genes. Boxplots indicate the median, while the notches represent the 95% confidence interval for the median. Box boundaries represent the first and third quartiles (interquartile range, IQR). Statistical significance was determined using a Wilcoxon rank-sum test (p denotes the p-value).

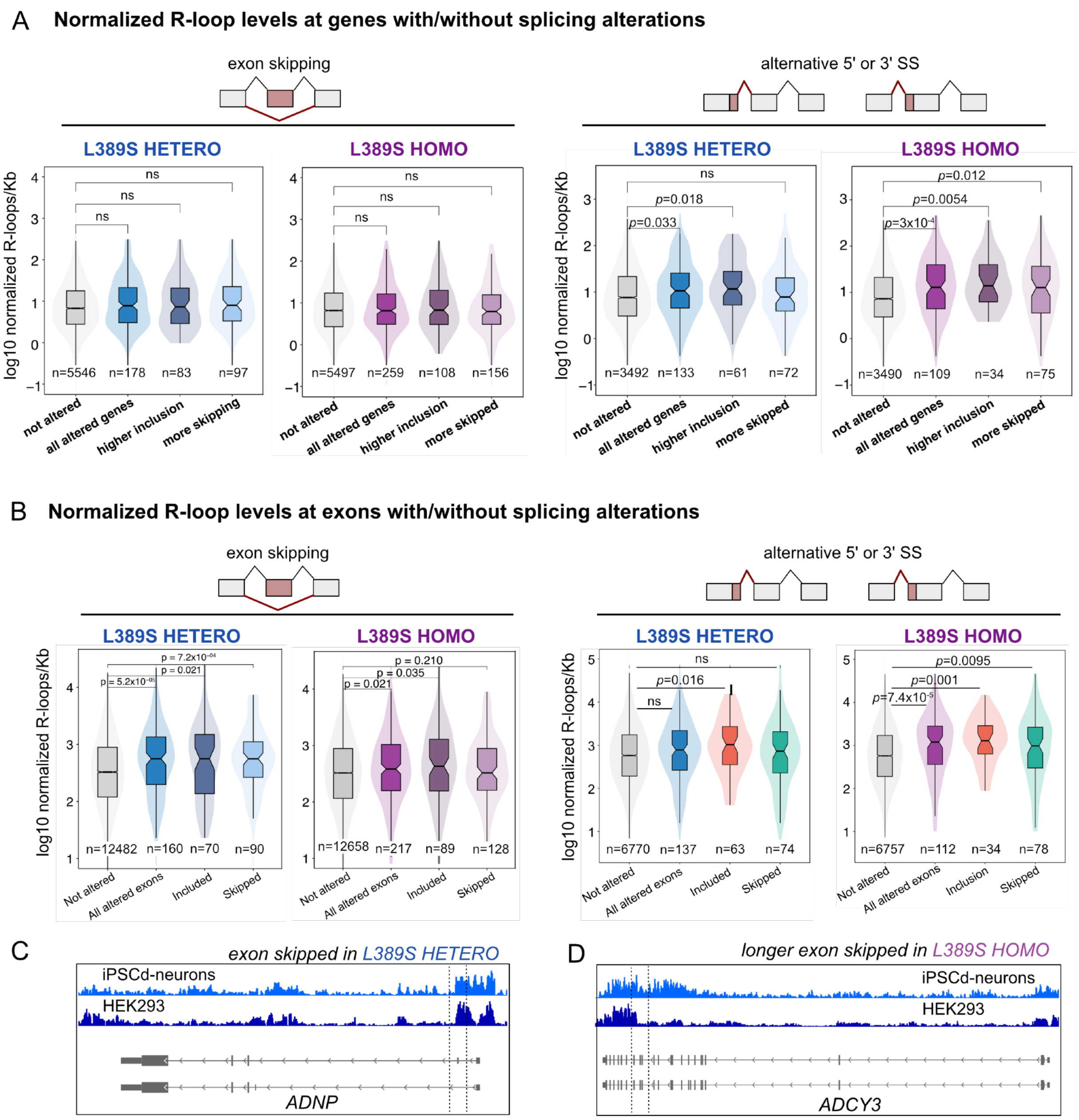


**Figure S6 : Correlation between R-loop propensity and splicing alterations in individual L389S mutants.**

**A)** Boxplots summarizing the R-loop propensity of whole genes carrying exons not exhibiting significant splicing alterations (not altered) or with differential exon skipping (left) or alternative 5’ or 3’ SS usage (right) in L389S mutant MNs. The horizontal line within each box indicates the median, while the notches represent the 95% confidence interval for the median. Box boundaries represent the first and third quartiles (interquartile range, IQR), and whiskers extend to 1.5 times the IQR. Statistical significance was determined using a Wilcoxon rank-sum test (*p* denotes the p-value). **B)** Boxplots representing the distribution of R-loop propensity score for exons not exhibiting significant splicing alterations (not altered) or with differential exon skipping (left) or alternative 5’ or 3’ SS usage (right) in HETERO and HOMO mutants separately. Statistics and formatting of boxplots as in **A**). **C)** and **D)** IGV snapshots of examples of exons displaying higher skipping of the canonical (**C**) or the longer form (**D**) of the exon in ALS4 mutants and with particularly high R-loop levels at flanking regions. Histograms correspond to DRIP-seq datasets generated in the indicated cell lines (iPSCd-neurons denote cell populations enriched in neurons and generated by differentiation of iPSCs). Dashed lines enclose the exon with differential splicing.

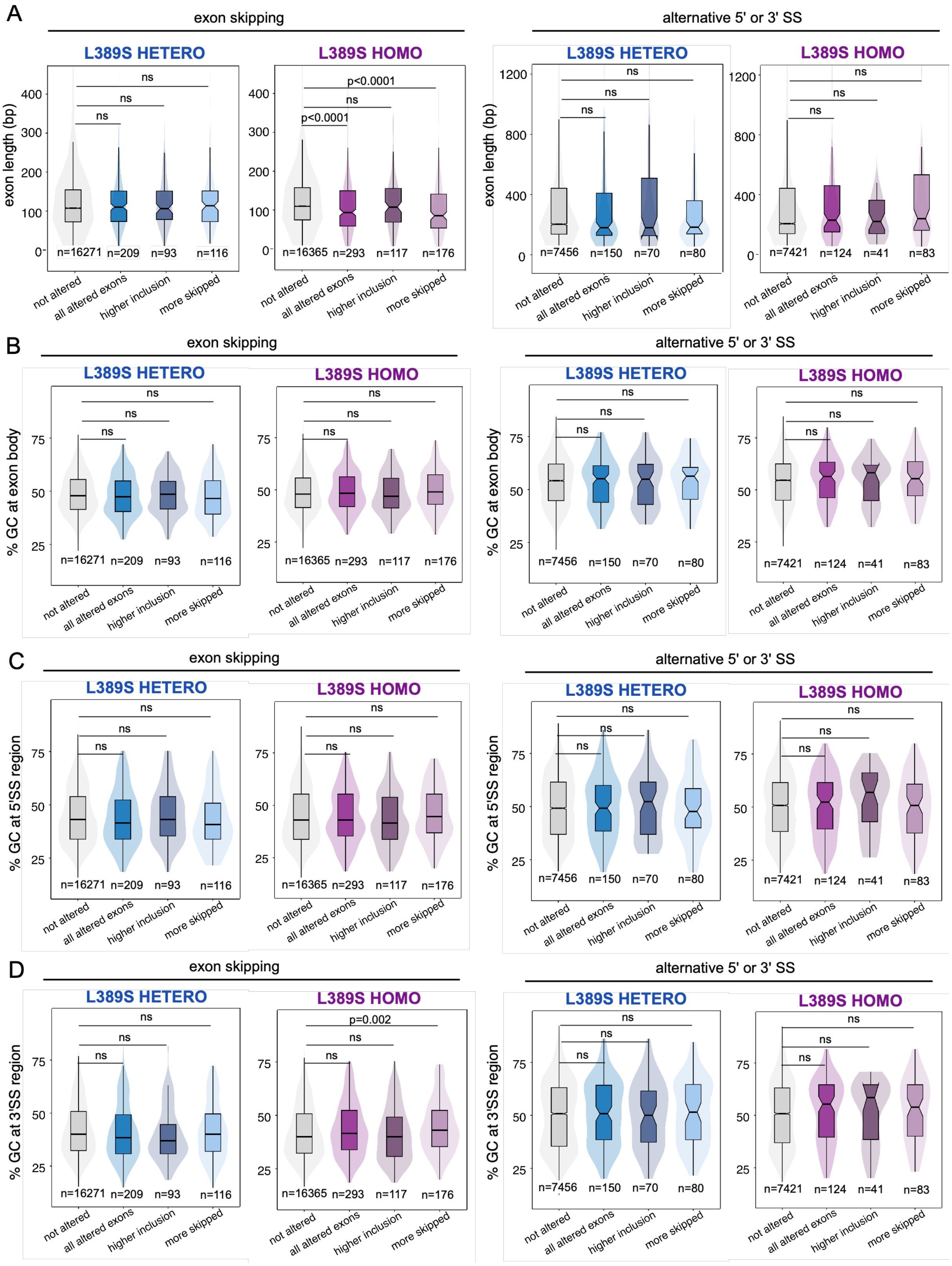


**Figure S7 : Comparison of various splicing-associated metrics between exons differentially spliced in ALS4 MNs.**

**A)** Boxplots representing the length distribution for exons presenting differential skipping or alternative 5’ or 3’ SS usage in L389S mutant MNs compared to the WT. Plots show the median and interquartile ranges. Statistical comparisons between each altered exon subset and the "not altered" reference group were performed using a two-sided Wilcoxon rank-sum test (exact $p$-values or $p < 0.0001$ thresholds are reported on the top, ns= not significant). **B)** Comparison of the GC content of exons differentially spliced in L389S mutants vs WT with respect to non-altered exons. **C)** and **D)** Comparison of the GC content at 5’ SS region (region from -15 bp in exon to +50 bp in intron) or the 3’ SS region (region from -50 bp in intron to +15 bp in exon). Box plots and statistical analyses performed as in **A**).

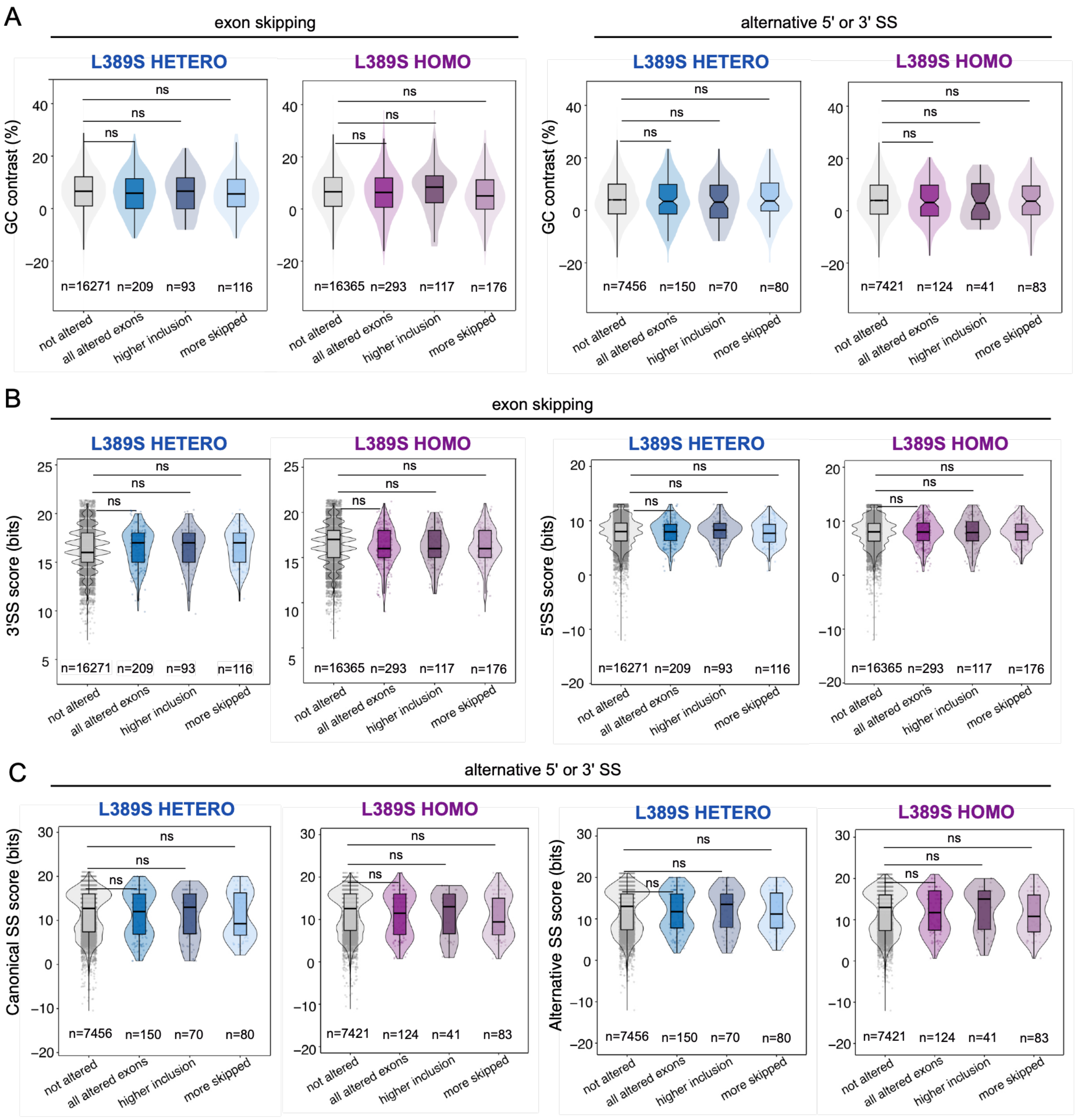


**Figure S8 : Comparison of GC contrast between exons and flanking introns and splice site strength for exons differentially spliced in ALS4 MNs.**

**A)** Boxplots representing the distribution of GC contrast value (difference in GC content between the exon body and 100 bp flanking regions) for exons presenting differential skipping or alternative 5’ or 3’

SS usage in in L389S mutant MNs compared to the WT. Plots show the median and interquartile ranges. Statistical comparisons between each altered exon subset and the "not altered" reference group were performed using a two-sided Wilcoxon rank-sum test.

**B)** and **C)** Boxplots representing the distribution of SS strength values computed for both SS in the case of exons with differential skipping and for the canonical and alternative SS in the case of exons displaying alternative SS usage in mutant MNs. Box plots and statistical analyses performed as in **A**).

A

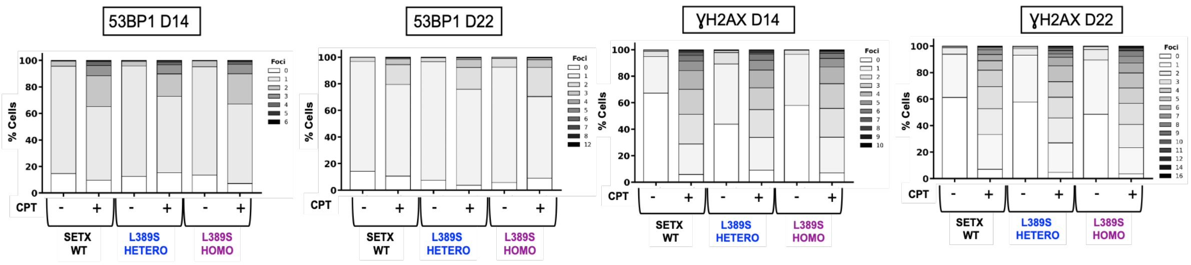


**Figure S9: Quantification of DNA damage foci in WT and mutant MNs**.

Stacked bar charts show the distribution of cells according to the number of nuclear foci for the DNA damage markers 53BP1 and γH2AX at day 14 (D14) and day 22 (D22) of differentiation from iPSCs into MNs in the absence (-) and in the presence of CPT (+) for WT, HETERO and HOMO mutant clones. Data are pooled from three independent experiments. This analysis highlights the shift of the cell population towards classes with a higher damage burden following CPT treatment.

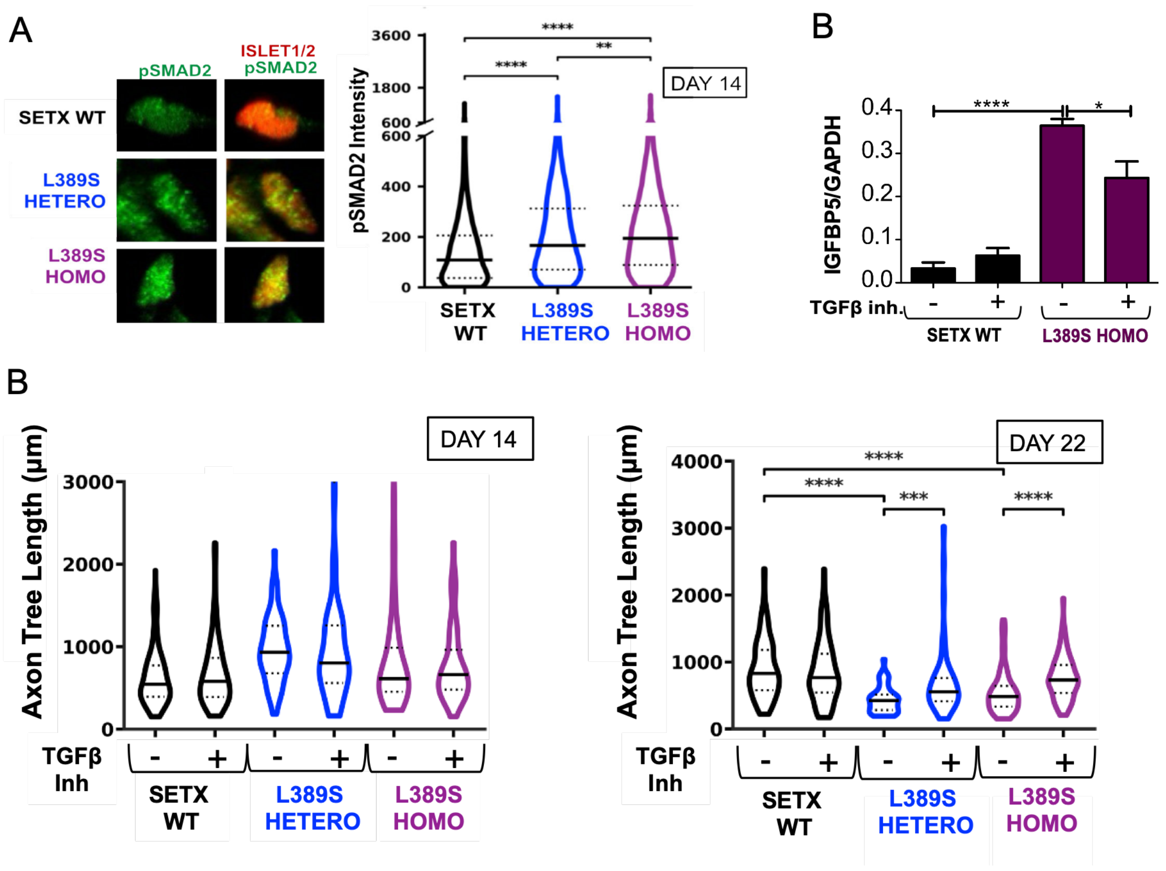


**Figure S10: TGF-β overactivation and effect of its inhibition on MNs axonal tree length.**

**A)** Analysis of TGF-β activation at day 14 by immunofluorescence. **Left:** representative images showing signal of pSMAD2 in SETX WT, HETERO and HOMO mutant MNs that are positive for the nuclear ISLET1/2 marker (red). **Right:** violin plots showing the intensity of nuclear fluorescence for pSMAD2 with median values indicated by solid central black line, and interquartile ranges by dashed lines. Data were pooled from 3 independent experiments performed on one SETX WT, one HETERO and one HOMO clones. For each experiment, an average of 100 cells were analyzed per condition. *, $P < 0.05$; **, $P < 0.01$; ***, $P < 0.001$; ****, $P < 0.0001$ (Mann-Whitney U test, two-tailed).

**B)** Analysis of the effect of treatment with TGF-β and BMP inhibitors on the expression of the IGFBP5 gene, involved in angiogenesis and osteogenesis. Plots represent the values of RNA abundance measured by RT-qPCR and normalized by the housekeeping *GAPDH* gene. Values correspond to the average and SD of three independent biological replicates. *, $P < 0.05$; **, $P < 0.01$; ***, $P < 0.001$; ****, $P < 0.0001$ (unpaired t-test, one-tailed).

**C)** Analysis of the effect of treatment with TGF-β and BMP inhibitors on MNs axonal tree length. Violin plots show the quantification of axonal tree length at days 14 and 22 for the different genotypes, with median values indicated by solid central black line, and interquartile ranges by dashed lines. Data were pooled from 4 independent experiments performed on two SETX WT (six replicates in total), one HETERO (four replicates in total) and one HOMO (four replicates in total) clones. For each experiment, an average of 25 cells were analyzed per condition. *, $P < 0.05$; **, $P < 0.01$; ***, $P < 0.001$; ****, $P < 0.0001$ (Mann-Whitney U test, two-tailed).

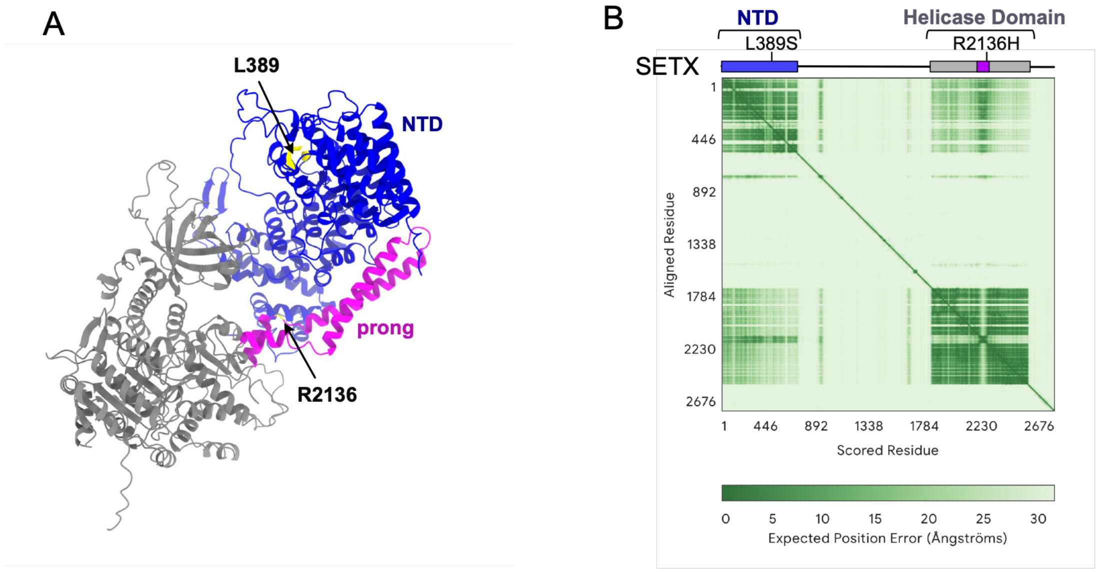


**Figure S11: AlphaFold prediction of SETX suggests a potential intra-molecular interaction between the N-terminal domain and the helicase prong subdomain.**

**A)** Predicted structure of SETX generated with AlphaFold and visualized in UCSF ChimeraX. Regions with low confidence (pLDDT < 70) are hidden for clarity. The N-terminal domain (NTD; residues 1–605) is shown in blue, the helicase domain (HD; residues 1699–2472) in grey, and the prong subdomain (residues 2061–2147), which protrudes from the HD, in magenta. The spatial proximity between the NTD and the prong suggests a potential intramolecular interaction.

**B)** Predicted aligned error (PAE) plot corresponding to the SETX model. A scheme of SETX domains indicating the position of two penetrant ALS4 mutations (L389S and R2136H) is shown on the top. The color scale indicates the expected position error (Å) between residue pairs. The relatively low PAE values observed between the NTD and prong regions support a defined relative positioning of these domains, consistent with a possible interaction.

| Gene | Sequence | Orientation |
|---|---|---|
| SETX | CGACTTGGGTGTTGAACC | Primer FWD |
| SETX | ATTGCTGTTCCTTTGGAGC | Primer REV |
| RGS7BP | TGCCATGCACGAGAATCAGT | Primer FWD |
| RGS7BP | GTCTTTACGAGGCCCTTGGGG | Primer REV |
| ACO16730.1 | AAGGCGTTCGTTATGTGGGA | Primer FWD |
| ACO16730.1 | ACAAGCCATTGCCCTTAGCT | Primer REV |
| AGPAT9 | CAAGTGTGTAGGGGCTGGAG | Primer FWD |
| AGPAT9 | AGGCACTGCAATACCACTCAA | Primer REV |
| RASSF8 | GGTGACTGTAGTTGCATTTGTGT | Primer FWD |
| RASSF8 | AAGGTCCCTGAGAGTGGTCAT | Primer REV |
| SETX | CGACTTGGGTGTTGAACC | Primer FWD |
| SETX | ATTGCTGTTCCTTTGGAGC | Primer REV |
| NEFH | CCAAGGTGAACACAGACGCT | Primer FWD |
| NEFH | CCAGTGAGTCCTTGGTGCTT | Primer REV |
| COL2A1 | GATGAGCAGGGCCTTCTTGA | Primer FWD |
| COL2A1 | GATGACCTTCCTACGCCTGC | Primer REV |
| FN1 | CCAGGCAGTACAATGTGGGT | Primer FWD |
| FN1 | ACGGTGTACTCAGATGCAGG | Primer REV |
| VIM | AACGCCAGATGCGTGAAATG | Primer FWD |
| VIM | ACGAAGGTGACGAGCCATTT | Primer REV |
| IGFBP5 | CGACAAACTTGGACTGGGTC | Primer FWD |
| IGFBP5 | ATCTCCGAGCTGAAGGCTGA | Primer REV |
| GAPDH | ACATCGCTCAGACACCATG | Primer FWD |
| GAPDH | TGTAGTTGAGGTCAATGAAGGG | Primer REV |

**Table S1: Oligonucleotides used in this study.**

Target genes, specific primer sequences (written in the 5′ to 3′ direction), and orientation (FWD: forward; REV: reverse) are indicated.

| Antibody | Provider | Dilution | Experiment |
|---|---|---|---|
| Islet 1/2 (Monoclonal Mouse MIgG2b) | 39.4D5, DSHB | 1:10 | IF |
| Islet-1 (Polyclonal Goat IgG) | AF1837, R&D systems | 1:1000 | IF |
| MNR2/HB9/MNX1 (Monoclonal Mouse MIgG1) | 81.5C10, DSHB | 1:50 | IF |
| NFL (Polyclonal Chicken IgY) | CH22105, Neuromics | 1:500 | IF |
| TDP-43 (Polyclonal Rabbit IgG) | 10782-2-AP, Proteintech | 1:50 | IF |
| Phospho-SMAD1 (Ser463/465)/ SMAD5 (Ser463/465)/ SMAD9 (Ser465/467) (Monoclonal Rabbit IgG) | D5B10, Cell Signaling Technology | 1:200 | IF |
| Phospho-SMAD2 (Ser465/Ser467) (Monoclonal Rabbit IgG) | E8F3R, Cell Signaling Technology | 1:800 | IF |
| 53BP1 (Polyclonal Rabbit IgG) | NB100-304SS, Novus | 1:500 | IF |
| Anti-phospho-Histone H2A.X (Monoclonal Mouse IgG1) | JBW301, Merck | 1:500 | IF |
| Donkey anti-Goat IgG (H+L) Secondary Antibody, Alexa Fluor™ 647 (Polyclonal) | A-21447, ThermoFisher | 1:1000 | IF |
| Donkey anti-Mouse IgG (H+L) Secondary Antibody, Alexa Fluor™ AF555 (Polyclonal) | A-31570, ThermoFisher | 1:1000 | IF |
| Goat anti-Chichen IgY (H+L) Secondary Antibody, Alexa Fluor™ AF647 (Polyclonal) | A-32933, ThermoFisher | 1:1000 | IF |
| Donkey anti-Rabbit IgG (H+L) Secondary Antibody, Alexa Fluor™ AF488 (Polyclonal) | A-21206, ThermoFisher | 1:1000 | IF |
| S9.6 (Mouse) | Hybridoma | 1:6000 | Slot blot |
| dsDNA (Monoclonal Mouse IgG2a) | 35I9, Abcam | 1:1000 | Slot blot |
| m-IgG Fc BP-HRP | sc-525409, Santa Cruz Biotechnology | 1:25000 | Slot blot |

**Table S2: Antibodies used in this study.**

The table reports the target antibody, host clonality/isotype, commercial provider, working dilution, and the specific experimental application.

**Data S1** (separate file): results of differential expression analyses of HETERO and HOMO mutants relative to WT. Genes encoding RNA-binding proteins with a possible role in splicing modulation are highlighted.

**Data S2** (separate file): results of gene ontology analyses of genes differentially expressed in both HETERO and HOMO mutants relative to the WT. BP, MF and CC denote Biological Process, Molecular Function and Cellular Component, respectively.

**Data S3** (separate file): results of GSEA analyses performed on the HETERO mutant using the MSigDB Hallmark gene sets.

**Data S4** (separate file): results of GSEA analyses performed on the HOMO mutant using the MSigDB Hallmark gene sets.

**Data S5** (separate file): results of gene ontology analyses of the 48 genes differentially expressed in all datasets analysed in figure 3B. BP, MF and CC denote Biological Process, Molecular Function and Cellular Component, respectively.

**Data S6** (separate file): results of differential splicing in HETERO and HOMO mutants compared to WT. SE, skipped exons; A5SS, alternative 5' splice site; A3SS, alternative 3' splice site; RI, retained intron; MXE, mutually-exclusive exons.

**Data S7** (separate file): prediction of the outcome of splicing alterations in HETERO and HOMO mutants compared to WT (type of protein/RNA alteration).

**Data S8** (separate file): list of neural PRC targets found among genes upregulated in L389S MNs. Common denotes the genes upregulated in both mutants while HOMO refers to genes upregulated in homozygous mutants.
